# NexuST: A Hierarchical Foundation Model for Spatial Transcriptomics

**DOI:** 10.64898/2026.09.22.753590

**Authors:** Haiping Liu, Qian Zhao, Lijing Lin, Zhiyong Zou, Wenhao Cai, Jingyuan Sun, Yuxi Zhou, Mauricio A. Alvarez, Andrew Gilmore, Magnus Rattray, Alejandro F. Frangi, Hongpeng Zhou

## Abstract

Spatial transcriptomics captures molecular states within cells and their organisation in tissue. However, integrating fine-grained gene information with spatial context at scale remains challenging for existing foundation models. Here we present NexuST, a hierarchical foundation model that repeatedly interleaves gene-level molecular modelling with cell-level spatial modelling, allowing the two levels to refine one another during end-to-end pretraining. For pretraining, we curated HumanST-46M, comprising 45.7 million human cells from 72 datasets across 11 organs and three imaging-based platforms. Across four held-out datasets totalling approximately 2.6 million cells, NexuST achieved state-of-the-art or competitive performance in cell-type annotation, region prediction, gene recovery and neighbourhood-composition prediction. We find that cell-intrinsic expression remains informative even for spatial tasks, as shown by an expression-only PCA baseline, while NexuST shows particularly strong gains where spatial context is essential. Overall, NexuST establishes a hierarchical framework that can serve as a general backbone for future spatial transcriptomics foundation models.

## 1 Introduction

Spatial transcriptomics (ST) measures gene expression while preserving cell locations within tissue, enabling cellular states to be studied in the neighbourhood and microenvironmental contexts that shape tissue organisation and disease. Numerous computational methods have been developed for tasks including cell-type annotation, spatial domain identification, cell–cell communication analysis and niche characterisation [1, 2]. However, most are designed for specific datasets or tasks, limiting their transferability across tissues, platforms and experimental settings while requiring spatial representations to be repeatedly relearned for different downstream applications. Foundation models offer an alternative: by learning general-purpose representations from large, heterogeneous and unlabelled datasets through self-supervised training, they can be adapted to diverse downstream tasks. This strategy has transformed natural language processing [3] and computer vision [4] and has recently been extended to single-cell transcriptomics [5–10].

Foundation models for single-cell RNA sequencing (scRNA-seq) typically represent genes as tokens and use Transformer self-attention to model gene–gene dependencies within individual cells, supporting a wide range of downstream scRNA-seq analysis tasks [5–9]. ST adds a second level of organisation: genes are nested within cells, and cells are arranged across tissue space. This spatial context in principle enables modelling of the cell–cell dependencies that characterise tissue microenvironments. However, modelling cell–cell dependencies across many cells while preserving gene– gene dependencies within each cell creates a fundamental gene–cell scalability tension. Specifically, concatenating gene tokens from many cells into a single sequence preserves detailed gene-level information but produces prohibitively long inputs, whereas compressing each cell into a single representation beforehand makes cross-cell modelling more tractable but discards explicit gene-level information. This trade-off is not merely computational, because gene-level transcriptional states and cell-level microenvironments are themselves biologically coupled. Signals from neighbouring cells can influence a cell’s expression, while a cell’s transcriptional program shapes how it interacts with the surrounding tissue. This interdependence motivates an architecture in which gene-level and cell-level representations can exchange information throughout training, rather than being modelled separately or sequentially. An ST foundation model should therefore retain fine-grained gene-level information while modelling spatial dependencies across large numbers of cells. Designing a computationally tractable architecture that satisfies these requirements remains a central challenge.

Existing ST foundation models span a progression of architectural strategies for addressing this challenge, from directly extending scRNA-seq foundation-model designs to introducing explicit cell-level modelling and, further, to integrating gene- and cell-level representations. Nicheformer [11] and scGPT-spatial [12] largely retain the gene-token architecture developed for scRNA-seq, preserving fine-grained modelling of within-cell gene–gene relationships while adapting this formulation to spatial transcriptomics. Nicheformer primarily learns cell representations from individual-cell expression profiles without explicitly modelling spatial relationships between cells. scGPT-spatial further incorporates neighbourhood information through spatially aware sampling and a neighbourhood-based reconstruction objective during continual pretraining. In both cases, however, cell representations are constructed primarily through within-cell gene-token interactions rather than direct cross-cell contextualisation. A second strategy, adopted by CellPLM [13] and STPath [14], represents each cell using a single token learned from its complete expression profile, which supports scalable cell-level modelling but removes the explicit representation of gene–gene relationships within individual cells. More recently, approaches such as HEIST [15], BrainBeacon [16] and SToFM [17] have moved towards more explicit integration of gene- and cell-level representations, although their cross-level coupling remains constrained. HEIST exchanges information along graphs whose topology is determined from clustering or spatial geometry before training, whereas BrainBeacon and SToFM instead train the two levels sequentially, with gene-level representations already finalised before cell-level modelling begins. Collectively, existing models lack a unified, scalable architecture that jointly optimises gene-level and cell-level representations end-to-end while enabling mutual refinement between the two levels.

Here we present NexuST, a hierarchical Transformer foundation model for spatial transcriptomics that closes this gap. By interleaving gene-level and cell-level Transformer layers within each encoder block, NexuST enables mutual refinement between gene-level and cell-level representations within a single end-to-end model (see Methods Section 4.4). To pretrain NexuST, we assembled HumanST-46M, a corpus of approximately 46 million cells from 72 datasets comprising 123 physical slides across 11 organs and three imaging-based platforms—MERFISH, CosMx and Xenium. HumanST-46M is one of the largest high-resolution human ST corpora assembled to date (Fig. 1a). We evaluated NexuST on held-out benchmarks comprising approximately 2.6 million cells across four downstream tasks under both linear probing and full fine-tuning, including cell-type annotation, tissue-region prediction, held-out gene recovery and neighbourhood-composition prediction. Across datasets and tasks, NexuST achieved state-of-the-art or competitive performance relative to existing foundation models, including Nicheformer [11], scGPT-spatial [12], and CellPLM [13], with particularly consistent gains on tasks that require spatial context. In lung fibrosis tissue, for example, NexuST better distinguished regions with similar coarse cellular compositions but distinct spatial organisations. Its largest gains occurred where cell identity alone was insufficient to determine regional identity, making the surrounding microenvironment essential for accurate prediction. Overall, this work establishes a scalable framework for integrating molecular and spatial information in foundation models for spatial biology, paving the way towards more general and transferable representations of tissue organisation.

**Fig. 1:**
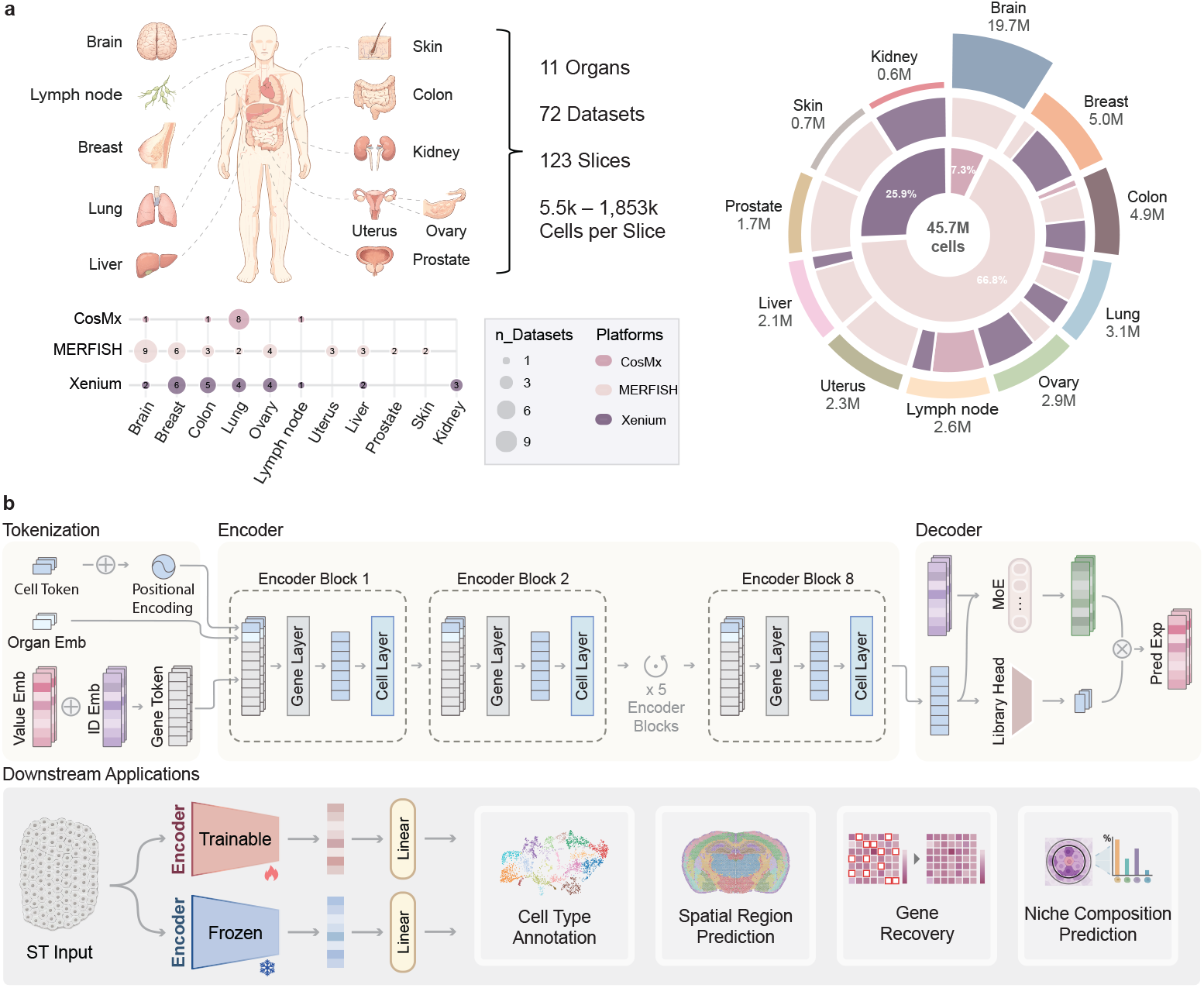
Overview of HumanST-46M and the NexuST framework. **a**, Composition of HumanST-46M, encompassing 45.7 million cells from 72 datasets comprising 123 physical slides, 11 organs and three imaging-based spatial transcriptomics platforms: CosMx, MERFISH and Xenium. **b**, Schematic of NexuST. Gene identities and expression values are tokenised together with cell-level, positional and organ embeddings, processed by hierarchical gene- and cell-interaction layers, and decoded through a factorised decoder with a mixture-of-experts module and library-size head. The learned representations are evaluated across downstream tasks including cell-type annotation, spatial-region prediction, gene recovery and niche-composition prediction.

## 2 Results

### 2.1 A hierarchical Transformer-based foundation model for spatial transcriptomics

#### Overview

NexuST is a Transformer-based foundation model for spatial transcriptomics built around a hierarchical encoder that jointly models gene-level and cell-level dependencies (Fig. 1b). Its novelty lies in representing tissue at the two scales at which it is organised—genes within cells, and cells within a spatial neighbourhood—and coupling these scales inside every encoder block rather than before or after encoding. NexuST was pretrained by self-supervision on HumanST-46M, a large-scale human ST corpus we curated for this work, and we evaluated the resulting representations on four downstream tasks across four held-out benchmarks (Fig. 1b).

HumanST-46M comprises 45.7 million cells from 72 datasets and 123 physical slides, spanning 11 organs and three imaging-based platforms: MERFISH (66.8% of cells), Xenium (25.9%) and CosMx (7.3%). Slides range from 5.5 thousand to 1.85 million cells, and organ coverage is led by brain, followed by breast, colon, lung and seven further organs (Fig. 1a). In curating it, gene names were harmonised across panels so that the same gene maps to the same token in every dataset, while each slide’s panel-specific measured gene set was preserved. We applied neither expression-based filtering nor highly variable gene selection, so that every measured gene remains available to the model through the tokenisation scheme described below. Pretraining jointly across platforms and organs exposes NexuST to broad biological and technical variation for transferable representation learning.

#### Cell and gene representation

Whole-slide training is impractical because a single ST slide can contain thousands to millions of cells. NexuST therefore samples local neighbourhoods of 512 cells from each tissue section and processes each neighbour-hood as one training sample, which preserves local tissue structure while fixing the input size. Within each cell, each measured gene instantiated in the current input is represented by a token obtained by adding a learned gene-identity embedding to a continuous embedding of its log-normalised expression value. Thus, each token encodes both gene identity and expression level. Each cell is represented by up to *M* = 300 valid gene tokens. This choice is supported by HumanST-46M, in which 92.1% of cells contain no more than 300 detected genes, allowing NexuST to effectively retain all detected genes in a single exposure for the large majority of cells while bounding the input length for highly multiplexed panels. For cells with more than 300 detected genes, different subsets are dynamically resampled across exposures (see details in Sections 4.2 and 4.3). Genes that are not measured by a slide’s panel are omitted from the input. In contrast, measured genes with zero expression remain valid sampling candidates and, when selected, enter the model as observed zero-valued inputs. This prevents unmeasured genes from being treated as observed zeros. Each cell’s gene-token sequence is prefixed with a learnable cell token that carries no fixed gene identity and serves as the interface between the gene and cell levels. An organ token is also added to every cell in the neighbourhood to provide tissue-of-origin context.

#### Hierarchical encoder design

The NexuST encoder stacks hierarchical blocks, each consisting of a gene-level Transformer layer followed by a cell-level Transformer layer (Fig. 1b). Within a block, gene-level attention is applied independently within each cell, so that after this layer the cell token summarises the gene-level content of its own cell. Cell-level attention then contextualises the 512 cell tokens across the spatial neighbourhood. Two-dimensional sinusoidal positional embeddings derived from the cell coordinates are applied only at the cell-level because genes within a cell form an unordered set and spatial position is meaningful only at the cell level. The updated spatially informed cell tokens are written back into their own gene-token sequences and therefore participate in gene-level attention in the next block, so that spatial context modulates within-cell gene representations and, in turn, refined gene representations shape the subsequent cross-cell attention. This factorisation is what makes joint modelling tractable: attending over all tokens of a neighbourhood at once would cost quadratically in their total number, whereas separating the two levels reduces this to gene-level attention within cells plus cell-level attention across cells. We pretrained NexuST with 8 hierarchical blocks (16 Transformer layers in total), an embedding dimension of 512 and 8 attention heads, yielding 67.9 million trainable parameters. After the final block, the cell tokens are layer-normalised to yield the cell representations used for all downstream tasks.

#### Pretraining objective and factorised decoder

NexuST is pretrained by masked gene-value reconstruction: for each cell, 20% of the valid gene tokens have their expression-value embedding replaced by a learnable [MASK] embedding while retaining their gene identities, and the model reconstructs the masked values. To account for platform-specific technical variation, the final cell representations are concatenated with a learned platform embedding before decoding, so that batch information is supplied to the decoder rather than absorbed into the encoder representation. The conventional single-output decoder, however, admits a shortcut. Because expression factorises into a cell’s library size and its relative gene composition, the reconstruction loss can be lowered by predicting library size well while falling back on the average composition across cells—collapsing exactly the compositional differences that the representation is meant to preserve (see detailed derivations in Supplementary Note 1). NexuST therefore uses a factorised decoder that predicts composition and library size with separate heads and forms the reconstruction from their product, so that the learned representation is encouraged to preserve biologically informative cell-specific gene distributions rather than sequencing depth. The complete masking procedure, mixture-of-experts decoder and loss are described in Methods Section 4.4. We empirically evaluated these design choices through controlled ablations of the encoder architecture, reconstruction decoder and spatial sampling strategy. The complete design provided the most consistent performance across downstream tasks and datasets (Supplementary Note 4 and Supplementary Tables 13 and 14).

#### Downstream evaluation

We curated four held-out downstream datasets for evaluation: MERFISH human brain, CosMx liver normal, CosMx liver cancer, and Xenium lung fibrosis. None of these datasets was seen during NexuST pretraining. Together, these datasets comprise approximately 2.6 million cells and span three platforms, three organs, healthy and pathological tissue states, and diverse cellular microenvironments (Supplementary Table 2). By adapting and evaluating each pretrained model separately within each downstream dataset, we assessed whether a pretrained representation remained effective across platforms, organs, and tissue states. To avoid task-specific dataset selection, we evaluated each task on all benchmarks for which the required annotations were available. Specifically, cell-type annotation, held-out gene recovery, and neighbourhood-composition prediction were evaluated on all four datasets. Region prediction was the sole exception, as MERFISH human brain does not provide region annotations. NexuST was compared with three foundation-model baselines that represent major design paradigms in the field: Nicheformer, scGPT-spatial and CellPLM, together with PCA as a classical non-pretrained expression-only baseline.

The four downstream tasks span cell-level, gene-level and spatial-context prediction (Fig. 1b). At the cell level, cell-type annotation tests whether the learned representations preserve cell identity. At the gene level, held-out gene expression recovery tests whether genes removed from the input panel can be inferred from the remaining measured genes. For spatial-context prediction, region prediction evaluates whether embeddings capture tissue domains defined by local cellular organisation, while neighbourhood composition prediction directly measures local microenvironmental cell-type composition. Within each benchmark, training and validation partitions were constructed by field of view rather than by random cell sampling. This ensures that neighbouring cells did not appear in both the training and validation sets, thereby reducing spatial data leakage. All methods used the same data partitions and a shared 300-gene candidate panel, with model-specific vocabulary filtering.

Each method was then evaluated under two evaluation protocols, linear probing and full fine-tuning, which both attach a linear head to the encoder output and differ only in whether the pretrained encoder is frozen or jointly trained. Under linear probing, the pretrained encoder is frozen and only the linear head is trained with identical optimisation hyperparameters across methods, thereby ensuring differences reflect the quality of pretrained representation rather than tuning. Under fine-tuning, the encoder and linear head are jointly optimised, measuring task-adapted performance. Optimisation hyperparameters were selected separately for each method under the same tuning budget. PCA, which has no pretrained encoder, was evaluated only under the linear-probing setting, using its principal components as frozen cell representations. Full implementation details for NexuST and all baseline methods are provided in Methods Sections 4.8 and 4.11.

### 2.2 NexuST improves cell-type annotation by resolving spatially contextual cell identities

Cell-type annotation is a fundamental task in spatial transcriptomics that can be used to assess whether the embeddings learned by a foundation model capture biologically meaningful signal. Following the unified evaluation protocol (Methods Section 4.8), we evaluated each model on four held-out datasets (i.e., MERFISH human brain, CosMx liver normal, CosMx liver cancer and Xenium lung fibrosis) under both linear probing and full fine-tuning, training a linear classification head supervised by ground-truth cell-type labels. NexuST achieved the highest macro F1 across all four datasets under both linear probing and fine-tuning, with Nicheformer ranking second (Fig. 2a). The magnitude of this advantage, however, varied substantially across datasets. Under linear probing, all methods performed best on the MERFISH human brain dataset, where NexuST, Nicheformer and PCA were nearly indistinguishable (macro F1 roughly 0.9). In contrast, these methods performed less well on the CosMx liver cancer sample, where NexuST achieved the best performance (0.53), exceeding Nicheformer by 0.10 and PCA by 0.16 in absolute macro F1. Fine-tuning narrowed these performance gaps across platforms and methods, with Nicheformer approaching NexuST in overall macro F1. Nevertheless, NexuST remained the best method on every dataset.

**Fig. 2:**
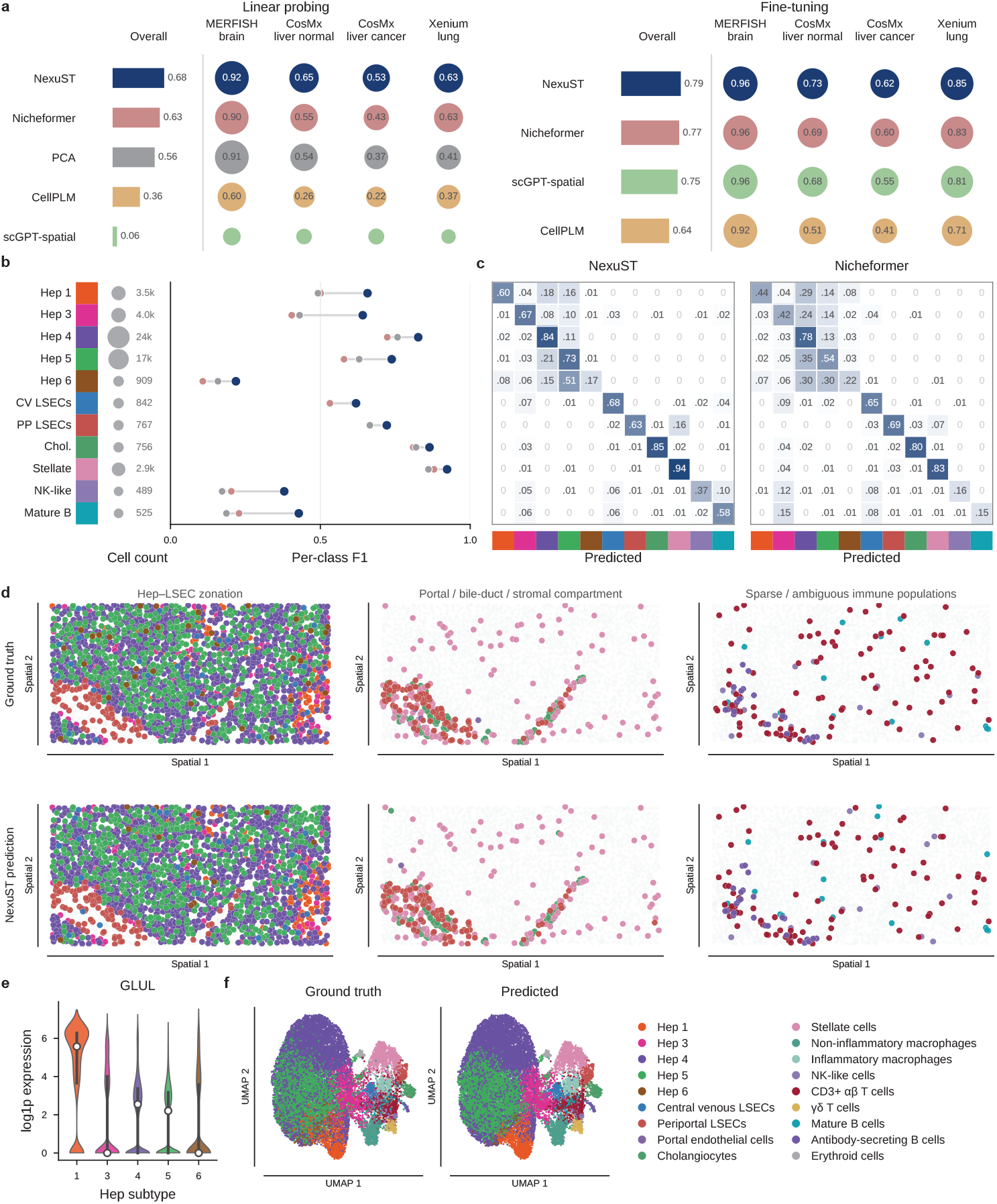
Cell-type annotation performance across spatial transcriptomics datasets. **a**, Macro F1 scores for cell-type annotation under linear probing and fine-tuning across four spatial transcriptomics datasets. Bars show the average performance across datasets, and circles show dataset-specific performance. **b**, Per-cell-type F1 scores for selected cell types in the CosMx liver normal dataset under linear probing. Point positions indicate F1 scores, and circle size indicates the number of cells in each cell type. **c**, Row-normalised confusion matrices comparing NexuST and Nicheformer predictions for the selected cell types in the CosMx liver normal dataset under linear probing. Rows correspond to ground-truth labels and columns correspond to predicted labels.**d**, Spatial distributions of ground-truth and NexuST-predicted labels for selected liver cell populations in the CosMx liver normal dataset under linear probing. Highlighted cell types are shown in colour, and other cells are shown in light grey. **e**, Distribution of GLUL expression across hepatocyte subtypes in the CosMx liver normal dataset, shown as log1p expression. **f**, UMAP visualisation comparing ground-truth annotations and NexuST-predicted labels for annotated cell types in the CosMx liver normal dataset after fine-tuning.

Under linear probing, PCA of log-normalised expression (50 components, Methods 4.11) served as a strong expression-only baseline, achieving competitive performance, particularly in the MERFISH human brain and CosMx liver normal datasets. We therefore treated PCA as a reference for what a linear representation of gene expression alone can achieve and interpreted NexuST’s margin over PCA as a gain attributable to incorporating spatial context. The CosMx liver normal dataset provided the cleanest test of this margin, where PCA performed near the foundation-model baselines (0.01 below Nicheformer) yet remained 0.11 below NexuST. To understand this variation, we examined per-cell-type performance under linear probing (Fig. 2b; Supplementary Fig. 4). NexuST achieved the highest per-class F1 scores across the annotated cell types, with its advantage being greatest for populations poorly resolved by PCA and smallest for populations already near the performance ceiling.

This pattern was most evident at the two ends of the difficulty range. Stellate cells and cholangiocytes exemplified the easy end, being well classified across models. Both achieved F1 scores above 0.8 with PCA alone (Supplementary Fig. 4), consistent with the expectation that these populations are transcriptionally well defined [18]. Cholangiocytes belong to the epithelial lineage, express canonical biliary markers such as KRT19, KRT7, and EPCAM, and typically form well-separated clusters in both single-cell and spatial transcriptomic datasets [19]. Spatially, cholangiocytes are also highly structured, being concentrated around the portal tract and bile duct compartment [20, 21], so both their expression profile and their location are highly stereotyped. Stellate cells, as mesenchymal-derived, perisinusoidal stromal cells, are characterised by a distinct extracellular matrix and mesenchymal gene programme, enabling robust separation even in low-dimensional PCA space. By contrast, mature B cells and NK-like cells were resolved poorly by every method, with PCA F1 scores below 0.2, consistent with their weaker transcriptional separability and more dispersed spatial distributions compared to CD3+*αβ* T cells and antibody secreting B cells. This was not a sample-size effect, since cell abundance was only weakly associated with per-cell-type F1 (Supplementary Fig. 12; *r* ≤ 0.35). These comparisons suggest that by incorporating spatial context, NexuST can improve the annotation of cell populations that are difficult to resolve from gene expression alone.

We compared NexuST with Nicheformer, a leading published spatial foundation model, and found that NexuST showed systematic advantages (Fig. 2a-c). These improvements may reflect its ability to better couple gene- and cell-level representations. This was particularly evident for hepatocyte subtypes. Hepatocytes are spatially zonated along the portal–central axis of the liver lobule, but their transcriptional states form a continuum rather than discrete clusters [18, 22]. This suggests that gene expression alone may be insufficient to separate these hepatocyte subtypes, whereas spatial position within the lobule is directly informative. However, among published foundation models, only Nicheformer achieved PCA-comparable performance, whereas scGPT-spatial and CellPLM performed poorly (Fig. 2a). This may reflect differences in how these models integrate gene expression and spatial information. NexuST explicitly couples gene- and cell-level representations inside every encoder block rather than before or after encoding, which underlies its clear advantage in the classification of hepatocyte subtypes. The row-normalised confusion matrix in Fig. 2c showed that NexuST resolved the most confusable hepatocyte subtypes, such as Hep 4 and Hep 5, more cleanly than Nicheformer. Complete confusion matrices are provided in Supplementary Figs. 7–9. Spatially, Hep 4 showed preferential concordance with periportal LSECs, whereas Hep 1 was more closely associated with central venous LSECs (Fig. 2d). GLUL, a canonical marker of the pericentral zone, varied across subtypes in the same direction (Fig. 2e), indicating that NexuST recovered the metabolic zonation of the lobule rather than an arbitrary partition [18, 22].

Following fine-tuning, all models showed improved performance on the CosMx liver normal dataset, with a noticeable reduction in inter-model differences (Fig. 2a). Nevertheless, NexuST remained the top-performing method, achieving the best results for 17 of 18 cell types (Supplementary Fig. 4). Importantly, it retained a clear advantage over Nicheformer in resolving hepatocyte subtypes (Supplementary Figs. 10 and 11). The corresponding NexuST predictions are visualised in Fig. 2f. NexuST also demonstrated consistent superiority on three other datasets (Supplementary Figs. 3, 5, 6). Together, these results suggest that, by integrating gene expression with cellular neighbourhood information, NexuST more effectively captures coupled gene- and cell-level dependencies, improving annotation performance, particularly for cell types shaped by both transcriptional programmes and microenvironmental context.

### 2.3 NexuST improves region prediction by encoding microenvironment context

A tissue region is an anatomical or histological domain defined by the composition and arrangement of multiple cell populations rather than by the identity of any single cell (Fig. 3a). Region prediction asks to which region each cell belongs. Because the same cell type recurs across distinct regions, a cell’s own expression constrains but cannot fully resolve its region, and closing the gap requires spatial context. We benchmarked region prediction by training a linear classification head with cross-entropy loss to assign each cell to its annotated region, using the three datasets that provide region annotations, i.e., CosMx liver normal, CosMx liver cancer, and Xenium lung fibrosis. The MERFISH human brain dataset provides no region annotation and was therefore excluded from this task. NexuST achieved the highest macro F1 on all three datasets under both linear probing and fine-tuning (Fig. 3b).

**Fig. 3:**
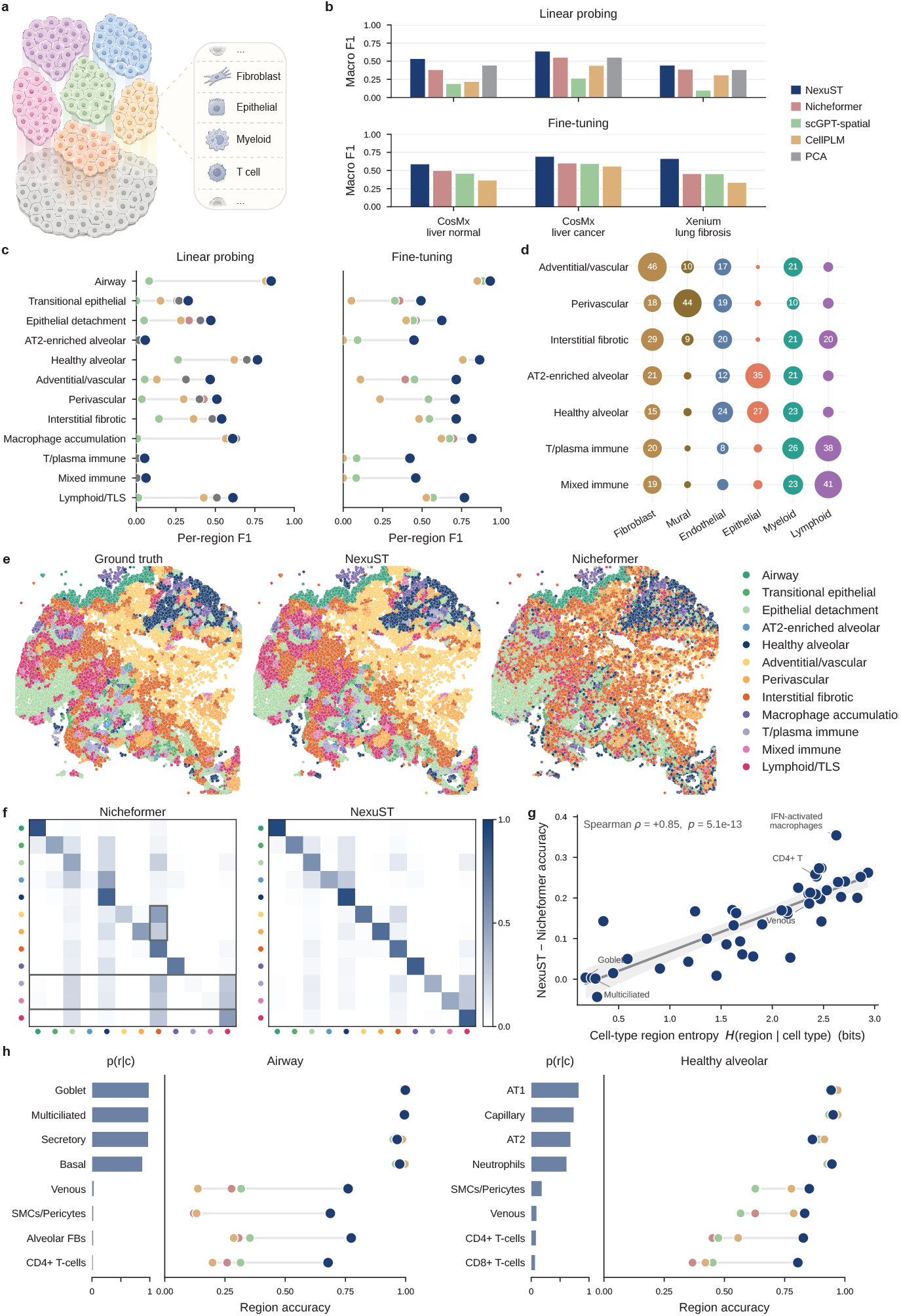
NexuST region prediction across datasets. **a**, Region-level prediction task. Schematic illustration of tissue-region prediction, where annotated regions are defined by local combinations of major cell lineages rather than by isolated cell identities alone.**b**, Overall region-prediction performance. Macro F1 across three spatial transcriptomics datasets under linear probing and full fine-tuning. **c**, Per-region performance on Xenium lung fibrosis. Per-region F1 under linear probing and fine-tuning. **d**, Lineage composition of selected regions. Bubble plot showing the major-lineage composition of selected Xenium lung fibrosis regions. Bubble size denotes the proportion of cells from each major lineage within each region, highlighting that several difficult regions share similar coarse lineage compositions. **e**, Spatial prediction maps. Ground-truth region labels and model predictions for Xenium lung fibrosis. NexuST better preserves local region structure and tissue boundaries, whereas Nicheformer produces more spatially fragmented or smoothed predictions. **f**, Region-level confusion matrices. Confusion matrices comparing Nicheformer and NexuST on Xenium lung fibrosis. Rows and columns correspond to annotated and predicted regions, respectively. **g**, Cell-type dispersion is associated with NexuST gain. Association between cell-type region entropy, *H*(region | cell type), and the accuracy gain of NexuST over Nicheformer. Each point represents a cell type and cell types distributed across more regions show larger NexuST gains. **h**, Representative region-level case studies. Case studies for Airway and Healthy alveolar regions. Left bars show *p*(*r* | *c*), the probability that cells of a given cell type belong to the target region, and right panels show region-prediction accuracy for each model within the same cell types.

As in the annotation benchmark, we used PCA as an expression-only reference under linear probing. NexuST outperformed PCA on every dataset, suggesting that it captures region-level signals beyond gene expression alone, consistent with its hierarchical attention-based encoding of spatial context. The more informative comparison, however, is between PCA and the published foundation models. Although PCA receives no spatial information of any kind, it outperformed all of them except NexuST on both CosMx liver datasets and ranked third on Xenium lung fibrosis, close to second-ranked Nicheformer. Therefore, what matters is not whether a model encodes spatial context, but how it does so. Under fine-tuning, NexuST maintained its lead, with Nicheformer the strongest competing foundation model (Fig. 3b). The advantage is particularly pronounced on Xenium lung fibrosis, where NexuST leads Nicheformer by 0.21 in F1. More notably, the frozen NexuST linear probe already surpassed every fine-tuned foundation model on both liver datasets. NexuST representations are therefore both strong without adaptation and readily adaptable when adaptation is available.

To identify which regions account for NexuST’s overall advantage, we computed per-region F1 on Xenium lung fibrosis for all methods (Fig. 3c). We first used the linear-probing results, together with PCA as an expression-only baseline, to stratify regions by their dependence on gene expression versus spatial context (Fig. 3c and d). For example, regions such as Airway and Healthy alveolar were well captured by PCA, both being defined by transcriptionally distinctive cell types. At the opposite extreme, AT2-enriched alveolar, T/plasma immune and Mixed immune were poorly resolved. NexuST gained most in the intermediate regions where gene expression provided partial but insufficient regional information, such as Adventitial/vascular and Perivascular (ΔF1 = 0.16 and 0.11 relative to PCA).

After fine-tuning, NexuST achieved the best performance across all regions and showed larger improvements than Nicheformer. These fine-tuning gains were most informative in two region categories. The first comprised regions poorly resolved from gene expression alone, including AT2-enriched alveolar, T/plasma immune and Mixed immune. These regions were not defined by a single transcriptionally distinct cell type, but instead reflected regional states shaped by local tissue composition and microenvironmental context (Fig. 3d). For example, AT2-enriched alveolar is a region enriched for AT2 cells. However, the same AT2 cells recur in Healthy alveolar, Epithelial detachment and Transitional epithelial regions. What marks an AT2 cell as belonging to AT2-enriched alveolar is therefore not its own expression profile. Similarly, T/plasma immune and Mixed immune regions are likely influenced by broad immune-cell infiltration, such that immune cells with similar transcriptional profiles may be distributed across these regions. As a result, expression-only information is insufficient to accurately assign these cells to their regional context. By incorporating spatial information, NexuST can leverage neighbourhood structure and local tissue organisation, leading to substantially improved recovery of these regions.

This interpretation is supported by comparison with Nicheformer (Fig. 3e,f): both the predicted spatial distribution and the confusion matrix show that Nicheformer frequently assigned AT2-enriched alveolar cells to Healthy alveolar, Epithelial detachment and Transitional epithelial regions, whereas NexuST recovered the AT2-enriched alveolar region far more accurately. The T/plasma immune and Mixed immune regions behave in a similar way. Nicheformer confused these immune regions with other inflammation-associated regions, including Epithelial detachment, Interstitial fibrotic and Lymphoid/TLS. By contrast, NexuST substantially reduced this confusion by leveraging spatial context, particularly by separating Interstitial fibrotic regions from neighbouring immune-dominated regions. This led to marked improvements in the classification of both immune regions and facilitated more accurate identification of important immune structures, including TLS.

The second comprised intermediate regions, such as Adventitial/vascular and Perivascular, for which NexuST already showed an advantage in the linear-probing analysis. Unlike the poorly resolved regions, where Nicheformer almost failed to recover the regional labels, intermediate regions retained partial signal from gene expression and could be partially identified. Their main limitation was substantial confusion with Interstitial fibrotic (Fig. 3d-f). For example, the cellular composition of Adventitial/vascular closely resembles Interstitial fibrotic, both containing abundant fibroblasts alongside substantial endothelial and myeloid populations (Fig. 3d; Supplementary Fig. 14a). This compositional similarity likely contributes to the frequent confusion between these two regions. Indeed, the Nicheformer confusion matrix assigns Adventitial/vascular to Interstitial fibrotic more often than to its own label (Fig. 3f). The high abundance of fibroblasts and endothelial cells also led to occasional misclassification into Healthy alveolar regions. By incorporating the surrounding microenvironment rather than composition alone, NexuST substantially reduced this composition-driven ambiguity and more accurately distinguished Adventitial/vascular from neighbouring fibrotic regions (F1 0.27 for Nicheformer versus 0.70 for NexuST). Region-level scores average over all the cell types present, some of which are far easier to assign than others. We therefore stratified by cell type. We reasoned that cell types distributed across many regions would be intrinsically harder to assign to a specific region than those cell types confined to a few regions. To quantify this regional ambiguity, we measured Shannon entropy of each cell type’s region distribution *H*(region | *c*), computed from the ground-truth annotations alone and therefore independent of any model (Methods 4.9.2). We found that the accuracy gain of NexuST over Nicheformer was strongly correlated with this entropy (Spearman *ρ* = 0.85; Fig. 3g), with the largest gains observed for cell types spread across many regions and negligible gains for region-specific cell types. This suggests that NexuST provides the greatest benefit when cell identity alone is insufficient to determine regional identity, consistent with an important contribution from spatial context.

Furthermore, we analysed performance differences between cell types within an individual region. For each cell type, we quantified its regional specificity as the proportion of cells annotated to the region of interest *p*(*r*| *c*). Consistent with the entropy analysis above, highly region-specific cell types showed only minor differences between methods, whereas broadly distributed cell types exhibited much larger performance gaps. For example, in the Airway region, Goblet and multiciliated cells were highly specific to the region, resulting in nearly identical accuracy across methods. By contrast, Venous and CD4+T cells occurred across multiple regions, making regional assignment substantially more difficult. In these cases, NexuST showed a markedly larger advantage (Fig. 3h; Supplementary Fig. 14c).

These observations were further supported by matched ablation experiments: replacing cell-level attention with additional gene-level layers reduced regionprediction performance across all three datasets (Supplementary Table 13b). Together, these analyses show that NexuST’s advantage comes not merely from recognising cell identity, but from encoding the surrounding microenvironment, allowing it to resolve tissue regions precisely where cell identity alone is insufficient.

### 2.4 NexuST supports held-out gene expression recovery under partial molecular observation

Imaging-based spatial transcriptomics assays profile predefined gene panels of a few hundred to roughly one thousand genes, with panel composition fixed at experimental design. This creates a practical limitation: biologically relevant genes may be unmeasured, and each assay provides only a partial view of the underlying molecular state. Gene expression within a panel reflects both gene co-regulation and the local tissue microenvironment. We therefore hypothesised that there are informative dependencies between genes, and that these dependencies can be recovered from gene expression and spatial context. We tested this by withholding a subset of the panel and evaluated whether NexuST could infer the expression of these held-out genes from the remaining observed genes and their spatial context.

In this controlled setting, we defined a vocabulary-compatible HVG panel across models by intersecting each dataset’s HVG panel with the pretrained vocabularies of all four foundation models. Within each dataset, a random 20% of the common panel was designated as held-out target genes and removed from the input before tokenisation; each cell representation was therefore built from the remaining 80% alone. The same split was applied to every model. The withheld genes were then regressed from that representation using a linear head trained with MSE loss. Notably, because the target genes are withheld entirely from the input and all models are evaluated with the same linear readout (Fig. 4a), this protocol places all foundation models on the same footing, avoiding confounding from model-specific choices in masked-gene encoding, zero-value treatment, tokenisation and decoder design. What it measures is therefore how much target-gene expression is recoverable from the representation formed by the observed panel.

**Fig. 4:**
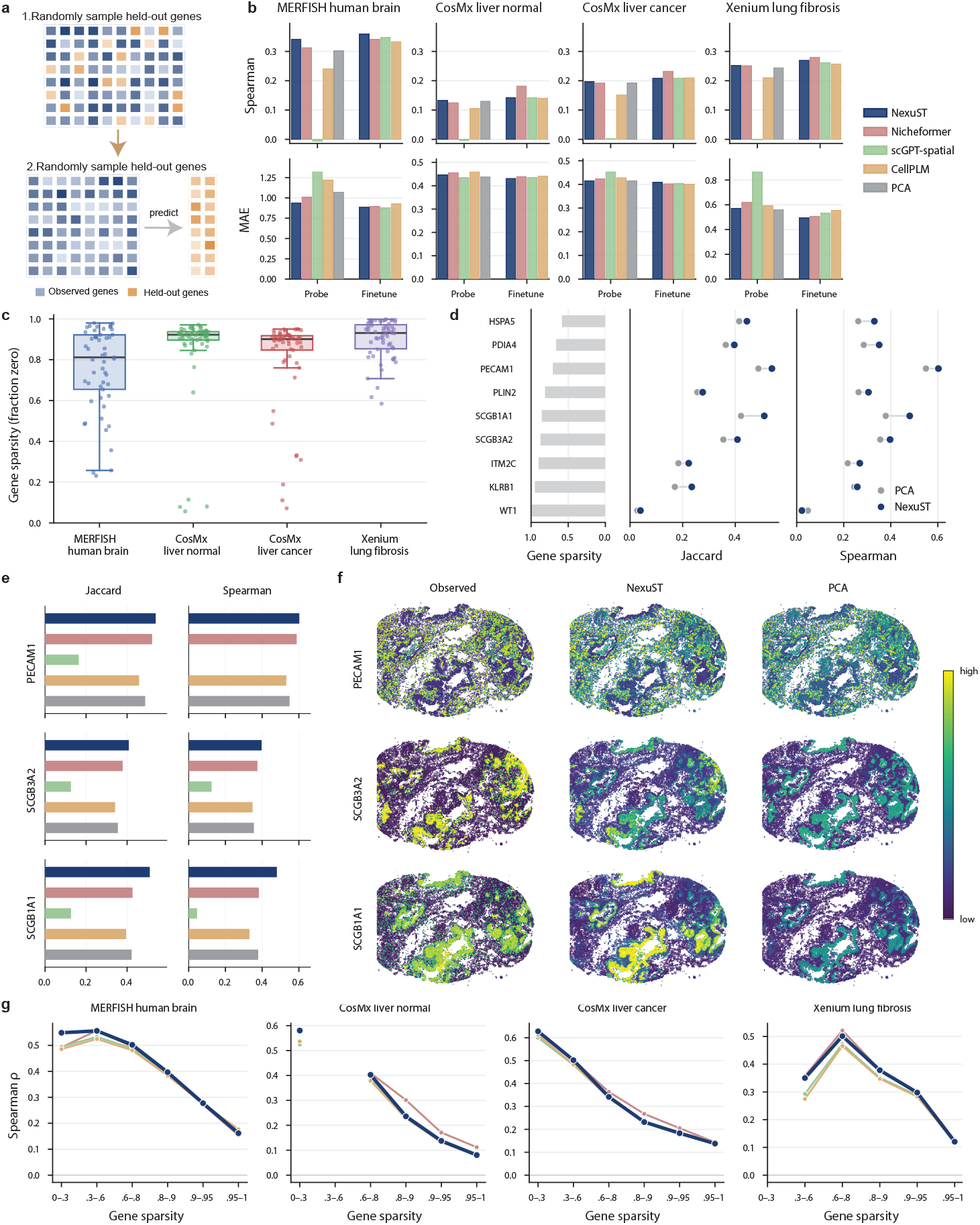
NexuST supports held-out gene recovery across spatial transcriptomic datasets. **a**, Schematic representation of the held-out gene recovery task. A subset of genes is randomly withheld from each cell, and the model is evaluated by predicting the withheld gene expression from the remaining observed genes. **b**, Spearman correlation and mean absolute error (MAE) of held-out gene recovery across MER-FISH human brain, CosMx liver normal, CosMx liver cancer and Xenium lung fibrosis datasets under linear probing and fine-tuning settings. **c**, Distribution of gene sparsity, defined as the fraction of cells with zero expression for each gene, across the four datasets. Box plots show the distribution of gene-level sparsity values, with individual genes shown as points. **d**, Gene sparsity and recovery performance of selected representative genes, comparing NexuST and PCA using Jaccard similarity and Spearman correlation. **e**, Method comparison for three representative genes, showing Jaccard similarity and Spearman correlation across NexuST, Nicheformer, scGPT-spatial, CellPLM and PCA. **f**, Spatial expression maps in the Xenium lung fibrosis dataset comparing observed expression with predictions from NexuST and PCA for PECAM1, SCGB3A2 and SCGB1A1. **g**, Spearman correlation stratified by gene sparsity bins across the four datasets and methods.

We benchmarked held-out gene expression recovery across four datasets, using Spearman correlation and mean absolute error (MAE) as evaluation metrics (Fig. 4b). Under the linear-probing setting, NexuST achieved the lowest median MAE and the highest median Spearman correlation, outperforming Nicheformer, scGPT-spatial, CellPLM, and PCA. As in the preceding tasks, PCA marks what gene expression alone can achieve, and no other method with the exception of NexuST exceeded it on all four datasets. For example, on both liver datasets, PCA ranked second only to NexuST; and on the other two datasets, it remains highly competitive, ranking just behind NexuST and Nicheformer. Motivated by these observations, we further investigated how NexuST’s advantage over the expression-only PCA baseline varies across genes and whether this advantage is associated with spatially organised expression patterns.

We compared NexuST with PCA at the individual gene level, using two metrics that separate what is being recovered (Fig. 4d and Supplementary Figs. 15, 16, 17, 18). Spearman correlation is sensitive both to which cells express a gene and to how much they express, whereas the Jaccard similarity reports only recovery of the expressing-cell pattern (Methods 4.9.3). A gain in Jaccard indicates better recovery of spatial pattern; a gain in Spearman without a matching gain in Jaccard indicates better estimation of expression magnitude. As shown in Fig. 4d, PCA and our model exhibit similar trends as sparsity decreases, and NexuST exceeded PCA for most genes. For the sparsest genes (sparsity *>* 0.9), the two methods converged and both performed poorly under either metric (Spearman typically below 0.2). These genes carry little co-regulation signal within the panel and, due to their sparsity, are less amenable to improvement through spatial context. For broadly expressed genes, the two methods achieved highly comparable performance in Jaccard similarity, but their Spearman correlation differs substantially. This suggests that the advantage of NexuST for these genes lies in expression magnitude estimation rather than expression pattern. Additionally, a subset of genes in an intermediate regime exhibited marked differences in Jaccard similarity, indicating the recovery of spatial pattern contributed more.

Fig. 4d–f illustrate representative genes for these scenarios. SCGB1A1 and SCGB3A2 both exhibit substantial performance improvements in both metrics. But for SCGB1A1, the gain in Spearman correlation is markedly more pronounced than the Jaccard gain. SCGB1A1 encodes a club-cell secretoglobin and a major airway lining-fluid protein, and shows spatially organised expression associated with airway epithelial context, making its recovery particularly amenable to representations that encode tissue organisation. In contrast, SCGB3A2 expression depends more on cell-intrinsic regulatory states, which may explain why the gain was more apparent in spatial pattern recovery.

Fine-tuning improved every model and changed the ranking. Nicheformer outperformed NexuST by a clearer margin on both CosMx liver datasets, while showing a marginal advantage on Xenium lung fibrosis. We therefore investigated what distinguishes those datasets. We assessed the sparsity distribution of the held-out genes in each dataset (Fig. 4c). In all four datasets, the randomly selected genes were predominantly sparse, with varying degrees of sparsity. Specifically, sparsity was near-uniform in the brain dataset; the two liver datasets spanned a broad range, with sparse genes concentrated at the high end; and the lung dataset contained no broadly expressed genes and was uniform within a narrow range. We then stratified genes by sparsity and found that recovery performance decreased as sparsity increased for all models. NexuST and Nicheformer were broadly comparable across most sparsity bins and datasets. The advantage of Nicheformer was clearest on CosMx liver normal in the intermediate-to high-sparsity bins, smaller on CosMx liver cancer, and not apparent on the MERFISH human brain and Xenium lung fibrosis datasets (Fig. 4g). This dataset-dependent pattern suggests that Nicheformer’s fine-tuning advantage may be associated with the distribution of target-gene sparsity and most apparent when the target genes span a broad sparsity range, as in the two CosMx liver datasets. In such settings, Nicheformer may benefit from gene co-expression priors inherited from its larger scRNA-seq pretraining corpus.

Overall, in the linear probing setting, NexuST achieved the best held-out gene recovery on all datasets, with its clearest gains for genes of lower sparsity, whereas Nicheformer required fine-tuning to overtake it on the liver datasets. This is consistent with NexuST already encoding the recoverable signal in its pretrained representation, and with Nicheformer holding co-expression priors that become accessible only once its encoder is adapted. Highly sparse genes remained difficult for every model.

### 2.5 NexuST recovers neighbourhood compositions across diverse spatial transcriptomics datasets

Neighbourhood composition, defined as the relative abundance of cell types within a fixed spatial radius of each cell [11], quantifies how cells co-localise into local tissue microenvironments that shape local signalling and cell behaviour (Fig. 5a). Cells of the same type can reside in distinct local microenvironments, so neighbourhood composition is difficult to infer from cell identity alone. Spatial transcriptomics at single-cell resolution makes this composition directly observable. Because the neighbours that define the target lie within the model’s spatial receptive field, reading composition out of a pretrained per-cell representation is a direct test of how much local spatial context that representation retains. Because microenvironmental structure is scaledependent, we evaluated four dataset-specific radii chosen to yield approximately 10, 20, 50, and 100 neighbours per cell (Methods Section 4.9.4; Supplementary Table 5), each treated as a separate prediction task. The actual number of neighbours remained cell-specific because neighbourhoods were defined by fixed spatial radii rather than *k*-nearest neighbours. We converted neighbour cell-type counts into proportions by dividing each count by the total number of neighbours, rather than applying the softmax transformation used by Nicheformer [11]. This preserves the relative abundance of minority cell types, which would otherwise be compressed by softmax (see Methods Section 4.9.4).

**Fig. 5:**
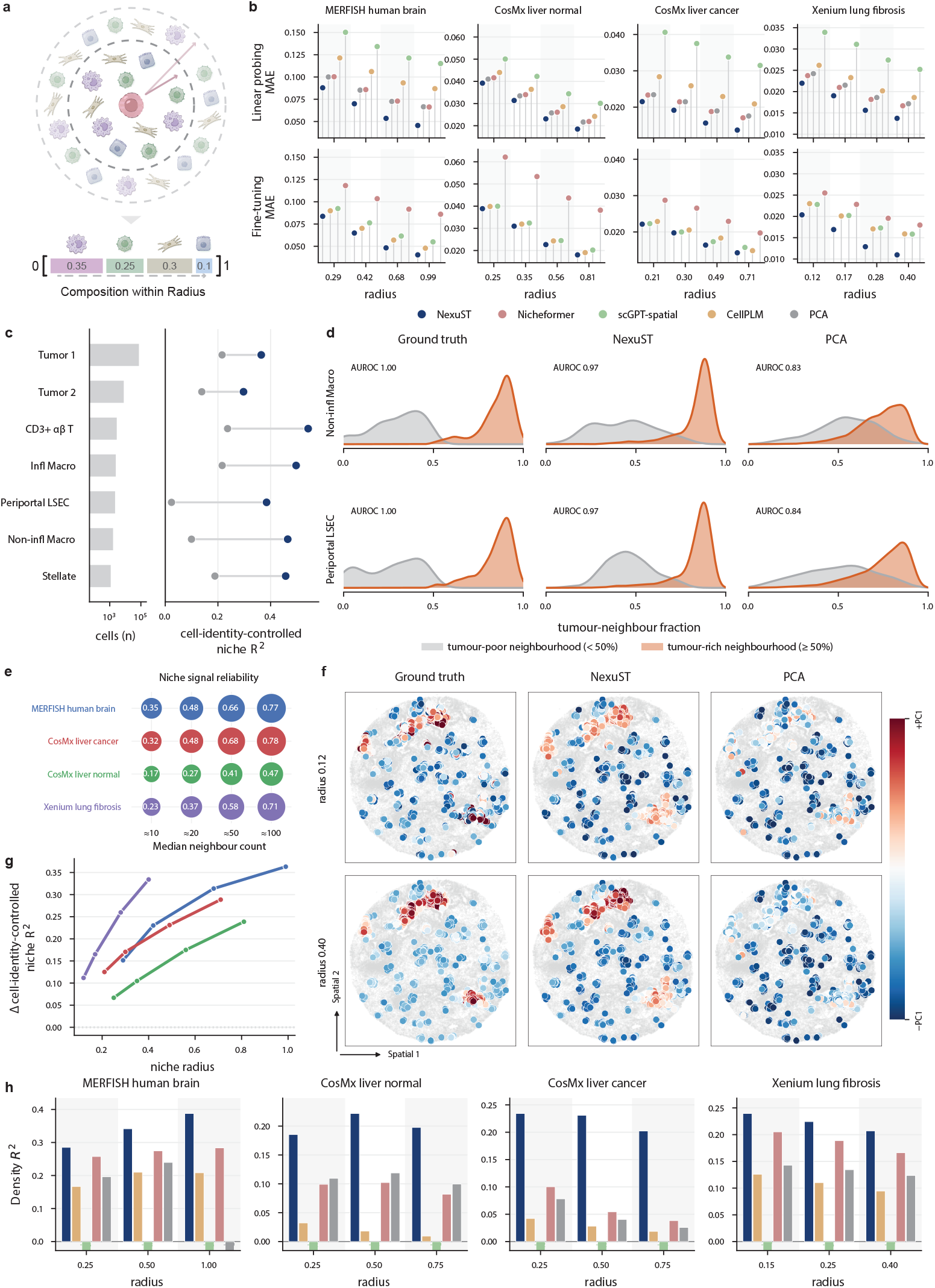
Niche composition prediction improves with spatial context and niche scale. **a**, Schematic of the neighbourhood-composition prediction task. For each focal cell, the neighbourhood composition is the relative abundance of cell types within a fixed spatial radius; the model predicts this composition vector from the focal cell’s pretrained representation. **b**, Niche composition prediction error as a function of neighbourhood radius across four spatial transcriptomics datasets, under linear probing (top) and full fine-tuning (bottom). Points show mean absolute error (MAE) for NexuST, Nicheformer, scGPT-spatial, CellPLM and PCA. **c**, Per-cell-type cell-identity-controlled niche *R*^2^ in CosMx liver cancer for NexuST (dark blue) and PCA (grey), with each pair joined by a line. The left panel shows the number of cells per type (log scale); cell types span the tumour, immune, endothelial and stromal lineages. **d**, Tumour-neighbour separability for two representative cell types (Non-infl Macro, Periportal LSEC) in CosMx liver cancer at the largest radius (0.71). Within each cell type, cells are split at the midpoint of the *true* tumour-neighbour fraction into tumour-poor (*<* 50%, grey) and tumour-rich (≥ 50%, orange) groups. Columns show the distribution of the tumour-neighbour fraction for the ground truth and for the NexuST- and PCA-predicted values; AUROC, quantifying how well the predicted score ranks tumour-rich above tumour-poor cells, labels each panel. NexuST recovers the ground-truth separation, whereas PCA collapses the two groups towards the type mean. **e**, Niche-signal split-half reliability as a function of niche size (median number of neighbours) across the four datasets; dot size and the printed value give the reliability, which increases as the niche enlarges. **f**, Representative spatial maps of within-type niche deviation (first principal component, *±* PC1) for SPP1^+^ macrophages in Xenium lung fibrosis at a small (0.12) and a large (0.40) radius, comparing the ground truth with NexuST and PCA predictions. **g**, Spatial advantage, the difference in cell-identity-controlled niche *R*^2^ between NexuST and PCA (Δ*R*^2^), as a function of neighbourhood radius across the four datasets. **h**, Local cell-density prediction *R*^2^ as a function of radius across the four datasets for NexuST, Nicheformer, scGPT-spatial, CellPLM and PCA. Local density is defined as the number of neighbouring cells within the radius of each cell.

NexuST achieved the lowest MAE across all four datasets, all four neighbourhood scales, and both linear probing and fine-tuning settings (Fig. 5b). Under linear probing, NexuST was the only model to outperform the expression-only PCA baseline at every dataset and neighbourhood scale. Under fine-tuning, NexuST remained the bestperforming method, although its margin narrowed on the two CosMx liver datasets. The ranking of the baselines also changed: CellPLM gained most and became the strongest baseline, whereas Nicheformer improved only marginally and became the lowest-performing method. In addition, MAE decreased with radius across all methods and settings, reflecting the fact that niche composition becomes progressively more stable at larger spatial scales (Fig. 5b and Methods Section 4.9.4). Absolute MAE values, however, are difficult to compare directly. Differences in MAE between datasets largely reflect their underlying composition distributions (e.g., the number and abundance of cell types) rather than model performance (Supplementary Fig. 19a). More fundamentally, MAE conflates two sources of predictability: the average niche associated with a cell’s identity, and the variation among the neighbourhoods of cells that share an identity. Separating them requires a different metric.

We therefore designed cell-identity-controlled niche *R*^2^, in which the mean neighbourhood composition of each cell type is used as the baseline, so that the reference is a predictor that knows a cell’s identity but nothing about its position (Methods Section 4.9.4). Under this metric, the apparent gain of PCA with increasing radius largely disappeared, and its R^2^ remained uniformly low — below 0.1 in both liver datasets and below 0.05 in the lung dataset (Fig. 5g and Supplementary Fig. 19c).

This indicates that, once the contribution of cell identity is removed, gene expression alone explains little of the residual variance in neighbourhood composition. In contrast, NexuST showed a consistent increase in cell-identity-controlled niche *R*^2^ with radius. Its performance was also substantially higher than that of PCA. For example, in the CosMx liver cancer dataset, NexuST achieved an *R*^2^ of around 0.6 at the largest radius (*r* = 0.71, *≈* 100 neighbours), representing a gap of nearly 0.5 compared with PCA (Supplementary Fig. 19c). Together, these results suggest that NexuST effectively leverages spatial information to recover neighbourhood composition beyond what can be inferred from gene expression alone.

To better understand the source of NexuST’s advantage across datasets, we further decomposed cell-identity-controlled niche *R*^2^ by cell type in all four datasets (Supplementary Figs. 21–24). In the CosMx liver cancer dataset, NexuST exceeded PCA for every cell type, with the same trend observed using raw niche-composition MAE (Fig. 5c; Supplementary Figs. 25–28). This indicates that, even after the average niche of each cell type had been removed, NexuST retained additional information about how cells of one type differ in their surroundings. Besides, the magnitude of the gain varied between cell types (Fig. 5c). The largest gains were observed for Periportal LSEC and non-inflammatory macrophages, both of which occupy diverse spatial niches despite relatively stable cell identities. In these cell types, neighbourhood composition is set by location rather than by cell identity, consistent with the near-zero R^2^ of PCA for these types. By contrast, tumour populations gained least, suggesting that much of their average neighbourhood composition is already captured by tumour identity itself. This is consistent with the biological expectation that tumour cells frequently occupy tumour-dominated microenvironments, making neighbourhood composition more predictable from cell identity alone.

To see what that residual signal is, we next examined the two cell types with the largest gains, i.e., non-inflammatory macrophages and Periportal LSECs (Fig. 5d). We quantified microenvironmental variation using the tumour-neighbour fraction, defined as the proportion of tumour cells within each cell’s neighbourhood. Cells within each type were divided into tumour-poor (*<* 50%) or tumour-rich (≥ 50%) groups according to their tumour-neighbour fraction (See Methods Section 4.9.4). The ground truth showed that both annotated populations spanned distinct tumour-poor and tumour-rich niches. NexuST largely recovered this separation (AUROC 0.97 for both cell types), whereas PCA produced overlapping predictions centred around the average tumour-neighbour fraction (AUROC 0.83–0.84). This pattern was also observed across neighbourhood radii, with clearer separation at larger radii (Supplementary Fig. 20). The relative insensitivity of PCA to neighbourhood scale prompted us to explore the biological basis of scale-dependent effects. We first implemented niche-signal split-half analysis, in which each cell’s neighbours were randomly halved and the two composition estimates were correlated (Methods Section 4.9.4). The reliability increased consistently with neighbourhood size across all four datasets, indicating that larger niches contain increasingly stable and reproducible biological information (Fig. 5e). To illustrate the biological basis of this effect, we examined SPP1^+^ macrophages in the Xenium lung fibrosis dataset (Fig. 5f; Supplementary Fig. 29). Neighbourhood-composition vectors were projected onto their first principal component, which mainly captured local macrophage self-aggregation. As the radius increased, the ground-truth aggregates became progressively more coherent and were recovered by NexuST, whereas PCA remained close to a spatially uniform prediction. Similar scale-dependent effects were also observed in smooth muscle cells/pericytes and secretory cells (Supplementary Figs. 30, 31). Matched ablation experiments further supported the contribution of cell-level modelling to niche-composition prediction: replacing cell-level attention with additional gene-level layers increased MAE across all four datasets at the largest neighbourhood radius (Supplementary Table 13c).

Finally, local organisation has a geometric component that composition does not capture. We predicted neighbourhood density, the number of cells within a given radius, capturing cellular packing rather than cell-type composition. Density distributions varied substantially across tissues, platforms and disease states, motivating us to ask whether pretrained representations retain this geometric information. We therefore trained a linear head on frozen embeddings to predict neighbourhood density across three radii (Methods Section 4.9.4; Supplementary Table 6). Among all methods, NexuST achieved the strongest performance across datasets and spatial scales, with particularly large gains in both liver datasets (Fig. 5h). NexuST therefore encodes not only which cell types surround a cell, but how densely they are packed.

### 2.6 Hierarchical attention progressively integrates cell-intrinsic and microenvironmental context

A key advantage of the NexuST architecture is that its hierarchical design enables attention analyses across multiple levels of biological organisation. In addition to gene-to-gene and cell-to-cell interactions, NexuST provides cell-to-gene attention links, allowing the exchange of information between molecular and cellular representations to be examined explicitly. To investigate whether the hierarchical attention learned by NexuST reflects biologically meaningful organisation, we analysed gene-level and cell-level attention patterns in the pretrained model without task-specific fine-tuning. We performed inference on five validation FOVs from each evaluation dataset and extracted attention weights from the corresponding gene-level and cell-level layers for analysis (Methods Section 4.10; Supplementary Table 10). The eight hierarchical blocks are referred to as L0–L7. Attention describes where the model reads; the analyses in this section are descriptive.

We first examined cell-to-gene attention across the pretrained gene layers using normalized Shannon entropy. Cell-to-gene attention is conditioned on the 300 gene-token keys and so sums to one (Methods, Eq.52). The Shannon entropy therefore measures how broadly a cell token distributes a fixed budget. Across all four datasets, attention became progressively more concentrated through the early layers, reached a pronounced minimum at L2, and then broadened toward deeper layers (Fig. 6a). This conserved pattern indicates that gene attention is systematically redistributed between L2 and L7 across cell types. To identify the genes underlying this redistribution, we compared L2 and L7 attention with mean expression in the Xenium lung fibrosis dataset (Fig. 6b,c; full gene-level attention matrices for all layers in Supplementary Fig. 32). At L2, attention was cell-type dependent and closely followed the focal cell’s transcriptional programme (Fig. 6c). For example, fibroblast populations showed high attention to collagen and extracellular-matrix genes such as *COL1A1* and *COL1A2*, whereas macrophage populations were characterized by high attention to macrophage markers including *CD68* and *MRC1*, consistent with their transcriptional identities. At L7, attention patterns diverged substantially from those observed at L2. Attention to several canonical cell-type markers fell, including *COL1A1* in fibroblasts, *CD68* in macrophages, and *SPP1* in SPP1+ macrophages. Similarly, *FN1*, a fibrosis-associated gene enriched in specific fibroblast and macrophage populations, also exhibited a marked decrease in attention. However, not all genes followed this trend. Notably, attention to *COL1A2* and, less markedly, *COL3A1* rose across multiple cell types. These observations indicate a systematic redistribution of attention in deeper layers and suggest that the features prioritized by the model at L7 are not solely determined by canonical cell-type-defining transcriptional programmes.

**Fig. 6:**
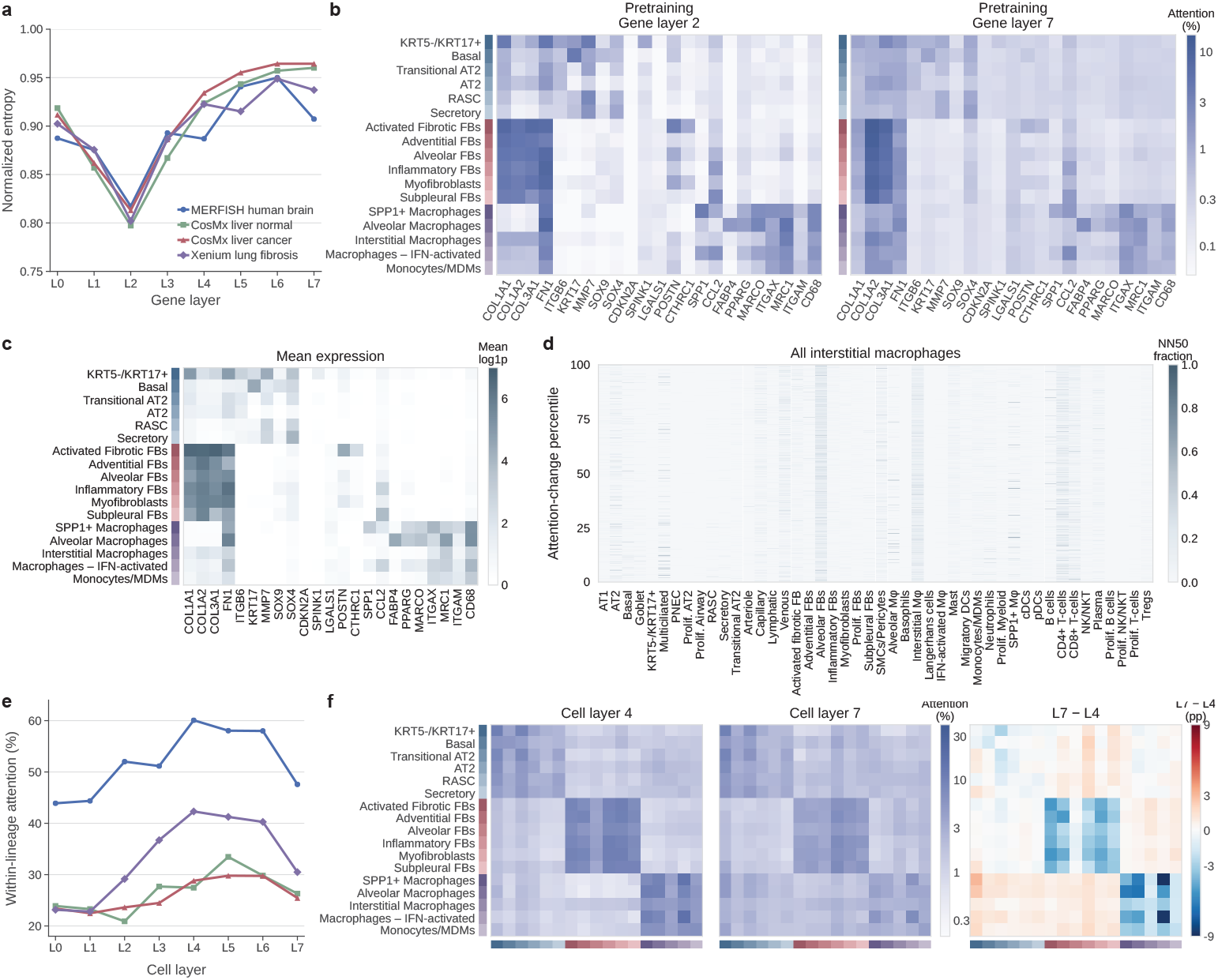
Layer-wise attention dynamics reveal progressive integration of molecular and spatial context. **a**, Normalized entropy of cell-token-to-gene attention across the eight gene layers in four held-out datasets. Lower entropy indicates attention concentrated on a smaller subset of genes. Attention is most concentrated at layer 2 and subsequently broadens toward deeper layers. **b**, Mean cell-token-to-gene attention for representative epithelial, fibroblast and macrophage populations in the Xenium lung fibrosis dataset at gene layers 2 and 7. Values were averaged across cells of each annotated cell type. Layer 2 corresponds to the minimum entropy observed in **a**, whereas layer 7 is the final gene layer. **c**, Mean log1p-normalised expression of the same genes and cell types shown in **b**, providing an expression reference for comparison with the attention patterns. **d**, Local neighbourhood composition of interstitial macrophages ordered by the change in *COL1A2* attention from gene layer 2 to layer 7. Within each FOV, macrophages were ranked by the L7–L2 attention change and converted to percentile ranks before being combined across FOVs. Heatmap values indicate the fraction of each cell type among the 50 nearest neighbouring cells. **e**, Fraction of cell-to-cell attention assigned to cells from the same major lineage across the eight cell layers in four held-out datasets. Within-lineage attention generally increases through intermediate layers before decreasing at the final layer. **f**, Mean cell-to-cell attention between representative cell types in the Xenium lung fibrosis dataset at cell layers 4 and 7, with their difference (L7 *−*L4, percentage points). Rows denote source cell types and columns denote target cell types; positive values indicate increased attention at layer 7.

Although genes showing reduced attention in L7 were frequently canonical cell-type markers, the biological significance of genes with increased attention remained unclear. We therefore focused on *COL1A2*. Across fibroblast subtypes, the increase in *COL1A2* attention did not follow the activation axis: adventitial fibroblasts, which are defined by anatomical position rather than by an activation programme, ranked alongside activated fibrotic fibroblasts, whereas subpleural fibroblasts changed little (Fig. 6b). This pattern is consistent with the possibility that *COL1A2* attention captures aspects of stromal organisation or local tissue context beyond canonical fibroblast activation programmes. More intriguingly, attention to *COL1A2* also rose across macrophage populations, although the magnitude of this increase was substantially smaller than that seen in fibroblast lineages. Notably, macrophages generally expressed little or no *COL1A2*. In contrast, *COL1A1* encodes the other chain of type I collagen, is co-regulated with COL1A2 in fibroblasts, and is likewise essentially unexpressed in macrophages, yet its attention changed little with depth. A general preference for abundant stromal transcripts would not separate two co-regulated, equally unexpressed genes.

Among macrophage subsets, interstitial macrophages exhibited the largest increase in *COL1A2* attention. We therefore focused subsequent analyses on interstitial macrophages. For each interstitial macrophage (IM), cells were ranked by their L7-L2 attention change. We then compared the cellular composition of the local neighbourhood, defined as the 50 nearest cells, between the top and bottom quartiles (Fig. 6d). Among fibroblast subsets, alveolar fibroblasts were the most enriched in the top quartile. In contrast, IMs themselves remained a relatively diffuse spatial distribution with little evidence of self-clustering. These findings suggest that increased *COL1A2* attention may be associated with differences in the surrounding stromal environment, supporting the possibility that deep-layer attention captures aspects of local niche organisation rather than the focal cell’s own expression of the gene.

We next asked whether an analogous reorganisation occurs between cells. We examined how cell-to-cell attention changed across layers and observed that attention in the intermediate layers increasingly concentrated on cell types from the same lineage. To quantify this pattern, we measured within-lineage attention across all four evaluation datasets. Consistent with this observation, within-lineage attention increased from the early to intermediate layers, reaching its highest levels around L4–L6 before declining at L7 (Fig. 6e). Comparing the L4 and L7 matrices in the Xenium lung fibrosis dataset (Fig. 6f; complete pretrained cell-to-cell attention matrices across all layers in Supplementary Fig. 33), several within-lineage attention blocks weakened at L7 while selected cross-lineage interactions increased. One of the clearest shifts occurred along the macrophage–fibroblast axis: total attention from alveolar macrophages to fibroblast populations rose from 3.74% at L4 to 10.25% at L7, while attention from interstitial macrophages increased from 10.01% to 14.55%. The increase was consistent across fibroblast subtypes (Fig. 6f). Importantly, this cell-level shift paralleled the gene-level *COL1A2* result: macrophages assigned greater deep-layer attention both to fibroblast cells and to the putative niche-associated gene *COL1A2*, a gene they barely express. Together, the gene- and cell-level analyses reveal a coordinated reorganization of attention across depth, from stronger organisation around cell-intrinsic and lineage-associated structure in intermediate layers to selective integration of information from the surrounding spatial microenvironment in deeper layers, highlighting how NexuST’s hierarchical attention progressively integrates cell-intrinsic and microenvironmental context.

## 3 Discussion

We introduce NexuST, a hierarchical foundation model for spatial transcriptomics that jointly models gene–gene dependencies within individual cells and cell–cell dependencies across tissue space, and lets the gene and cell levels interact in every encoder block throughout the end-to-end framework. Pretrained on HumanST-46M, a corpus spanning three imaging-based platforms and 11 organs, NexuST achieved the best or competitive performance across four downstream tasks on held-out datasets covering multiple platforms, organs, and tissue states, under both linear probing and full fine-tuning settings.

The ablation analyses indicate that explicit cell-level modelling is essential for capturing spatial context (Supplementary Note 4; Supplementary Table 13). Removing cell-level attention caused the largest degradation in region prediction and niche-composition modelling, while cell-type annotation was comparatively preserved, suggesting that cell-level attention is particularly important for encoding tissue-level spatial context beyond cell identity. How the gene- and cell-level layers are coupled also matters. An alternative design is to stack all gene-level layers before the cell-level layers, so that within-cell representations are formed first and spatial context is added afterwards. This sequential design remained highly competitive for cell-type annotation but was consistently weaker on region prediction and niche-composition modelling than the interleaved architecture used in NexuST. This suggests that repeatedly exchanging information between the gene and cell levels is particularly beneficial for integrating spatial context. This interpretation was further supported by cell-type-level analysis of the CosMx liver normal dataset, where the interleaved model showed clearer advantages for populations such as hepatocytes, whose discrimination depends more strongly on spatial context (Supplementary Fig. 13). Two-stage training, in which the gene layers are frozen before the cell layers are trained, was weaker still, indicating that the two levels are better learned jointly than introduced in sequence (Supplementary Table 13). The marked degradation after removing the library-size prediction head further supports the importance of explicitly factorising gene composition and library size, helping the model preserve cell-specific gene distributions rather than relying on a library-size shortcut (Methods Section 4.12; Supplementary Note 4).

The primary benchmark in this study used a common HVG300 input to control for differences in the gene-sequence lengths supported by different models and thereby enable a more comparable evaluation. Because this constraint may obscure an architecture’s ability to exploit larger panels, we also evaluated every model without HVG selection, allowing each to use the complete measured gene panel supported by its native input formulation. NexuST remained strongest across nearly all settings, so its advantage was not specific to the fixed HVG300 benchmark. More importantly, despite being pretrained with a fixed 300-gene token budget per cell, NexuST could directly accommodate larger gene panels during downstream evaluation without further pretraining or encoder adaptation, demonstrating its ability to generalise to broader gene panels as spatial transcriptomics assays become increasingly multiplexed (Methods Section 4.13; Supplementary Note 5).

A most notable observation from our evaluation is that PCA, despite using only cell-intrinsic expression and no explicit spatial information, was unexpectedly competitive with pretrained spatial foundation models under linear probing. Across all four tasks, including explicitly spatial tasks such as region prediction and niche composition prediction, PCA often ranked second only to NexuST. A strong expression-only baseline measures how much of a task is already answerable from cell identity. The performance of PCA suggests that a large part of the spatial signal is already reflected in each cell’s own transcriptome. In other words, spatial organisation is often tightly coupled to cell identity and state: if a model can recover what a cell is from expression, it can often infer what kinds of neighbour it has. The effect is especially clear for cell types with stereotyped niches. For example, tumour cells in the CosMx liver cancer dataset form spatially clustered aggregates, so their average neighbourhood composition is largely predictable from identity alone. Similarly, cholangiocytes are defined by canonical biliary markers and are recovered accurately even by PCA in cell-type annotation. However, expression alone has a clear limit, and NexuST’s advantage is concentrated exactly where identity is insufficient. Hepatocyte subtypes, for instance, lie along a zonated transcriptional continuum and remain difficult to separate when expression is interpreted without spatial context. Likewise, periportal LSECs occur in both tumour-poor and tumour-rich microenvironments, and PCA collapses toward a type-average neighbourhood prediction for them while NexuST recovers the separation. In these settings, the local tissue context provides information that is not fully recoverable from the individual cell transcriptome. Cell-intrinsic expression and spatial microenvironment are therefore complementary: cell identity explains a substantial fraction of the spatial signal, but spatial context resolves cases where identity alone is ambiguous. This supports the central design of NexuST: modelling gene-level and cell-level dependencies jointly.

NexuST also has several limitations. The first limitation is the pretraining corpus. Although HumanST-46M is one of the largest human-focused high-resolution spatial transcriptomics corpora to date, it is still bounded by the panels, platforms, tissues, and species covered by current datasets. Emerging assays combining high spatial resolution with broader transcriptome coverage would extend it, such as Visium HD. Besides, large scRNA-seq resources could provide useful prior knowledge for modelling gene–gene dependencies within cells. They may serve as a complementary source for improving gene-level representations before integrating spatial context even though they carry no spatial information. For example, a gene-level encoder could first be pretrained on large-scale scRNA-seq data and then used to initialize the gene layer of NexuST before spatial pretraining. A second limitation concerns the scope of the downstream evaluation. Our FOV-level splits assess within-dataset adaptation to held-out spatial fields, which is distinct from cross-slide or cross-donor transfer. Because training and validation FOVs may share section- or donor-specific characteristics, the current results should not be interpreted as evidence for transfer across independent biological samples. Assessing performance on slides or donors that are entirely excluded from downstream training would provide a complementary evaluation of such transferability.

A third limitation is computational scalability, which is bounded by the quadratic cost of self-attention. The full-panel analysis shows that NexuST can accommodate substantially expanded targeted panels. However, under standard Transformer self-attention, the number of genes and cells that can be jointly modelled is constrained by the quadratic cost of attention. This may limit the extension of NexuST to whole-transcriptome spatial assays, or to substantially larger tissue regions containing more cells. Future directions could explore more efficient attention variants, including linear attention [23], sparse attention [24], and window-based attention [25]. In addition, a Perceiver-style gene compression module [26–28] could be introduced before gene-level self-attention to compress large gene panels into a fixed number of latent gene tokens. This would allow the model to retain informative gene-level structure while scaling to broader transcriptomic inputs. These components could be further combined in a hierarchical architecture, where large gene panels are first compressed into latent gene tokens, efficient attention is applied at the gene level, and window-based or sparse attention is used at the cell level to process larger tissue regions. Such a design could increase both the gene input length and the spatial receptive field while keeping the overall computational cost tractable.

The pretraining objective is another limitation. NexuST uses a reconstruction-based objective. Although the factorised decoder reduced the tendency of absolute-expression regression to over-emphasize cell-wise library size, the objective remains a simplified approximation of the gene expression distribution. In particular, MSE-based reconstruction may not fully capture the count-like, sparse, compositional nature and even technical noise of transcriptomic measurements. This limitation motivates future objectives that define the self-supervised prediction task in the embedding space rather than directly in the raw expression space. By reducing the reliance on point-wise reconstruction of sparse and noisy measurements, such objectives may encourage the model to learn higher-level biological representations. Joint-embedding predictive architectures (JEPAs), which learn representations by predicting target embeddings rather than reconstructing raw inputs, provide one possible direction. JEPA-style objectives have shown promise in computer vision [29, 30] and have recently begun to be explored in single-cell transcriptomics [31, 32]. Extending this idea to spatial transcriptomics remains an important future direction, where the prediction target could include not only masked gene representations, but also spatial neighbourhood representations and tissue-context embeddings.

A broader limitation concerns the current maturity of the spatial transcriptomics foundation-model ecosystem. At present, benchmark tasks and evaluation protocols remain insufficiently standardized, particularly across gene-level, cell-level, and spatial-level prediction. And studies differ in datasets, annotations, splits, and baselines, making it difficult to tell whether reported performance gains reflect genuinely better spatial representation learning or dataset-specific biases. The two controls developed in this work — an expression-only baseline and an identity-controlled spatial metric — are offered as concrete steps towards evaluations that separate cell-intrinsic from spatial-context information. Future progress will therefore require more community-level benchmarks with clearly defined tasks, standardized held-out datasets, consistent data splits, strong non-spatial baselines, and evaluations that explicitly separate cell-intrinsic, gene-level, and spatial-context information. Moreover, current pretraining and evaluation settings are also largely static, such as cell-type annotation and region prediction. Although useful, these tasks do not fully capture dynamic biological processes such as temporal tissue remodelling, treatment response, perturbation effects, or counterfactual changes in cellular neighbourhoods. Progress in this direction is currently limited by the scarcity of large-scale spatial datasets with matched time-course, perturbation, longitudinal, or interventional measurements.

Overall, NexuST provides one of the first large-scale attempts to build a foundation model natively for spatial transcriptomics by jointly modelling molecular and spatial dependencies. By coupling gene–gene interactions within cells with cell–cell interactions across tissue space, NexuST moves beyond treating spatial transcriptomics as either independent single-cell profiles or purely spatial neighbourhood graphs. The ablations indicate that this coupling, and joint optimisation of the two levels, is where its advantage on spatial tasks comes from. This hierarchical design offers a principled backbone for learning representations that connect transcriptional state, cellular identity, and tissue microenvironment. Together with large-scale pretraining, systematic benchmarking, and biological interpretation, our study highlights a path toward the next generation of spatial transcriptomics foundation models. We anticipate that this gene–cell hierarchical modelling paradigm will serve as a reusable backbone for future models, helping to guide the development of more scalable, transferable, and biologically grounded spatial foundation models. More broadly, NexuST may support more generalizable tissue representation learning and help drive the transition from cell-level atlases to spatially resolved tissue atlases.

## 4 Methods

### 4.1 Pretraining Dataset curation and preprocessing

#### Pretraining data curation

We curated HumanST-46M from 72 datasets comprising 123 physical slides and 45.7 million cells across 11 organs, all generated by image-based single-cell-resolution spatial transcriptomics (Fig. 1). The corpus draws from two complementary sources. Public data releases from the three commercial vendors that dominate the field—Vizgen MERSCOPE [33], 10x Genomics Xenium [34], and Bruker CosMx SMI [35]—contribute 26.71M cells (58.5%) from 64 datasets, each corresponding to one physical slide. Peer-reviewed academic publications contribute 18.95M cells (41.5%) from 8 datasets comprising 59 physical slides. HumanST-46M therefore contains 72 datasets and 123 physical slides in total. It spans three image-based platforms: MERFISH (30.49M cells, 66.8%), 10x Genomics Xenium (11.84M, 25.9%), and Bruker CosMx SMI (3.34M, 7.3%). A complete list of physical slides, including per-slide cell counts, panel sizes, and source URLs, is provided in Supplementary Table 1.

#### Data preprocessing

Raw data were collected in their native vendor or publication formats and reorganised into a unified AnnData (`h5ad`) container. We first removed platform-specific control probes by name-pattern matching, including patterns such as `Blank, NegPrb`, and `FalseCode`, retaining only biological gene features; beyond this, we did not apply any expression-based filtering of cells or genes. Gene symbols across panels were harmonised by mapping non-standard symbols to HGNC-approved nomenclature, while preserving the panel-specific measured gene sets for each slide. Expression values were subsequently library-size normalised to 10,000 counts per cell followed by log(1 + *x*) transformation, while raw counts were preserved in `layers[“counts”]` to support downstream analyses and alternative preprocessing. Spatial coordinates were translated to start at zero along each axis and isotropically rescaled independently for each slide by its larger coordinate range, such that the longer spatial axis spans [0, 100] while preserving the slide’s aspect ratio. We did not apply highly variable gene selection; instead, all measured genes from each slide remain available to the model through the tokenisation scheme described in Section 4.3. Each slide was further annotated with organ and platform labels in `uns`.

### 4.2 Spatial sampling strategy

Slides in our training corpus, HumanST-46M, contain thousands to millions of cells, making whole-slide training computationally impractical for NexuST, in contrast to graph-based methods that typically operate on much smaller tissue graphs [1, 2]. To enable efficient large-scale training, we convert each slide into fixed-size local cell patches using a three-stage spatial sampling strategy (Supplementary Fig. 1). First, each slide is divided into spatially coherent subslides by applying *k*-means clustering to the two-dimensional cell coordinates. The number of clusters is chosen adaptively so that each subslide contains approximately 5,000 cells on average. This target size is defined relative to the model input of 512 cells. Second, within each subslide, 12 patch centres are selected by farthest-point sampling (FPS), which promotes spatially well-separated centres and reduces redundant sampling. Assigning the same number of centres to each subslide provides a comparable patch budget across spatial regions and promotes broad tissue coverage, rather than enforcing strictly uniform per-cell sampling within a single epoch. For each FPS centre, we construct a local patch from its 512 nearest neighbours in Euclidean space within the corresponding *k*-means subslide. Patches are allowed to overlap when their neighbourhoods intersect. FPS centres are independently resampled at each training epoch, progressively exposing the model to different cells and neighbourhood configurations. Empirically, cumulative unique-cell coverage rapidly approached complete coverage across all platforms and subslide-size quartiles, including the largest subslides (Supplementary Fig. 2). Each 512-cell patch is treated as one training sample, enabling efficient pretraining while preserving local tissue structure and maintaining a fixed input size.

### 4.3 Tokenization and input construction

After spatial sampling, each training sample is a local cell patch containing *N* = 512 cells. We first construct a global gene vocabulary *V*_gene_ from the union of genes in HumanST-46M, together with a dedicated padding token PAD, yielding |*V*_gene_| = 19,228 tokens in total. Because imaging-based spatial transcriptomics datasets are typically generated using predefined gene panels, each dataset or slide measures only a subset of this vocabulary. For a patch sampled from dataset or slide *d*, we denote its measured gene panel by *P*_*d*_ *⊆ V*_gene_ \ *{*PAD*}*.

During input construction, the raw count *c*_*i,g*_ of each measured gene *g ∈ P*_*d*_ in cell *i* is library-size normalised to a total of 10,000 across *P*_*d*_, yielding *u*_*i,g*_. A log1p transformation is then applied, giving the model input

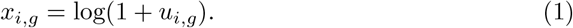

The resulting expression matrix for the patch is 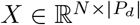, with entries *x*_*i,g*_. Genes outside *P*_*d*_ are unmeasured for slide *d* and are therefore neither instantiated as input tokens, nor looked up in the gene embedding table, nor treated as zero-expression genes.

#### Gene input construction

Given the measured gene panel *P*_*d*_ for slide *d*, NexuST uses a fixed gene-token budget of *M* = 300 to bound GPU memory and computational cost. Across HumanST-46M, 92.1% of pretraining cells contain no more than *M* = 300 detected genes, so this budget is sufficient to accommodate all detected genes for the large majority of cells. For cell *i*, let 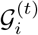 denote the fixed-length gene input used at pretraining step *t*:

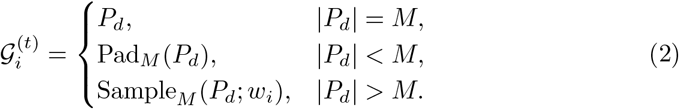

Here, Pad_*M*_ (*P*_*d*_) pads the measured gene panel to length *M* with the dedicated padding token PAD, with padded positions excluded from attention, and Sample_*M*_ (*P*_*d*_; *w*_*i*_) samples *M* genes without replacement from *P*_*d*_ independently for each cell exposure. For the sampling case |*P*_*d*_|*> M*, sampling is performed dynamically at every pretraining step in which the cell is used, with expression-aware weights:

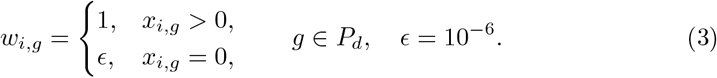

Thus, zero-expression genes in *P*_*d*_ are not removed from the measured panel or treated as missing values; they remain valid observed states in the sampling pool, while detected genes are strongly prioritised during input construction. Given *ϵ* = 10^*−*6^, cells with at most *M* detected genes are effectively represented with all of their detected genes in a single exposure, whereas, for cells with more than *M* detected genes, each exposure contains a dynamically resampled subset of those genes. Consequently, expression-aware sampling enriches detected genes relative to the complete measured panel and does not preserve its original zero-expression frequency in highly multiplexed cells. We adopt this strategy as a computational trade-off that prioritises observed molecular signals while maintaining a fixed gene-token budget. Across repeated exposures, dynamic resampling progressively broadens detected-gene coverage, with slide-level entry-weighted estimates at selected optimizer steps reported in Supplementary Table 3.

For simplicity, we omit the pretraining-step superscript *t* below and write the resulting input as

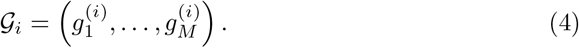

The valid non-padding positions are indexed by

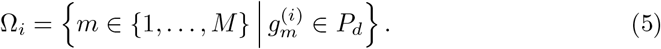

Thus, Ω_*i*_ indexes the genes instantiated in the current input. When |*P*_*d*_| *≤ M*, these positions cover the complete measured panel; when | *P*_*d*_ | *> M*, they correspond to the gene subset sampled for the current cell and pretraining step.

#### Gene ID embedding

Using the global gene vocabulary *V*_gene_ defined above, we learn a shared gene embedding table 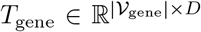, where *D* is the embedding dimension. For a cell *i* from slide *d*, we write the fixed-length gene input as 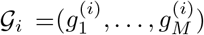, and map it to a gene ID embedding matrix 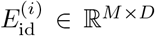 by table lookup. Specifically, each input position *m* is assigned a *D*-dimensional embedding vector:

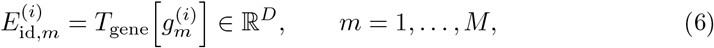

The gene ID embedding matrix for cell *i* is therefore constructed as

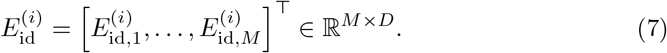

At the patch level, the cell-wise embedding matrices are collected as

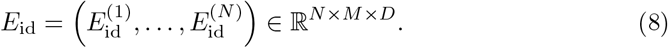

#### Gene value embedding

For each valid position *m ∈* Ω_*i*_, the corresponding measured gene 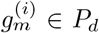 is aligned with its log-normalised expression value 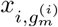 in cell *i*. We then apply a lightweight MLP *f*_val_ : ℝ *→* ℝ^*D*^ to map each such value:

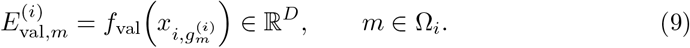

For padding positions *m ∈/* Ω_*i*_, a placeholder value embedding is used and the position is excluded from attention, whereas zero-expression measured genes remain valid observed states with expression value zero. Collecting the embeddings across all *M* positions gives the gene value embedding matrix for cell *i*:

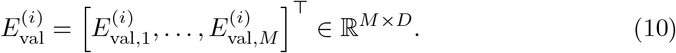

At the patch level, the cell-wise value embedding matrices are collected as

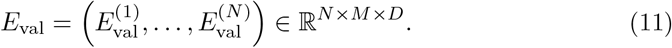

#### Organ token

To incorporate organ information into the model input, we construct an organ vocabulary *O* from HumanST-46M, comprising 11 organs. Similar to the gene embedding table, we learn an organ embedding table *T*_organ_ *∈* ℝ^|*O*|*×D*^. For a patch from organ *o*_*d*_ *∈ O*, we obtain its organ embedding as

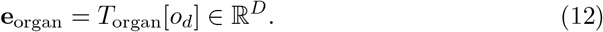

This embedding serves as the organ token, is shared by all cells in the patch, and is broadcast along the cell dimension to form

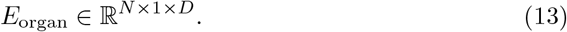

#### Input construction

For each cell *i*, we form its gene token embeddings by adding the gene ID and gene value embeddings:

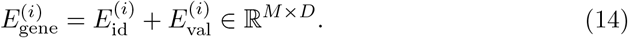

Padding positions are masked out in gene-level attention. We then prepend a learnable cell token **c**_*i*_ ∈ ℝ^*D*^ to cell *i*, and insert the organ token **e**_organ_ ∈ℝ^*D*^, which is shared by all cells from the same patch. The initial token representation of cell *i* is therefore

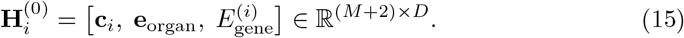

Stacking the resulting representations for all *N* cells gives the initial encoder input for the patch:

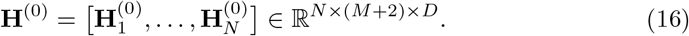

### 4.4 NexuST architecture

#### Hierarchical encoder

For mini-batch training, we stack *B* patch inputs along a batch dimension and continue to denote the resulting tensor by **H**^(0)^:

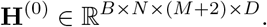

NexuST applies *S* stacked hierarchical blocks, each consisting of a gene-level layer followed by a cell-level layer, both implemented as standard pre-norm Transformer encoder layers. This factorisation aligns with the two scales of a tissue—gene–gene coordination within each cell and cell–cell interaction across the patch—and avoids the *O*((*NM*)^2^) cost of attending over all tokens jointly.

Before the first block, two-dimensional sinusoidal positional embeddings derived from the cell coordinates are added only to the cell-token states. For a single patch, these states are collected as

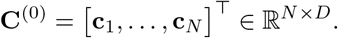

By stacking *B* patches, we use the same notation for the batched cell-token states,

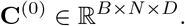

The positional embeddings are then added as

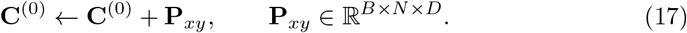

The updated cell tokens are placed back into the corresponding cell-token positions of **H**^(0)^. Gene tokens receive no positional encoding, as genes within a cell form an unordered set and spatial position is meaningful only at the cell level.

After adding positional embeddings to the cell-token states, the input is passed through *S* hierarchical blocks. At block *s* ∈ {1, …, *S*}, gene-level attention is applied independently within each cell. We first merge the batch and cell dimensions and then apply the gene-level Transformer layer:

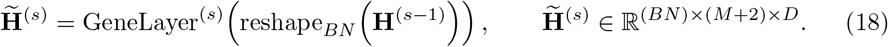

Thus, after the gene-level layer, each cell token summarises the gene-level information of its corresponding cell, conditioned on the organ token. The resulting hidden states are reshaped back to ℝ^*B×N ×*(*M* +2)*×D*^; for simplicity, we continue to denote the reshaped tensor by 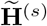. The corresponding cell-token states are then extracted as

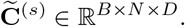

The cell-level layer then allows attention across the *N* cells within each patch:

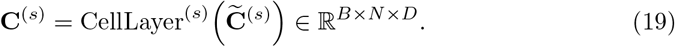

The updated cell tokens **C**^(*s*)^ are written back to the corresponding cell-token positions of **H**^(*s*)^, forming the next block input 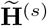.

Across the *S* hierarchical blocks, the cell token serves as the bridge between within-cell gene modelling and cross-cell spatial contextualisation. Information aggregated by the cell-level layer is carried back into the next block, where the spatially informed cell tokens participate in gene-level attention within each cell and modulate the gene-token representations. After the final block, the cell tokens are layer-normalised to obtain the cell representations used for downstream tasks.

#### Batch-conditioned decoder context

The decoder reconstructs gene expression from the cell tokens produced by the hierarchical encoder. Let

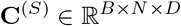

denote the final layer-normalised cell representations. To account for batch effects arising from platform-specific technical variation, we concatenate these representations with learned batch or platform embeddings. The embeddings for all cells are stacked as

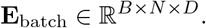

The decoder context is then obtained by concatenating **C**^(*S*)^ and **E**_batch_ along the feature dimension, yielding **Z** ∈ ℝ^*B×N ×*2*D*^. For simplicity, we use **z**_*i*_ ∈ ℝ^2*D*^ below to denote the decoder context of a single cell. The final updated gene-token states are not passed directly to the decoder; instead, static gene ID embeddings serve as gene-specific queries conditioned on **z**_*i*_, as detailed below.

#### Factorised decoder

Our self-supervised objective reconstructs masked entries of the log-normalised expression *x*_*i,g*_ using an MSE loss; the masking procedure and loss definition are described below. Using the corresponding pre-log expression *u*_*i,g*_ defined above, we define the sample-restricted library size for cell *i* as

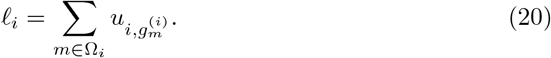

Here, *l*_*i*_ is defined over the current valid gene set after library-size normalisation and therefore does not denote the original raw-count library size. The sampled pre-log expression profile can then be factorised as

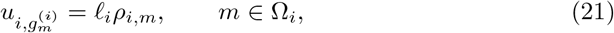

where 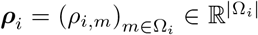 denotes the relative gene-expression composition over the current valid gene set, with

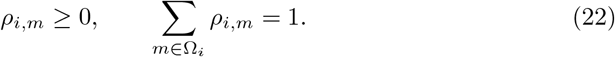

The corresponding log-normalised target is therefore

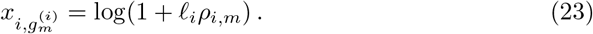

When the sample-restricted library size *l*_*i*_ varies across cells, this dependence can create a library-size shortcut under the MSE objective: a decoder can reduce reconstruction loss by accurately capturing *l*_*i*_, while replacing the cell-specific composition *ρ*_*i,m*_ with a gene-specific average composition. We denote by 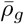 the average relative composition associated with gene *g* across cells in which that gene is instantiated.

Under this shortcut, the decoder may produce

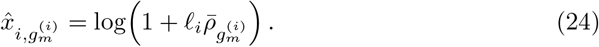

The resulting predictions preserve global expression magnitude but suppress the cell-specific compositional variation that the learned representation is intended to encode (Supplementary Note 1).

We therefore use a factorised decoder comprising a dense mixture-of-experts (MoE) head for the predicted normalised gene composition 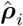 and a lightweight head for the predicted sample-restricted library size 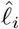, encouraging the model to capture cell-specific gene distributions rather than relying on the library-size shortcut.

#### MoE composition head

We employ a dense mixture-of-experts (MoE) composition head to predict the gene-wise composition ***ρ***_*i*_. The routing mechanism is gene-specific.

For each valid gene position *m ∈* Ω_*i*_, the gating network takes the gene ID embedding 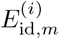 as input and produces a probability distribution over *K* experts:

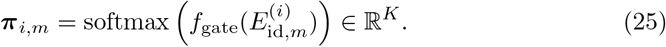

Here, *π*_*i,m,k*_ denotes the routing weight assigned to the *k*-th expert, with 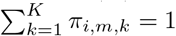. This allows the expert mixture to vary across genes.

Each lightweight expert takes the gene ID embedding 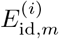 and the cell-level decoder context **z**_*i*_ as inputs and produces a scalar composition logit. Rather than activating only a subset of experts through sparse routing, our dense routing scheme evaluates all *K* experts and computes their gating-weighted sum:

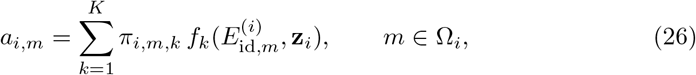

where *f*_*k*_(*·*) denotes the scalar logit produced by the *k*-th expert. The normalised gene composition is predicted by applying a softmax to these logits over the current valid non-padding positions:

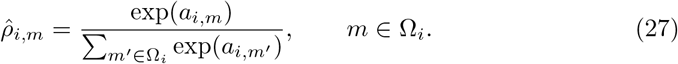

#### Library-size head and reconstruction

In parallel with the MoE composition head, a lightweight library-size head predicts a positive sample-restricted library size from the decoder context **z**_*i*_:

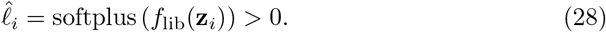

For each valid position *m ∈* Ω, corresponding to gene 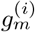, the predicted pre-log expression is obtained by combining the predicted library size and gene composition.

For notational simplicity, we write the reconstruction loss for a single patch; during mini-batch training, the loss is averaged over all patches in the batch:

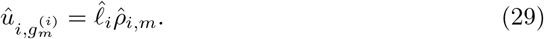

The final log-normalised reconstruction is then

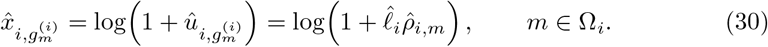

### 4.5 Pretraining

#### Masked gene-value reconstruction objective

We adopt masked gene-value reconstruction as the self-supervised pretraining objective. Given the fixed-length gene input 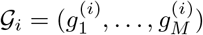 for cell *i*, recall that the valid non-padding positions are

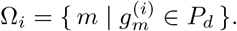

For each cell, we randomly select 20% of the valid positions for masking, denoted by Ω_*i*,mask_ *⊆* Ω_*i*_. For each *m ∈* Ω_*i*,mask_, the gene identity 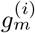 is kept unchanged, while its value embedding 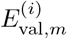 is replaced by a learnable `[MASK]` value embedding. Each masked position therefore retains its gene identity, while its corresponding scalar expression value serves as the reconstruction target.

Using the decoder prediction 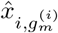 defined above, we compute the mean squared error over the masked valid positions:

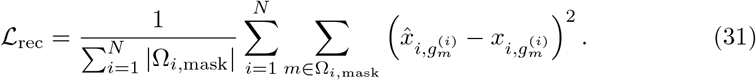

For panels with |*P*_*d*_| *> M*, gene sampling is performed before masking, and the loss is computed only over the masked subset of the sampled genes. When sampled, measured genes with zero expression remain valid reconstruction targets with target value zero.

#### Pretraining strategy

We pretrain NexuST using 8 hierarchical encoder blocks. Each block contains one gene-level Transformer layer and one cell-level Transformer layer, resulting in 16 Transformer layers in total. The embedding dimension is set to *D* = 512, with 8 attention heads and a dropout rate of 0.1. The dense MoE decoder contains 8 experts. The complete pretraining model contains 67.9M trainable parameters, including the encoder and reconstruction decoder. The decoder is discarded for downstream evaluation, leaving an encoder with 60.6M parameters.

Each training patch contains *N* = 512 cells, and each cell is represented by a fixed-length gene input of *M* = 300 genes. The gene value masking ratio is set to 20%. We optimize the model using AdamW with a learning rate of 3 *×* 10^*−*4^, weight decay of 0.01, *β*_1_ = 0.9, and *β*_2_ = 0.999. Gradient clipping is applied with a maximum norm of 1.0. The model is trained for 10,000 steps, with 1,000 warm-up steps followed by cosine learning-rate decay for the remaining 9,000 steps. The minimum learning rate is set to 10^*−*6^.

Large-scale pretraining is performed on 32 NVIDIA A100 64GB GPUs across 8 nodes, with 4 GPUs per node. We use a per-GPU batch size of 6, resulting in a global batch size of 192. To reduce GPU memory consumption, we use fully sharded data parallelism (FSDP), which shards model parameters, gradients, and optimizer states across GPUs. We further enable activation checkpointing and use mixed-precision training with bfloat16. The full pretraining takes approximately 40 hours. The detailed pretraining configuration is summarized in Supplementary Table 4. The corresponding training pseudocode is provided in Supplementary Note 2.

### 4.6 Inference

Whole-slide inference is challenging because a slide may contain from thousands to millions of cells, whereas the model takes a fixed-size local patch as input. The stochastic spatial sampling strategy used during pretraining, described in Section 4.2, provides local context and data augmentation, but may produce overlapping patches and is therefore not ideal for deterministic whole-slide inference.

To enable efficient and non-overlapping inference, we order all cells in a slide using a Hilbert curve, a space-filling curve that maps two-dimensional spatial coordinates to a one-dimensional sequence while approximately preserving local neighborhoods [36]; see Supplementary Note 3. Cells are sorted by their Hilbert indices and divided into consecutive segments, each containing *N* = 512 cells to match the pretraining setting. If the final segment contains fewer than 512 cells, it is padded to the required length, and padded cells are excluded from downstream output aggregation. At inference time, we use only the encoder to compute cell-level representations, while the pretraining decoder and reconstruction heads are discarded. This allows the model to scan the entire slide efficiently while avoiding redundant computation from overlapping windows.

For the gene dimension, we keep the inference protocol consistent with pretraining. For each target dataset, we select a fixed set of 300 highly variable genes from its measured gene panel and apply the same preprocessing pipeline. Zero-expression genes are retained if they belong to the selected measured gene set, whereas genes outside the measured panel are not treated as zero-expression inputs. This preserves the distinction between true zero expression and structural gene absence.

### 4.7 Downstream datasets

We evaluated NexuST on four held-out benchmark datasets spanning the three image-based platforms represented in HumanST-46M: MERFISH human brain [37], CosMx liver normal, CosMx liver cancer, and Xenium lung fibrosis [38]. The two CosMx liver benchmarks were derived from the same public Bruker CosMx SMI Human Liver FFPE dataset [39], which contains one normal liver sample and one liver cancer sample. All four datasets were processed using the same control-probe removal, gene-symbol harmonization, expression normalization, and coordinate normalization pipeline as the pretraining corpus. Dataset-specific details, including sample origin, panel size, cell count, and available annotations, are described below.

#### MERFISH human brain

We collected the MERFISH human brain dataset from [37], deposited at Zenodo (https://zenodo.org/records/14422018). The dataset consists of a single section of primary visual cortex (Brodmann area 17, sample `adult umb5958`) from one adult donor, measured with a 300-gene MERFISH panel and yielding 218,099 cells. Each cell is annotated with 8 cell type labels as provided by the original publication: EN-UL, EN-DL, EN-L2, IN, Astro, Olig, Glia, EC; no region annotation is provided. The fetal cortex samples from the same publication are included in HumanST-46M for pretraining; the pretraining and evaluation samples share only the 300-gene panel design, with no donor, section, or developmental stage reused, and the adult section used here is entirely held out from pretraining.

#### CosMx liver normal

We collected the CosMx liver normal dataset from the Bruker CosMx SMI Human Liver FFPE public release [39] (https://brukerspatialbiology.com/products/cosmx-spatial-molecular-imager/ffpe-dataset/human-liver-rna-ffpe-dataset/). The dataset consists of a single tissue section from one healthy donor (male, 35 years old), measured with the 1000-plex CosMx Human Universal Cell Characterization Panel (999 genes) and yielding 332,877 cells across 301 vendor-defined fields of view. Each cell is annotated with 18 cell type labels and 6 region labels as provided by the public release. The 18 cell type labels comprise five hepatocyte subtypes (Hep 1, Hep 3, Hep 4, Hep 5, Hep 6), stellate cells, cholangiocytes, portal endothelial cells, periportal liver sinusoidal endothelial cells (LSECs), central venous LSECs, CD3+ alpha beta T cells, gamma delta T cells, NK-like cells, mature B cells, antibody-secreting B cells, inflammatory macrophages, non-inflammatory macrophages, and erythroid cells. The 6 region labels follow the canonical zonation of the hepatic lobule along the portal-central axis: portal vein (zone 1a), zone 1b, zone 2a, zone 2b, zone 3a, and central vein (zone 3b).

#### CosMx liver cancer

Following the CosMx liver normal dataset above, we collected the CosMx liver cancer dataset from the same public release [39]. The dataset consists of a single tissue section from one hepatocellular carcinoma patient (female, 65 years old; grade G3, stage II), measured with the same 1000-plex CosMx Human Universal Cell Characterization Panel (999 genes) and yielding 460,441 cells across 383 vendor-defined fields of view. Each cell is annotated with 17 cell type labels and 4 region labels as provided by the public release. The 17 cell type labels comprise two tumor populations (tumor 1, tumor 2), CD3+ alpha beta T cells, gamma delta T cells, NK-like cells, mature B cells, antibody-secreting B cells, inflammatory macrophages, non-inflammatory macrophages, erythroid cells, stellate cells, cholangiocytes, portal endothelial cells, periportal liver sinusoidal endothelial cells (LSECs), central venous LSECs, hepatocytes (Hep), and an undetermined category (NotDet). The 4 region labels partition the tissue section into compartments of the tumor microenvironment: tumor, tumor subtype, interface (the boundary between malignant and non-malignant tissue), and non-malignant.

#### Xenium lung fibrosis

We collected the Xenium lung fibrosis dataset from Vannan et al. [38], deposited at the Gene Expression Omnibus (GEO) under accession GSE250346. The dataset consists of 45 tissue samples from 35 donors, comprising 9 unaffected donors and 26 donors with pulmonary fibrosis (PF) spanning eight clinical diagnoses: idiopathic pulmonary fibrosis (IPF, *n* = 12), interstitial lung disease (ILD, *n* = 4), chronic hypersensitivity pneumonitis (cHP, *n* = 4), connective tissue disease– associated ILD (CTD-ILD, *n* = 2), and one donor each with combined pulmonary fibrosis and emphysema (CPFE), nonspecific interstitial pneumonia (NSIP), interstitial pneumonia with autoimmune features (IPAF), and sarcoidosis. All samples were measured with a 342-gene Xenium panel, yielding 1,630,319 cells in total. Each cell is annotated with 47 cell type labels and 12 region labels as provided by the original publication. For lineage-based region and attention analyses, we grouped the 47 cell types into six broad lineages: endothelial (4 cell types), epithelial (12), fibroblast/stromal (8), mural (1), myeloid (13) and lymphoid (9). These lineages cover both canonical healthy lung populations (e.g., AT1, AT2, alveolar fibroblasts, capillary endothelial cells, alveolar macrophages) and PF-emergent populations (e.g., KRT5^*−*^/KRT17^+^ aberrant basaloid cells, transitional AT2 cells, activated fibrotic fibroblasts, SPP1^+^ macrophages); the complete mapping and cell type list are provided in Supplementary Table 12. The 12 region labels correspond to the 12 cell-based niches (C1–C12) identified by Vannan et al. [38]; we relabel them with descriptive names reflecting their dominant cell composition and tissue context (Supplementary Table 11): Healthy alveolar, AT2-enriched alveolar, Transitional epithelial, Epithelial detachment, Air-way, Adventitial/vascular, Perivascular, Interstitial fibrotic, T/plasma immune, Mixed immune, Lymphoid/TLS, and Macrophage accumulation.

### 4.8 Downstream evaluation protocol

For each downstream benchmark, we evaluated models within the corresponding dataset rather than under a leave-one-dataset, leave-one-donor, or leave-one-platform transfer setting. We constructed training and validation partitions by field of view (FOV) rather than by random cell sampling. This design reduces cell-level spatial leakage caused by randomly assigning neighbouring cells to different partitions, while preserving a consistent evaluation protocol across methods. Specifically, 80% of FOVs were randomly assigned to training and the remaining 20% to validation. The Xenium lung fibrosis dataset provides vendor-defined FOVs, which we used directly for splitting. The CosMx liver normal and CosMx liver cancer datasets are single-section benchmarks with pre-defined acquisition FOVs, which we used as within-section partitions. The MERFISH human brain dataset does not include vendor-defined FOVs, so we partitioned the tissue into a 10 *×* 10 grid of pseudo-FOVs prior to splitting. The resulting partitions were identical across methods. Within each dataset, a common set of 300 highly variable genes (HVGs) was selected from the full unlabelled expression matrix before the FOV split and used as the candidate input panel for all methods. This transductive choice reflects our treatment of HVG selection as dataset-level preprocessing. Using the same candidate panel across training and validation FOVs and methods reduces feature-selection variability. Gene identifiers were then converted to each method’s expected format and intersected with its pretrained vocabulary. Consequently, the final input gene sets were not identical across models, although most model–dataset combinations retained more than 90% of the shared HVG300 panel; exact coverage is reported in Supplementary Table 7. All results are reported as mean *±* s.d. over three random seeds.

For each applicable task–dataset pair, all learnable foundation models were evaluated under both linear probing and fine-tuning. Cell-type annotation, held-out gene recovery, and neighbourhood-composition prediction were evaluated on all four datasets, whereas region prediction was evaluated on the three datasets with region annotations. In both regimes, the same linear head was attached to the encoder output; the regimes differed only in whether the encoder was frozen (linear probing) or jointly trained (fine-tuning). PCA, which has no pretrained encoder, was treated as a non-pretrained baseline whose features were passed to the same linear head under the linear-probing configuration. Under linear probing, we used identical hyperparameters across all methods: batch size 64, weight decay 1 *×*10^*−*6^, and task-specific learning rates (1 *×*10^*−*3^ for regression tasks including niche composition prediction and held-out gene recovery, 1 *×*10^*−*2^ for region prediction, and 1 *×*10^*−*1^ for cell-type annotation), reflecting the differing optimisation scales of regression and classification objectives (Supplementary Table 8). Training was conducted for up to 50 epochs, with early stopping based on the validation task metric and a patience of 5 epochs.

Under fine-tuning, tokenisation schemes, input pipelines, and memory requirements differed substantially across foundation models. We therefore did not enforce a single batch size or learning rate across methods. Instead, we used the original or officially recommended model-specific settings when available and adopted the most stable setting under our shared downstream training framework when such settings were not provided, as in scGPT-spatial. All models were trained for up to 50 epochs, with early stopping based on the validation task metric. Model- and task-specific learning rates are reported in Supplementary Table 9. In both regimes, the checkpoint achieving the best validation metric was retained, and its performance on the same validation FOVs was reported.

### 4.9 Downstream task

#### 4.9.1 Cell-type annotation

##### Task definition and evaluation

For cell-type annotation, each model representation **h**_*i*_ was used to predict the annotated cell-type label *y*_*i*_ through a task-specific linear classification head. The head was trained with cross-entropy loss, and performance was evaluated using macro F1 to account for class imbalance across cell types. Cell-type-level F1 scores and confusion matrices were further used for detailed error analysis.

#### 4.9.2 Region prediction

##### Task definition and evaluation

For region prediction, each model representation **h**_*i*_ was used to predict the annotated tissue-region label *r*_*i*_ through a task-specific linear classification head. The head was trained with cross-entropy loss, and performance was evaluated using macro F1 across annotated regions. In addition to the overall macro F1, we reported per-region F1 scores and confusion matrices to analyse which tissue regions were correctly resolved or confused by each method.

##### Cell-type-stratified region-prediction accuracy

To examine whether model gains were concentrated in cell types whose regional identity was weakly determined by cell identity alone, we further computed cell-type-stratified region-prediction accuracy. For model *m*, let 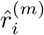 denote the predicted region of cell *i, r*_*i*_ its ground-truth region, and *t*_*i*_ its annotated cell type. For each cell type *c*, we computed

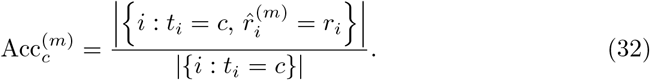

This region-marginalised accuracy measures the fraction of cells of a given type that were assigned to their correct tissue region.

We also computed a region-stratified version for case-study analyses. For each region *r* and cell type *c*, we computed

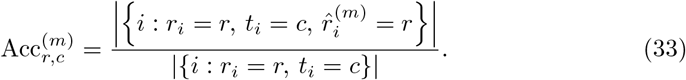

This measures the fraction of cells of type *c* within ground-truth region *r* that were correctly assigned to region *r*. Cell-type and region–cell-type groups containing fewer than 30 and 20 cells, respectively, were excluded from these stratified analyses.

##### Cell-type regional dispersion

To quantify how strongly cell identity constrains regional location, we measured the dispersion of each cell type across annotated tissue regions using the ground-truth cell-type and region annotations of the held-out Xenium lung fibrosis dataset. For a cell type *c*, let *n*_*rc*_ be the number of cells of type *c* annotated to region *r*. Column-normalising the count vector gives the conditional region distribution

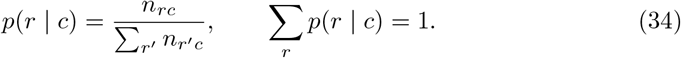

We then quantified regional dispersion using the Shannon entropy

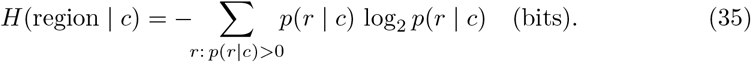

Low entropy indicates that a cell type is largely restricted to a small number of regions, so its regional location is strongly constrained by cell identity. High entropy indicates that a cell type is dispersed across multiple tissue contexts, so its regional assignment is less determined by cell identity alone and is expected to depend more strongly on local spatial context. This entropy was used to analyse the relationship between cell-type regional dispersion and NexuST’s gain over baseline models in region prediction.

#### 4.9.3 Held-out gene recovery

##### Task definition and target-gene selection

Starting from each dataset’s shared HVG300 candidate panel defined in the downstream evaluation protocol, we first took the intersection of genes present in the pretrained vocabularies of all four foundation models—NexuST, scGPT-spatial, Nicheformer, and CellPLM—to define a single cross-model gene set. We then randomly selected 20% of this common set, using a fixed seed of 42, as the held-out target genes for all methods. Because vocabulary filtering was performed before target selection, the number of held-out genes was dataset specific and could be fewer than 60, as reported in Supplementary Table 7. The remaining genes formed the observed input panel. Thus, all methods received the same observed gene identities and were evaluated on the same held-out target genes. The held-out genes were completely removed from the encoder input before tokenisation and were retained only as regression targets. For each method, a linear regression head mapped the resulting cell representation to the expression values of all held-out genes simultaneously and was trained using mean squared error.

##### Gene-wise recovery metrics

We evaluated held-out gene recovery from two complementary perspectives: recovery of the expressing-cell pattern and recovery of expression values. All metrics were first computed separately for each held-out gene across validation cells. Rank-based metrics quantify whether the relative expression pattern across cells is preserved, whereas error-based metrics quantify the deviation between predicted and observed expression values.

##### Spearman correlation

For each held-out gene *g*, we computed the Spearman rank correlation between its predicted and observed expression values across all validation cells:

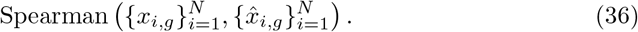

Zero-expression cells were retained in this calculation. Consequently, the metric reflects both the ability to distinguish expressing from non-expressing cells and the ability to recover relative expression levels among cells. Dataset-level performance was summarised across held-out genes.

##### Jaccard similarity of expressing-cell pattern recovery

To evaluate whether a model correctly identified the cells expressing each held-out gene, ground-truth expression was binarised using *x*_*i,g*_ *>* 0. For gene *g*, let

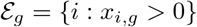

denote the set of expressing cells and let *k*_*g*_ = |*ε*_*g*_|. Genes with fewer than 10 expressing validation cells, or with expression in all validation cells, were excluded from this analysis. The predicted expressing set 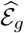 was defined as the *k*_*g*_ cells with the highest predicted expression values. The Jaccard similarity was then computed as

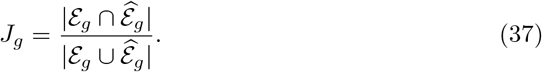

This top-*k*_*g*_ construction fixes the predicted prevalence to the observed prevalence and therefore evaluates recovery of the expressing-cell pattern using the ranking of predicted expression values, independently of the absolute prediction scale or a model-specific threshold. For comparison, we defined a prevalence-matched random baseline by replacing the overlap with its expectation under random selection of *k*_*g*_ cells, yielding

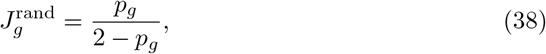

where *p*_*g*_ = *k*_*g*_*/N* is the fraction of validation cells expressing gene *g*.

##### Gene-sparsity-stratified recovery analysis

For each held-out gene *g*, gene sparsity was defined as

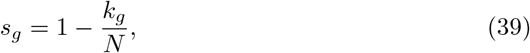

where *k*_*g*_ is the number of validation cells with *x*_*i,g*_ *>* 0, and *N* is the total number of validation cells in the corresponding dataset. Genes were grouped into six sparsity intervals:

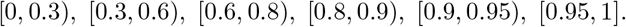

The high-sparsity range was divided more finely because most genes in the imaging-based panels were concentrated between sparsity values of 0.8 and 1.0. For each model, dataset, and sparsity interval, we reported the mean gene-wise Spearman correlation across all genes assigned to that interval. Intervals containing no genes were treated as missing values and were not interpolated, so lines in the corresponding plots were broken across empty intervals.

##### Mean absolute error

For each held-out gene *g*, mean absolute error was computed across validation cells as

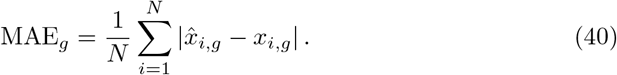

MAE was calculated on the same normalised and log-transformed expression scale used for model input and regression targets. Dataset-level performance was summarised across the held-out genes.

#### 4.9.4 Niche composition prediction

##### Calculation of niche composition

For each dataset, niche compositions were computed independently for each field of view and each spatial radius. Within a field of view containing *n* cells, we constructed a binary radius graph using the provided two-dimensional spatial coordinates. Two cells were connected if their Euclidean distance was within the specified radius, with self-loops excluded. Let *A* ∈ {0, 1}^*n×n*^ denote the resulting adjacency matrix, and let *Y* ∈ {0, 1}^*n×L*^ denote the one-hot cell-type matrix under a fixed dataset-wide ordering of the *L* annotated cell types. The neighbor cell-type count matrix was calculated as

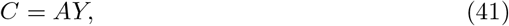

where *C*_*il*_ denotes the number of neighbouring cells of type *l* surrounding cell *i*. These neighbour counts were then normalised by their row-wise sums to obtain the target niche compositions:

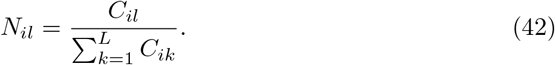

This yielded *N* ∈ [0, 1]^*n×L*^, where each row represents the cell-type composition of the spatial neighbourhood surrounding one cell and satisfies

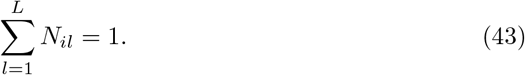

Because the cellular microenvironment is inherently scale-dependent, we evaluated niche composition prediction at four neighbourhood scales. To account for variation in cell density across platforms, tissues, and disease states, we selected dataset-specific radii corresponding to approximately 10, 20, 50, and 100 neighbouring cells. Each neighbourhood scale was treated as a separate prediction task and trained and evaluated independently. The radii used for each dataset are reported in Supplementary Table 5.

##### Task definition and evaluation

For niche composition prediction, each model was trained to regress the target composition vector *N*_*i*_ ∈ [0, 1]^*L*^ for each cell *i*. The *L* output scores were transformed using the softplus function to ensure non-negativity and then normalised by their sum, yielding a predicted composition vector 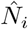 whose entries sum to one. Models were optimised using the element-wise mean squared error between the predicted and target compositions:

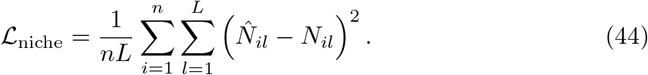

MSE was used for model optimisation and checkpoint selection, while MAE was reported as the primary evaluation metric.

##### Split-half reliability of niche-composition targets

To assess the sampling stability of niche-composition targets at each neighbourhood radius, we performed a model-free split-half reliability analysis. For each centre cell, its neighbours were randomly assigned with equal probability to two non-overlapping subsets, from which two cell-type composition estimates, **a**_*i*_ and **b**_*i*_, were calculated. Centre cells for which either subset was empty were excluded from that split. Reliability was calculated separately for each centre-cell type. To remove centre-cell-type-specific average niche compositions and FOV-level composition differences, each composition channel was mean-centred within each combination of centre-cell type and FOV:

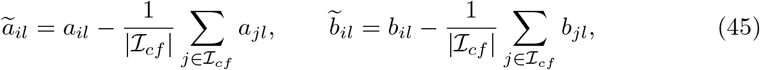

where *a*_*il*_ and *b*_*il*_ denote the estimated proportions of neighbour cell type *l* for centre cell *i*, and *ℐ*_*cf*_ denotes the set of valid centre cells of type *c* in FOV *f*. The centred estimates were then pooled across FOVs. Reliability for centre-cell type *c* at radius *r* was defined as

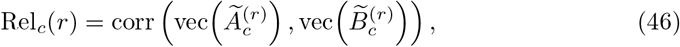

where 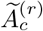 and 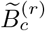 contain the centred composition estimates for centre-cell type *c*, and vec(*·*) denotes matrix vectorisation. The correlation was therefore computed jointly across centre cells and neighbour-cell-type channels. The random split was repeated five times, and reliability was averaged across repeats. Centre-cell types represented by fewer than 50 valid cells in a split were excluded.

##### Cell-identity-controlled niche *R*^2^

To quantify the niche variation recovered beyond cell identity, we computed a cell-identity-controlled coefficient of determination separately at each neighbourhood radius. For each centre-cell type *c*, let *ℐ*_*c*_ denote the corresponding set of cells and let

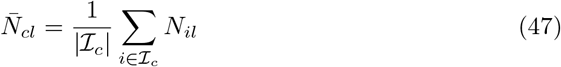

denote the mean proportion of neighbour cell type *l*. For each method, predictions from three independently trained models with different random seeds were first averaged to obtain 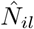, and 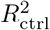 was then computed once using these averaged predictions.

We defined

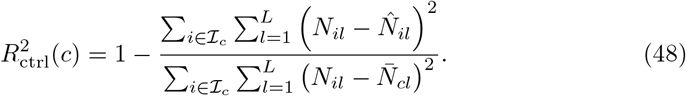

The same prediction-averaging procedure was applied consistently to all compared methods. Using the centre-cell-type-specific mean as the baseline removes between-type differences and isolates within-type niche variation. Values above zero indicate recovery beyond this baseline, whereas values below zero indicate worse performance than the baseline. Centre-cell types with fewer than 50 cells were excluded.

##### Tumour-neighbour separability

For each focal cell, the tumour-neighbour fraction was defined as the proportion of neighbouring cells annotated as tumour cells. Within each focal cell type, cells were labelled as tumour-poor if the ground-truth fraction was below 0.5 and tumour-rich otherwise. Separability was quantified by AUROC, using the model-predicted tumour-neighbour fraction as the score and the ground-truth tumour-poor/tumour-rich assignment as the binary label. AUROC was computed separately for each focal cell type and neighbourhood radius. For visualisation, the distributions of tumour-neighbour fractions were smoothed using Gaussian kernel density estimation; AUROC was computed from the unsmoothed per-cell values.

##### Neighbourhood density prediction

As a complementary probe of local tissue organisation, we predicted neighbourhood density, defined as the number of neighbouring cells within a given spatial radius of each cell. This captures local cellular packing rather than cell-type composition. Using the same radius-graph construction as for niche composition, graphs were constructed independently within each FOV, with self-loops excluded. For a radius *r*, the density target for cell *i* was defined as the raw neighbour count

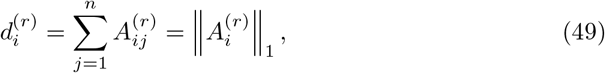

where *A*^(*r*)^ *∈ {*0, 1*}*^*n×n*^ denotes the binary radius-graph adjacency matrix.

Three dataset-specific radii were evaluated, with each radius defining a separate prediction task that was trained and evaluated independently. The radii used for each dataset are reported in Supplementary Table 6. A linear prediction head mapped the per-cell representation **h**_*i*_ to a scalar density prediction 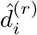. Models were optimised using mean squared error, with validation MSE used for model selection, and the coefficient of determination (*R*^2^) was reported as the primary evaluation metric.

### 4.10 Attention analysis

#### Attention extraction

For each evaluation dataset, we analysed all cells from five prespecified validation FOVs using the pretrained NexuST encoder (Supplementary Table 10). Cells were exhaustively partitioned into non-overlapping, size-balanced Hilbert chunks of at most 512 cells, and cell-level attention was evaluated only among cells belonging to the same chunk. The encoder was placed in evaluation mode.

For each gene-level and cell-level Transformer layer in hierarchical block *s*, attention was reconstructed separately for each head and then averaged:

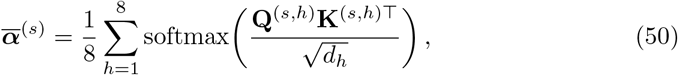

where *h* indexes the attention head and *d*_*h*_ is the head dimension. For consistency with the figures, hierarchical blocks *s* = 1, …, 8 are reported as layers L0–L7. Padded gene keys were masked before the softmax, whereas measured genes with zero expression remained valid keys. Cell-to-gene attention was defined as attention from a cell token to the gene tokens associated with the same cell. Cell-to-cell attention was defined as attention from one cell token to the cell tokens of cells in the same Hilbert chunk.

#### Gene-level attention entropy

Gene-level analyses were performed using the pretrained NexuST encoder. For each dataset, hierarchical block *s*, and cell type *c*, we pooled cell-to-gene attention over all cells of that type across the five selected validation FOVs. Specifically, if 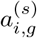 denotes the head-averaged attention from the cell token of cell *i* to gene token *g*, the pooled cell-type profile was

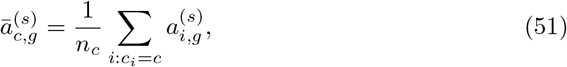

where *n*_*c*_ is the total number of cells of type *c* across the selected FOVs. Thus, FOVs contributed in proportion to their number of eligible cells rather than through an equal-weight average of FOV-level estimates.

Because the cell token could also attend to its own cell-token key and the organ-token key, the pooled cell-type profile was conditioned on the total attention assigned to the *G* = 300 gene-token keys:

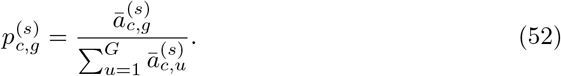

We then calculated the normalised Shannon entropy of each pooled cell-type profile and macro-averaged it across the cell types observed in that dataset:

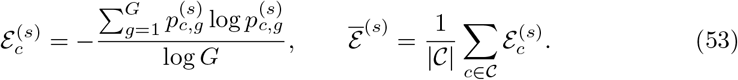

This normalisation gives 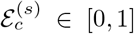, with lower values indicating attention concentrated on a smaller subset of genes and higher values indicating a more broadly distributed gene-attention profile.

**Cell-to-gene attention and expression heatmaps**

Cell-to-gene attention heatmaps were generated from the pooled, gene-conditioned profiles 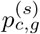 of the pretrained NexuST encoder evaluated on the Xenium lung fibrosis dataset. For visualisation, we used a fixed panel of 17 cell types spanning epithelial, fibroblast/stromal and myeloid populations, together with 23 genes covering extracellular-matrix, epithelial-remodelling, fibroblast-activation and myeloid programmes. The 23 displayed genes were selected after conditioning attention on the complete set of *G* = 300 gene tokens and were not renormalised as a display subset. The same cell types, genes, ordering and fixed logarithmic colour scale were used for the L2-versus-L7 comparison and the complete L0–L7 display.

To compare attention with the focal cells’ transcriptional programmes, we also calculated mean expression for the same 17 cell types and 23 genes using the same five Xenium lung fibrosis validation FOVs. Expression values were taken from the same library-size-normalised and log1p-transformed matrix used as model input and averaged across all eligible cells of each annotated cell type. The expression heatmap used the same cell-type and gene ordering as the attention heatmaps.

#### Abundance-normalised cell-to-cell attention

Using the pretrained encoder, for each cell layer *s*, source cell type *c* and target cell type *t*, we pooled head-averaged attention over all eligible ordered cell pairs within the same Hilbert chunk. Let *J*_*i*_ denote the cells in the chunk containing source cell *i*. After excluding the exact self-pair *j* = *i*, the mean attention per available source–target pair was

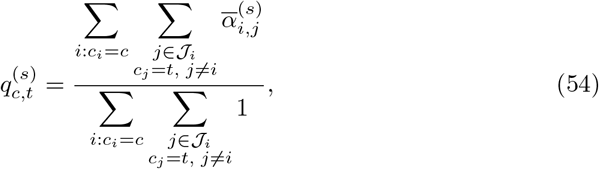

with 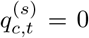 when no such pair was available. The abundance-normalised cell-type attention matrix was then obtained by normalising each source-cell-type row across the complete set of observed target cell types,

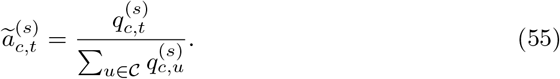

Attention mass and available-pair counts were pooled across all Hilbert chunks and validation FOVs before normalisation. Complete Xenium matrices included all 46 cell types observed in the selected validation FOVs and were ordered according to the six analysis lineages defined in Supplementary Table 12. Focused displays used the same fixed 17-cell-type panel as the cell-to-gene heatmaps, extracted only after normalisation over all 46 target cell types and without renormalising the displayed subset.

#### Within-lineage attention

Using the broad-lineage assignments defined in Supplementary Table 12 and the corresponding dataset-specific lineage mappings, within-lineage attention for source cell type *c* at layer *s* was defined as

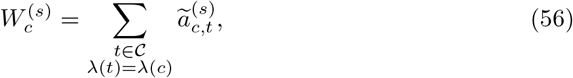

where *λ*(*c*) denotes the broad lineage of cell type *c*. Exact self-cell pairs had already been excluded, whereas attention to other cells of the same annotated cell type was retained. Dataset-level within-lineage attention was then obtained by macro-averaging across source cell types:

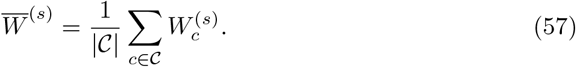

### 4.11 Benchmarking methods

In this section, we describe the implementation details for all benchmarking methods. All pretrained baselines were adapted to the same downstream benchmark protocol described above in Section 4.8. Because the models differ in gene identifier systems and fixed vocabularies, the shared HVG300 set was first converted to each model’s expected gene identifier format and then intersected with the corresponding model vocabulary before model-specific input construction. Nicheformer and CellPLM use Ensembl gene IDs, whereas NexuST and scGPT-spatial use gene symbols. The resulting gene coverage is reported in Supplementary Table 7, with most model–dataset combinations retaining over 90% of the shared HVG300 set. For the held-out gene recovery task, target genes were selected under the unified task protocol after harmonizing gene identifiers across models. The candidate target set was restricted to genes present in the pretrained vocabularies of all compared models, including NexuST, Nicheformer, scGPT-spatial and CellPLM. This ensured that gene withholding and evaluation were shared across methods, rather than being determined separately for each model. For all pretrained baselines, encoder-derived cell representations were passed to the shared task-specific linear heads defined in the downstream evaluation protocol. As linear probing used the same training settings across methods, the following model-specific sections focus on input construction, representation extraction and fine-tuning settings unique to each baseline.

#### NexuST

For NexuST, the shared HVG300 set was represented using HGNC gene symbols and processed using the same expression preprocessing pipeline as during pretraining. Because the number of selected genes matched the model input length, *M* = 300, all HVG300 genes were provided to the model for every cell. Genes with zero expression were retained as valid zero-valued inputs rather than being removed from the input sequence.

During training, for both linear probing and fine-tuning, cells were organised into spatial patches of *N* = 512 cells using the same spatial sampling procedure as during pretraining, described in Section 4.2. The updated cell token produced by the final hierarchical encoder block was used as the cell representation for downstream prediction. The pretrained reconstruction decoder was not used, and the resulting cell representation was passed directly to the shared task-specific linear head. During validation, both linear probing and fine-tuning used the deterministic whole-slide inference strategy described in Section 4.6.

As described above, linear probing used identical optimisation settings across all methods, whereas fine-tuning hyperparameters were model specific because of differences in architecture and memory requirements. For NexuST fine-tuning, the effective global batch size was fixed at 64 spatial patches for all downstream tasks. Task-specific learning rates and other optimisation settings are reported in Supplementary Table 9.

#### Nicheformer

For the comparison with Nicheformer, we used its latest official GitHub repository. To align with its data infrastructure, gene symbols were mapped to its fixed vocabulary of 20,310 Ensembl genes, and input data were stored in parquet format rather than H5AD. Following the original implementation, for each cell, non-zero genes from the vocabulary-compatible HVG set were selected and sorted in descending order according to normalised counts. This rank-ordered gene list was used as the input sequence, with positional encodings providing discretised expression-rank information to the model. For representation extraction, we followed the strategy used in the original Nicheformer paper. Unlike NexuST, which uses an explicit cell token to obtain the cell embedding, Nicheformer obtains the cell embedding by pooling gene-token embeddings from the last transformer layer. We therefore used only gene tokens and excluded meta tokens such as platform information. For fine-tuning, although the default number of fine-tuning epochs in the original repository and paper is 1, we found that one epoch was insufficient for our downstream tasks. We therefore fine-tuned Nicheformer under the shared training budget described above, using a maximum of 50 epochs with early stopping. The global batch size was set to 1,024 across eight GPUs, corresponding to a per-GPU batch size of 128, and the learning rate was set to 1 *×* 10^−4^ for all tasks.

#### scGPT-spatial

For the comparison with scGPT-spatial, we used its original fixed gene-symbol vocabulary containing 60,697 genes. Unlike Nicheformer, scGPT-spatial does not convert expression values into a rank-ordered gene sequence. Instead, following the original model design, gene identities and expression values were separately embedded and then combined as the model input. Similar to NexuST, scGPT-spatial uses a [CLS] token as the cell embedding, which was used as the cell representation for downstream prediction. For fine-tuning, because the original scGPT-spatial repository does not provide task-specific fine-tuning code or hyperparameters for our downstream benchmarks, we implemented the fine-tuning procedure for these tasks. We used a global batch size of 512, which was the largest stable batch size under our hardware setting, and a learning rate of 1 *×* 10^−4^.

#### CellPLM

For the comparison with CellPLM, gene symbols were mapped to Ensembl gene IDs and aligned to its fixed vocabulary of 19,374 genes. Unlike Nicheformer and scGPT-spatial, which tokenize genes within each cell and apply attention across genes, CellPLM constructs a cell-level token by aligning each cell’s expression profile to its fixed gene vocabulary and multiplying this expression vector with a learnable gene-embedding matrix. Equivalently, each cell token is obtained as an expression-weighted sum of per-gene embedding vectors, so genes with zero expression do not contribute to the aggregation. The resulting cell tokens are then processed by a transformer that attends across cells within a tissue. Following the original CellPLM design, the vocabulary-compatible HVG features were placed into the fixed CellPLM gene space, with unmatched or unobserved genes contributing zero. We used the CellPLM encoder together with its latent module to produce the cell representation for downstream prediction. For fine-tuning, CellPLM was trained according to its cell-level modelling design, where each cell is first aggregated into a single cell token and attention is then applied across cells within a tissue. Fine-tuning was therefore performed on a single GPU, with each optimisation step processing one field of view (FOV) rather than a fixed-size mini-batch. Cells within each FOV were internally chunked into groups of at most 70,000 for memory efficiency. To remain consistent with the original CellPLM implementation, task-specific learning rates were used; the values are listed in Supplementary Table 9.

#### PCA

As a no-pretraining reference, we included a principal component analysis (PCA) baseline that operates purely on gene expression. Each cell was represented only by its preprocessed expression vector, and no spatial information, including cell coordinates or neighbourhood-derived features, was provided to the model. Because PCA has no learnable encoder, it was evaluated only under the linear-probing setting. For each dataset, PCA with 50 components was fitted on the training split using random seed 42 and then used to project both training and validation cells. The resulting 50-dimensional representation was treated as a frozen cell embedding and evaluated under the shared linear-probing protocol. For the held-out gene recovery task, held-out genes were removed before fitting PCA and before every projection, ensuring that PCA never observed the target genes and that both fitting and inference operated on the same reduced gene panel.

### 4.12 Architectural ablations

We conducted ablation studies to assess the key design choices in NexuST. To reduce computational cost while preserving controlled comparisons, all ablation experiments used the same reduced-size configuration, with an embedding dimension of 384, six hierarchical blocks and six attention heads. The matched ablation baseline retained alternating gene- and cell-level attention, an eight-expert mixture-of-experts decoder and a library-size prediction head.

To examine the encoder design, we evaluated three variants. First, a gene-only encoder replaced the cell-level attention layers with additional gene-level layers while preserving the total network depth, testing the contribution of intercellular communication. Second, a stacked end-to-end encoder applied all gene-level layers before all cell-level layers, testing the contribution of repeated gene–cell information exchange.

Third, a two-stage encoder pretrained the gene-level stack before freezing it and optimising the cell-level stack, allowing us to compare joint end-to-end learning with sequential optimisation.

To examine the decoder design, we replaced the mixture-of-experts decoder with a single masked-value decoder or removed the library-size prediction head, thereby testing expert specialisation and the factorisation of expression into gene composition and library size, respectively. Finally, we included a shuffled-sampling control that disrupted spatially coherent patch membership while preserving the selected cells, the number and size of patches, and the exposure frequency of each cell.

All variants were trained on the same 25% subset of the pretraining corpus, constructed by selecting complete subslides from every slide until the target fraction was reached. This retained local cell density and neighbourhood structure within the selected regions while preserving the diversity of organs, experimental platforms and biological conditions in the full corpus. The matched baseline and all single-stage variants were pretrained for 2,500 steps, whereas the two-stage variant was trained for 2,500 steps in each stage, for a total of 5,000 steps. Apart from the component or training strategy under investigation, all settings were held fixed across the ablation variants.

Evaluation followed the same frozen-encoder linear-probing protocol, hyperparameters, three random seeds and validation-based early stopping described above. Cell-type annotation, region prediction and niche-composition prediction were evaluated on all datasets for which the corresponding labels were available. For niche-composition prediction, only the largest neighbourhood radius, *r*_3_, was evaluated.

### 4.13 Full-panel evaluation

The primary benchmark used a common dataset-specific HVG300 panel across models to standardise input length and gene content, ensuring that performance differences were not driven by unequal feature sets. However, this controlled setting may not fully reflect the capacity of models that can process longer gene sequences. We therefore performed a complementary full-panel evaluation in which highly variable gene selection was removed and each foundation model was allowed to use the complete measured gene panel supported by its pretrained vocabulary and native input formulation.

The measured panels contained 300 genes for MERFISH human brain, 999 genes for CosMx liver normal and liver cancer, and 342 genes for Xenium lung fibrosis. Gene identifiers were aligned to each foundation model’s pretrained vocabulary, after which all compatible measured genes were provided using the model’s native input-encoding procedure. The maximum input length was set to 1,000 genes for sequence-based models, and none of the evaluated panels exceeded this limit. For the non-pretrained expression-only baseline, PCA with 50 components (PCA-50) was fitted on the full measured panel of the training split and then used to project the validation cells. Except for replacing the HVG300 input with the full measured panel, we used the same preprocessing, training and validation splits, linear-probing protocol, optimisation hyperparameters, three random seeds and validation-based early stopping as in the primary benchmark. Cell-type annotation was evaluated on all four datasets, region prediction on the three datasets with region annotations, and niche-composition prediction on all four datasets at the largest neighbourhood radius, *r*_3_ (Supplementary Note 5 and Supplementary Table 15).

## Data availability

The publicly available datasets used in this study are described in the Methods, and the sources of the HumanST-46M pretraining corpus are listed in Supplementary Table 1. The processed data will be made available on Hugging Face at https://huggingface.co/datasets/Haiping-UoM/HumanST-46M.

## Code availability

The NexuST training, inference and evaluation code will be made available on GitHub at https://github.com/UoM-HealthAI/NexuST. Pretrained model weights will be made available on Hugging Face at https://huggingface.co/Haiping-UoM/NexuST.

## Acknowledgements

We acknowledge the EuroHPC Joint Undertaking for awarding us access to Vega hosted by IZUM, Slovenia, and Leonardo hosted by CINECA, Italy, under project EHPC-DEV-2025D11-121, and to LUMI hosted by CSC, Finland, under project EHPC-DEV-2026D04-068.

This work is also supported by funds from the Cancer Research UK (Ref: PRCBTP-Nov24/100012).

## Author contributions

Haiping Liu collected and curated the data, designed and implemented the model, conducted model training and analyses, visualised the results and drafted the manuscript. Qian Zhao contributed to data collection, analysis and manuscript writing. Lijing Lin contributed to data analysis and critically reviewed the manuscript. Zhiyong Zou contributed to data analysis. Wenhao Cai contributed to visualisation. Jingyuan Sun and Yuxi Zhou contributed to data analysis and critically reviewed the manuscript. Mauricio A. Alvarez, Andrew Gilmore, Magnus Rattray and Alejandro F. Frangi critically reviewed the manuscript. Hongpeng Zhou supervised the study.

