## Supplementary Information for "NexuST: A Hierarchical Foundation Model for Spatial Transcriptomics"

### Supplementary Notes

#### Supplementary Note 1: The library-size shortcut of MSE reconstruction

We provide a formal explanation for why direct MSE reconstruction of absolute expression can encourage a library-size shortcut. For notational simplicity, we suppress the cell index and consider one gene $g$. We write absolute expression as

$$x_g = \ell \rho_g,$$

where $\ell = \sum_h x_h$ is the cell-specific library size and $\rho_g = x_g/\ell$ is the relative composition of gene $g$, with $\sum_g \rho_g = 1$. Let $\bar{\rho}_g = \mathbb{E}[\rho_g]$.

Under a squared-error objective, the optimal prediction given an information set $\mathcal{I}$ is the conditional expectation,

$$\hat{x}_g^\star(\mathcal{I}) = \mathbb{E}[x_g \mid \mathcal{I}].$$

A decoder that has learned the library size $\ell$, but carries little cell-specific information about the composition $\rho_g$, effectively has access to the information set $\mathcal{I} = \{\ell\}$. Its optimal prediction is therefore

$$\mathbb{E}[x_g \mid \ell] = \ell\, \mathbb{E}[\rho_g \mid \ell].$$

If the conditional mean composition varies weakly with library size, or under the simplifying approximation $\rho_g \perp \ell$, this becomes

$$\mathbb{E}[x_g \mid \ell] \approx \ell\, \bar{\rho}_g.$$

Thus, a decoder can reduce MSE by predicting a library-size-scaled average composition,

$$\hat{x}_g = \ell\, \bar{\rho}_g,$$

without encoding cell-specific compositional information.

The amount of error removable by this shortcut follows directly from the law of total variance:

$$\mathrm{Var}(x_g) = \underbrace{\mathrm{Var}(\mathbb{E}[x_g \mid \ell])}_{\text{explained by library size}} + \underbrace{\mathbb{E}[\mathrm{Var}(x_g \mid \ell)]}_{\text{residual at fixed library size}}.$$

A constant predictor achieves the minimal MSE

$$\min_c \mathbb{E}\big[(x_g - c)^2\big] = \mathrm{Var}(x_g),$$

whereas the optimal predictor that uses only $\ell$ achieves

$$\mathbb{E}\big[(x_g - \mathbb{E}[x_g \mid \ell])^2\big] = \mathbb{E}[\mathrm{Var}(x_g \mid \ell)]\,.$$

Therefore, the MSE reduction obtained from knowing only the library size is exactly

$$\mathrm{Var}(x_g) - \mathbb{E}[\mathrm{Var}(x_g \mid \ell)] = \mathrm{Var}(\mathbb{E}[x_g \mid \ell])\,.$$

This reduction does not require the decoder to learn the cell-specific composition $\rho_g$; it only requires the decoder to capture the dependence of absolute expression on $\ell$.

Using the factorisation $x_g = \ell\rho_g$, the exact decomposition can be written as

$$\mathrm{Var}(x_g) = \mathrm{Var}(\ell\, m_g(\ell)) + \mathbb{E}\left[\ell^2\, \mathrm{Var}(\rho_g \mid \ell)\right],$$

where $m_g(\ell) = \mathbb{E}[\rho_g \mid \ell]$. Under the simplifying approximation that $\rho_g$ is independent of $\ell$, this reduces to

$$\mathrm{Var}(x_g) \approx \underbrace{\bar{\rho}_g^2\, \mathrm{Var}(\ell)}_{\text{removable by fitting } \ell} + \underbrace{\mathbb{E}[\ell^2]\, \mathrm{Var}(\rho_g)}_{\text{requires cell-specific composition}}.$$

The first term can be removed by fitting the cell-wise scale alone, while the second term requires modelling the cell-specific composition. Since library sizes in spatial transcriptomics can vary substantially across cells, the library-size term may explain a large fraction of the MSE reduction available during reconstruction. This creates a shortcut for a direct absolute-expression MSE objective: the decoder can substantially lower the loss by recovering $\ell$, while leaving $\rho_g$ close to its population average.

This motivates separating the reconstruction of cell-wise scale and gene-wise composition in the decoder. By modelling library size explicitly, the decoder reduces the incentive for the cell tokens to encode only global expression scale, and places greater pressure on them to preserve composition-related information that is more informative of cell state and tissue context.

#### Supplementary Note 2: NexuST training pseudocode

**Algorithm 1** NexuST training pseudocode

```
# x: gene values, ids: gene index, p: platform ID
# Padding positions are excluded from attention and softmax.
mask = build_mask(ids, ratio=0.2)

# 1. Embedding & Tokenization
# ValEmb handles masking internally
H = GeneEmb(ids) + ValEmb(x, mask) # [BN, G, D]
H = cat([OrganEmb(organ), H], dim=1)

# cls_token: [1, 1, D] -> [BN, 1, D]
c = cls_token.expand(BN, 1, D) + PE(coords)

# 2. Hierarchical Encoder
for i in range(num_blocks):
    H, c = GeneLayer(H, c)
    c = c.view(B, N, D)
    c = CellLayer(c).view(BN, 1, D)

# 3. MoE Decoder
# Add platform information
z = cat([c.view(BN, D), PlatformEmb(p)], dim=-1)

# Query: gene embeddings, Context: cell tokens
logits = MoE(query=GeneEmb(ids), context=z)
lib = Softplus(LibHead(z))

# Loss on masked positions
rec = log1p(lib * Softmax(logits))
loss = MSE(rec, x, mask)
loss.backward()
```

PE: Positional Encoding; BN: Batch size × Number of cells.

#### Supplementary Note 3: Hilbert Curve

A Hilbert curve is a space-filling curve that recursively visits all locations in a two-dimensional grid and assigns each location a one-dimensional order index. It provides a way to map two-dimensional coordinates to a scalar value:

$$(x_i, y_i) \longrightarrow h_i,$$

where $(x_i, y_i)$ denotes the spatial coordinate of cell $i$, and $h_i$ is its corresponding Hilbert index. In practice, the continuous spatial coordinates are first normalized and discretized onto a finite two-dimensional grid. The Hilbert index is then computed for each discretized grid location, and cells can be sorted according to this index to obtain a one-dimensional sequence.

The main property of the Hilbert curve is locality preservation. Compared with simple row-wise or column-wise ordering, nearby points along the Hilbert order are more likely to remain close in the original two-dimensional space. This makes Hilbert ordering suitable for converting spatial point sets into sequences while reducing the disruption of local neighborhoods. However, the mapping is not perfectly distance-preserving: two cells that are close in two-dimensional space may still be separated along the one-dimensional Hilbert order, and vice versa. Therefore, Hilbert ordering should be understood as a practical locality-preserving linearization, rather than an exact representation of two-dimensional tissue geometry.

In this work, Hilbert ordering is used only to construct fixed-size inference segments from whole-slide cell coordinates. It does not replace the model's spatial encoding or introduce an additional biological assumption.

#### Supplementary Note 4: Ablation analysis of NexuST design choices

We assessed the ablation variants across 11 task–dataset combinations spanning cell-type annotation, region prediction and niche-composition prediction (Supplementary Tables 13 and 14). Because these tasks use metrics with different absolute scales, we compared the variants using their within-dataset ranks. The complete NexuST design achieved the lowest overall mean rank of 1.91, followed by the stacked end-to-end variant at 2.36. The complete design ranked first for region prediction and jointly first for niche-composition prediction, while remaining competitive for cell-type annotation. Its principal advantage was therefore consistent performance across tasks and datasets rather than uniformly achieving the best result in every individual setting.

The encoder ablations revealed a clear distinction between cell-intrinsic and spatially contextual tasks. The gene-only encoder achieved a slightly better mean rank than the baseline for cell-type annotation (2.25 versus 2.50), which is expected because cell identity is determined predominantly by cell-intrinsic gene expression. In contrast, it ranked last for both region prediction and niche-composition prediction, with a mean rank of 7.00 for each task. This marked decline on the two spatial tasks demonstrates the importance of hierarchical cell-level modelling for capturing tissue context beyond the expression profile of an individual cell. Under the same encoder depth, the stacked end-to-end variant remained competitive for cell-type annotation but performed less well than the interleaved baseline for region prediction (2.33 versus 1.33) and niche-composition prediction (2.75 versus 1.75). Alternating gene- and cell-level layers gives the two representations repeated opportunities for mutual refinement: updated cell-context representations can inform subsequent gene-level processing, which in turn updates the representations used in the next round of cell-level communication. The stronger spatial performance of the baseline therefore supports repeated gene–cell interaction over a single sequential transition from gene-level to cell-level processing. The two-stage variant introduced a more complex optimization procedure, training the gene stack for 2,500 steps before freezing it and training the cell stack for a further 2,500 steps. Although this strategy achieved the same mean rank as the baseline for niche-composition prediction, it remained below the baseline for region prediction (3.33 versus 1.33) and substantially reduced cell-type annotation performance (5.25 versus 2.50). Thus, even with matched encoder depth and more total update steps, separating gene- and cell-level training impaired the preservation of cell identity. These results suggest that an independently trained cell stage can learn useful spatial structure, but that joint end-to-end optimization is required to integrate this structure without sacrificing cell-intrinsic information.

Cell-type-resolved annotation in CosMx liver normal further localised the benefit of interleaving (Supplementary Fig. 13). Averaged across the five hepatocyte subtypes, the matched baseline achieved an F1 of 0.574, compared with 0.527 for the stacked encoder and 0.518 after spatially coherent sampling was disrupted. The largest separation occurred for Hep 5, for which F1 decreased from 0.670 in the matched baseline to 0.570 with the stacked encoder and 0.492 with shuffled sampling. The gene-only encoder remained closer to the baseline across hepatocyte subtypes (mean F1 0.566), indicating that cell-intrinsic expression already carries substantial information about hepatocyte state, while repeated gene–cell exchange and coherent local neighbourhoods provide additional information for resolving spatially zonated states. Immune populations showed mixed responses across these variants, suggesting that this benefit is cell-state dependent rather than a uniform improvement in annotation.

The decoder ablations provided further support for the NexuST reconstruction design. Replacing the mixture-of-experts decoder with a single masked-value decoder produced an overall mean rank of 3.91 and remained competitive in individual datasets, suggesting that expert specialisation primarily improves the consistency of representation learning across heterogeneous expression patterns. More importantly, removing the library-size prediction head resulted in the worst overall mean rank among all evaluated variants (5.82). This variant ranked last for cell-type annotation (7.00) and also performed poorly for region prediction (5.00) and niche-composition prediction (5.25). These results provide direct empirical support for explicitly separating library size from gene composition during pretraining, consistent with the library-size shortcut derived in Supplementary Note 1. Because the reconstruction decoder was discarded before downstream evaluation and all variants used the same frozen-encoder linear probes, the performance decline indicates that this factorisation changes the biological information retained in the pretrained cell embeddings, rather than merely improving the capacity of the reconstruction decoder.

The spatial-sampling ablation tested whether the hierarchical encoder benefited merely from processing multiple cells together or specifically from their organisation into local tissue neighbourhoods. Randomising KNN-based patch membership reduced performance relative to the baseline across all 11 task–dataset combinations, resulting in an overall mean rank of 5.27, with mean ranks of 4.75, 5.33 and 5.75 for cell-type annotation, region prediction and niche-composition prediction, respectively. Because this control preserved the selected cells, the number and size of patches, and the exposure of each cell during pretraining, the decline cannot be explained by differences in data quantity or sampling frequency. Instead, it isolates the loss of spatially coherent patch membership: cell-level attention is applied to cells that no longer form a genuine local microenvironment, preventing the encoder from learning recurring relationships between cell state and tissue context. The reduction in region- and niche-composition performance directly supports the importance of local neighbourhood structure for spatial representation learning. The accompanying decline in cell-type annotation further indicates that spatial context can refine cell identity, even when the primary signal for annotation remains cell-intrinsic gene expression. This control does not establish KNN-based patching as the uniquely optimal neighbourhood-construction strategy, but demonstrates that preserving spatially coherent neighbourhoods during pretraining is important for the transferable representations learned by NexuST.

#### Supplementary Note 5: Scalability to expanded gene panels

The primary benchmark used a common dataset-specific HVG300 panel to standardise input length and gene content across models. Although this design provides a controlled comparison, it does not fully assess whether a model can accommodate broader gene panels when additional measured genes are available. We therefore performed a complementary full-panel evaluation using each foundation model's complete vocabulary-compatible measured gene panel and native input formulation, together with an expression-only PCA-50 baseline fitted on the full measured panel. This expanded the input to 999 genes in the two CosMx datasets and 342 genes in Xenium lung fibrosis, while MERFISH human brain contained 300 measured genes. We evaluated the resulting representations on cell-type annotation, region prediction and niche-composition prediction.

Across the 11 task–dataset combinations and all five methods, NexuST achieved the best performance in 10 (Supplementary Table 15). For cell-type annotation, NexuST ranked first on MERFISH human brain and both CosMx liver datasets, giving the lowest mean rank of 1.25; Nicheformer performed better only on Xenium lung fibrosis. PCA-50 remained a strong expression-only reference, ranking second on MERFISH human brain and third overall for annotation. NexuST ranked first on all three region-prediction datasets and achieved the lowest niche-composition prediction error on all four datasets at the largest neighbourhood radius, giving a mean rank of 1.00 for both tasks. Its strong performance on every task for the two 999-gene CosMx panels indicates that the advantage was maintained when the input was expanded substantially beyond the controlled HVG300 setting.

Together, these results demonstrate that NexuST is not tied to a fixed HVG300 feature set. Its gene-token formulation enables the pretrained encoder to accommodate broader vocabulary-compatible gene panels—up to 999 genes in this evaluation—without additional pretraining or encoder adaptation, while preserving strong transferable representations. This flexibility demonstrates the practical scalability of NexuST across spatial transcriptomics assays with increasing gene coverage.

#### Supplementary Tables

**Supplementary Table 1**: **Composition of HumanST-46M at physical-slide granularity.** One row represents one tissue slide; for h5ad files containing multiple sections, slide identifiers are written as *file::section*. Slides are grouped by platform and source. The "Cells $\leq$ 300 (%)" column reports the percentage of cells in which no more than 300 genes have non-zero expression after QC and gene-vocabulary filtering; across the full pretraining corpus, this proportion is 92.1%. The "Sparsity (%)" column reports the percentage of zero cell–gene entries in the corresponding expression matrix for each slide.

| Slide | Platform | Organ | Source | Cells | Panel | Cells $\leq$ 300 (%) | Sparsity (%) |
|---|---|---|---|---|---|---|---|
| HuBrain | MERFISH | brain | Vizgen MERSCOPE | 153,301 | 815 | 80.23 | 74.14 |
| HuBrain_1K_Alzheimer | MERFISH | brain | Vizgen MERSCOPE | 137,627 | 960 | 97.39 | 87.61 |
| HubcTMA_R1 | MERFISH | breast | Vizgen MERSCOPE | 92,461 | 815 | 98.99 | 83.79 |
| HubcTMA_R2 | MERFISH | breast | Vizgen MERSCOPE | 48,900 | 815 | 99.97 | 89.02 |
| HubcTMA_R3 | MERFISH | breast | Vizgen MERSCOPE | 54,279 | 815 | 100.00 | 94.60 |
| HubcTMA_R4 | MERFISH | breast | Vizgen MERSCOPE | 72,219 | 815 | 99.99 | 88.56 |
| HubcTMA_R5 | MERFISH | breast | Vizgen MERSCOPE | 79,268 | 815 | 100.00 | 91.40 |
| HumanBreastCancerPatient1 | MERFISH | breast | Vizgen MERSCOPE | 713,121 | 500 | 99.70 | 75.28 |
| HuColonCan | MERFISH | colon | Vizgen MERSCOPE | 703,879 | 815 | 99.91 | 90.38 |
| HumanColonCancerPatient1 | MERFISH | colon | Vizgen MERSCOPE | 677,451 | 500 | 99.99 | 77.17 |
| HumanColonCancerPatient2 | MERFISH | colon | Vizgen MERSCOPE | 817,588 | 500 | 99.99 | 78.48 |
| HumanLiverCancerPatient1 | MERFISH | liver | Vizgen MERSCOPE | 568,355 | 500 | 99.99 | 81.31 |
| HumanLiverCancerPatient2 | MERFISH | liver | Vizgen MERSCOPE | 598,141 | 500 | 100.00 | 81.76 |
| HumanLungCancerPatient1 | MERFISH | lung | Vizgen MERSCOPE | 353,762 | 500 | 100.00 | 83.32 |
| HumanLungCancerPatient2 | MERFISH | lung | Vizgen MERSCOPE | 836,739 | 500 | 99.95 | 76.65 |
| HumanOvarianCancerPatient1 | MERFISH | ovary | Vizgen MERSCOPE | 358,485 | 500 | 99.99 | 76.49 |
| HumanOvarianCancerPatient2Slice1 | MERFISH | ovary | Vizgen MERSCOPE | 254,347 | 500 | 100.00 | 79.38 |
| HumanOvarianCancerPatient2Slice2 | MERFISH | ovary | Vizgen MERSCOPE | 71,381 | 500 | 100.00 | 79.51 |
| HumanOvarianCancerPatient2Slice3 | MERFISH | ovary | Vizgen MERSCOPE | 212,425 | 500 | 99.99 | 76.85 |
| HumanProstateCancerPatient1 | MERFISH | prostate | Vizgen MERSCOPE | 721,668 | 500 | 100.00 | 82.96 |
| HumanProstateCancerPatient2 | MERFISH | prostate | Vizgen MERSCOPE | 993,825 | 500 | 100.00 | 86.08 |
| HumanMelanomaPatient1 | MERFISH | skin | Vizgen MERSCOPE | 468,138 | 500 | 100.00 | 83.54 |
| HumanMelanomaPatient2 | MERFISH | skin | Vizgen MERSCOPE | 207,869 | 500 | 100.00 | 82.33 |
| HumanUterineCancerPatient1 | MERFISH | uterus | Vizgen MERSCOPE | 843,285 | 500 | 100.00 | 78.14 |

*(continued on next page)*

(continued from previous page)

| Slide | Platform | Organ | Source | Cells | Panel | Cells ≤ 300 (%) | Sparsity (%) |
|---|---|---|---|---|---|---|---|
| HumanUterineCancerPatient2-RACostain | MERFISH | uterus | Vizgen MERSCOPE | 741,749 | 500 | 100.00 | 75.11 |
| HumanUterineCancerPatient2-ROCostain | MERFISH | uterus | Vizgen MERSCOPE | 758,506 | 500 | 99.98 | 73.19 |
| GSE284005_merfish_all::ms10r0 | MERFISH | brain | Publication [40] | 35,559 | 500 | 100.00 | 96.43 |
| GSE284005_merfish_all::ms11r1 | MERFISH | brain | Publication [40] | 34,952 | 500 | 100.00 | 95.87 |
| GSE284005_merfish_all::ms12r0 | MERFISH | brain | Publication [40] | 10,814 | 500 | 100.00 | 96.60 |
| GSE284005_merfish_all::ms12r1 | MERFISH | brain | Publication [40] | 5,550 | 500 | 100.00 | 96.43 |
| GSE284005_merfish_all::ms15r0 | MERFISH | brain | Publication [40] | 22,006 | 500 | 100.00 | 97.12 |
| GSE284005_merfish_all::ms15r1 | MERFISH | brain | Publication [40] | 24,415 | 500 | 100.00 | 96.72 |
| GSE284005_merfish_all::ms16r0 | MERFISH | brain | Publication [40] | 19,373 | 500 | 100.00 | 95.75 |
| GSE284005_merfish_all::ms16r1 | MERFISH | brain | Publication [40] | 44,000 | 500 | 100.00 | 96.29 |
| GSE284005_merfish_all::ms1r1 | MERFISH | brain | Publication [40] | 26,082 | 500 | 100.00 | 95.93 |
| GSE284005_merfish_all::ms1r2 | MERFISH | brain | Publication [40] | 31,404 | 500 | 100.00 | 95.96 |
| GSE284005_merfish_all::ms4r1 | MERFISH | brain | Publication [40] | 31,244 | 500 | 100.00 | 95.85 |
| GSE284005_merfish_all::ms4r2 | MERFISH | brain | Publication [40] | 20,149 | 500 | 100.00 | 96.58 |
| GSE284005_merfish_all::ms5r1 | MERFISH | brain | Publication [40] | 25,153 | 500 | 100.00 | 95.71 |
| GSE284005_merfish_all::ms6r0 | MERFISH | brain | Publication [40] | 10,768 | 500 | 100.00 | 95.53 |
| GSE284005_merfish_all::ms6r1 | MERFISH | brain | Publication [40] | 22,299 | 500 | 100.00 | 96.18 |
| GSE284005_merfish_all::ms7r1 | MERFISH | brain | Publication [40] | 12,134 | 500 | 100.00 | 96.75 |
| GSE284005_merfish_all::ms8r0 | MERFISH | brain | Publication [40] | 25,892 | 500 | 100.00 | 95.75 |
| gw15::UMB1117—F1a | MERFISH | brain | Publication [37] | 548,567 | 300 | 100.00 | 84.76 |
| gw15::UMB1117—F1b | MERFISH | brain | Publication [37] | 410,492 | 300 | 100.00 | 87.70 |
| gw15::UMB1117—F2a | MERFISH | brain | Publication [37] | 339,105 | 300 | 100.00 | 86.72 |
| gw15::UMB1117—F2b | MERFISH | brain | Publication [37] | 447,267 | 300 | 100.00 | 85.50 |
| gw15::UMB1117—O1 | MERFISH | brain | Publication [37] | 579,531 | 300 | 100.00 | 85.72 |
| gw15::UMB1117—P1 | MERFISH | brain | Publication [37] | 793,577 | 300 | 100.00 | 85.15 |
| gw15::UMB1117—T1 | MERFISH | brain | Publication [37] | 520,843 | 300 | 100.00 | 82.29 |
| gw15::UMB1367—O1 | MERFISH | brain | Publication [37] | 887,043 | 300 | 100.00 | 84.83 |
| gw15::UMB1367—P1 | MERFISH | brain | Publication [37] | 613,477 | 300 | 100.00 | 88.79 |
| gw18_umb1759 | MERFISH | brain | Publication [37] | 927,296 | 1,000 | 100.00 | 97.35 |
| gw20::FB080—F1 | MERFISH | brain | Publication [37] | 665,204 | 300 | 100.00 | 88.11 |
| gw20::FB080—F2a | MERFISH | brain | Publication [37] | 596,754 | 300 | 100.00 | 90.86 |
| gw20::FB080—F2b | MERFISH | brain | Publication [37] | 322,648 | 300 | 100.00 | 90.68 |
| gw20::FB080—O1a | MERFISH | brain | Publication [37] | 632,853 | 300 | 100.00 | 80.56 |

(continued on next page)

(continued from previous page)

| Slide | Platform | Organ | Source | Cells | Panel | Cells ≤ 300 (%) | Sparsity (%) |
|---|---|---|---|---|---|---|---|
| gw20::FB080—O1b | MERFISH | brain | Publication [37] | 340,393 | 300 | 100.00 | 83.77 |
| gw20::FB080—O1c | MERFISH | brain | Publication [37] | 467,082 | 300 | 100.00 | 87.78 |
| gw20::FB080—O1d | MERFISH | brain | Publication [37] | 344,168 | 300 | 100.00 | 82.86 |
| gw20::FB080—P1a | MERFISH | brain | Publication [37] | 345,889 | 300 | 100.00 | 88.20 |
| gw20::FB080—P1b | MERFISH | brain | Publication [37] | 643,168 | 300 | 100.00 | 85.79 |
| gw20::FB080—P2 | MERFISH | brain | Publication [37] | 436,856 | 300 | 100.00 | 87.16 |
| gw20::FB080—T1 | MERFISH | brain | Publication [37] | 451,806 | 300 | 100.00 | 86.23 |
| gw20::FB121—F1 | MERFISH | brain | Publication [37] | 553,738 | 300 | 100.00 | 75.87 |
| gw20::FB121—F2 | MERFISH | brain | Publication [37] | 293,593 | 300 | 100.00 | 81.25 |
| gw20::FB121—O1 | MERFISH | brain | Publication [37] | 167,000 | 300 | 100.00 | 74.69 |
| gw20::FB121—P1 | MERFISH | brain | Publication [37] | 474,743 | 300 | 100.00 | 75.14 |
| gw20::FB121—P2 | MERFISH | brain | Publication [37] | 643,601 | 300 | 100.00 | 88.31 |
| gw20::FB121—T1 | MERFISH | brain | Publication [37] | 664,950 | 300 | 100.00 | 83.54 |
| gw20_umb1031 | MERFISH | brain | Publication [37] | 677,475 | 1,000 | 100.00 | 96.70 |
| gw21_umb1932 | MERFISH | brain | Publication [37] | 571,528 | 1,000 | 100.00 | 95.01 |
| gw22::FB123—F1 | MERFISH | brain | Publication [37] | 508,417 | 300 | 100.00 | 78.91 |
| gw22::FB123—F2 | MERFISH | brain | Publication [37] | 439,986 | 300 | 100.00 | 88.17 |
| gw22::FB123—F3 | MERFISH | brain | Publication [37] | 333,113 | 300 | 100.00 | 84.35 |
| gw22::FB123—O1 | MERFISH | brain | Publication [37] | 407,591 | 300 | 100.00 | 74.15 |
| gw22::FB123—O2 | MERFISH | brain | Publication [37] | 439,060 | 300 | 100.00 | 82.48 |
| gw22::FB123—P1 | MERFISH | brain | Publication [37] | 483,951 | 300 | 100.00 | 78.91 |
| adata_healthy_diseased_merfish::AM031 | MERFISH | liver | Publication [41] | 71,056 | 317 | 100.00 | 89.05 |
| adata_healthy_diseased_merfish::AM042 | MERFISH | liver | Publication [41] | 39,188 | 317 | 100.00 | 73.40 |
| adata_healthy_diseased_merfish::AM048 | MERFISH | liver | Publication [41] | 38,366 | 317 | 100.00 | 72.46 |
| adata_healthy_diseased_merfish::AM061 | MERFISH | liver | Publication [41] | 34,066 | 317 | 100.00 | 75.88 |
| adata_healthy_diseased_merfish::AM062 | MERFISH | liver | Publication [41] | 65,894 | 317 | 100.00 | 88.06 |
| adata_healthy_diseased_merfish::AM066 | MERFISH | liver | Publication [41] | 244,691 | 317 | 100.00 | 82.86 |
| adata_healthy_diseased_merfish::AM072 | MERFISH | liver | Publication [41] | 82,450 | 317 | 100.00 | 86.50 |
| Xenium_V1_FFPE_Human_Brain_Healthy_With_Addon | Xenium | brain | 10x Xenium | 24,406 | 319 | 100.00 | 78.69 |
| Xenium_V1_Human_Brain_GBM_FFPE | Xenium | brain | 10x Xenium | 816,769 | 480 | 100.00 | 86.69 |
| Xenium_Prime_Breast_Cancer_FFPE | Xenium | breast | 10x Xenium | 699,110 | 5,090 | 94.06 | 98.36 |
| Xenium_V1_FFPE_Human_Breast_IDC | Xenium | breast | 10x Xenium | 574,852 | 280 | 100.00 | 83.43 |
| Xenium_V1_FFPE_Human_Breast_IDC_Big_1 | Xenium | breast | 10x Xenium | 892,966 | 280 | 100.00 | 82.76 |

(continued on next page)

*(continued from previous page)*

| Slide | Platform | Organ | Source | Cells | Panel | Cells ≤ 300 (%) | Sparsity (%) |
|---|---|---|---|---|---|---|---|
| Xenium_V1_FFPE_Human_Breast_IDC_Big_2 | Xenium | breast | 10x Xenium | 885,523 | 280 | 100.00 | 83.16 |
| Xenium_V1_FFPE_Human_Breast_IDC_With_Addon | Xenium | breast | 10x Xenium | 576,963 | 380 | 100.00 | 85.29 |
| Xenium_V1_FFPE_Human_Breast_ILC | Xenium | breast | 10x Xenium | 356,746 | 280 | 100.00 | 84.44 |
| Xenium_V1_Human_Colorectal_Cancer_Addon_FFPE | Xenium | colon | 10x Xenium | 388,175 | 480 | 100.00 | 90.07 |
| Xenium_V1_hColon_Cancer_Add_on_FFPE | Xenium | colon | 10x Xenium | 587,115 | 425 | 100.00 | 89.59 |
| Xenium_V1_hColon_Cancer_Base_FFPE | Xenium | colon | 10x Xenium | 647,524 | 325 | 100.00 | 89.79 |
| Xenium_V1_hColon_Non_diseased_Add_on_FFPE | Xenium | colon | 10x Xenium | 275,822 | 425 | 100.00 | 89.83 |
| Xenium_V1_hColon_Non_diseased_Base_FFPE | Xenium | colon | 10x Xenium | 270,984 | 325 | 100.00 | 89.67 |
| Xenium_V1_Human_Kidney_FFPE_Protein_updated | Xenium | kidney | 10x Xenium | 465,534 | 405 | 100.00 | 91.26 |
| Xenium_V1_hKidney_cancer_section | Xenium | kidney | 10x Xenium | 56,510 | 377 | 100.00 | 89.36 |
| Xenium_V1_hKidney_nondiseased_section | Xenium | kidney | 10x Xenium | 97,560 | 377 | 100.00 | 92.87 |
| Xenium_V1_hLiver_cancer_section_FFPE | Xenium | liver | 10x Xenium | 162,628 | 474 | 100.00 | 87.33 |
| Xenium_V1_hLiver_nondiseased_section_FFPE | Xenium | liver | 10x Xenium | 239,271 | 377 | 100.00 | 91.10 |
| Xenium_Preview_Human_Lung_Cancer_With_Add_on_2_FFPE | Xenium | lung | 10x Xenium | 531,165 | 392 | 100.00 | 83.98 |
| Xenium_Preview_Human_Non_diseased_Lung_With_Add_on_FFPE | Xenium | lung | 10x Xenium | 295,883 | 392 | 100.00 | 87.46 |
| Xenium_V1_Human_Lung_Cancer_Addon_FFPE | Xenium | lung | 10x Xenium | 161,000 | 480 | 100.00 | 84.45 |
| Xenium_V1_hLung_cancer_section | Xenium | lung | 10x Xenium | 150,365 | 377 | 100.00 | 85.78 |
| Xenium_Prime_Human_Lymph_Node_Reactive_FFPE | Xenium | lymph_node | 10x Xenium | 708,983 | 4,624 | 78.95 | 95.21 |
| Xenium_Prime_Human_Ovary_Cancer_FF | Xenium | ovary | 10x Xenium | 200,900 | 5,001 | 11.51 | 84.25 |
| Xenium_Prime_Human_Ovary_FF | Xenium | ovary | 10x Xenium | 1,157,659 | 5,001 | 11.92 | 83.48 |
| Xenium_Prime_Ovarian_Cancer_FFPE_XRrun | Xenium | ovary | 10x Xenium | 407,124 | 5,090 | 75.47 | 96.17 |
| Xenium_V1_Human_Ovary_Cancer_FF | Xenium | ovary | 10x Xenium | 205,082 | 477 | 100.00 | 76.23 |
| frontal_cortex | CosMx | brain | Bruker CosMx SMI | 188,686 | 6,077 | 32.52 | 90.86 |
| colon_wtx | CosMx | colon | Bruker CosMx SMI | 493,834 | 18,935 | 9.43 | 94.11 |
| nsclc_ffpe_Lung12 | CosMx | lung | Bruker CosMx SMI | 73,997 | 960 | 95.27 | 86.53 |
| nsclc_ffpe_Lung13 | CosMx | lung | Bruker CosMx SMI | 82,843 | 960 | 96.66 | 86.37 |
| nsclc_ffpe_Lung5_Rep1 | CosMx | lung | Bruker CosMx SMI | 100,292 | 960 | 98.40 | 86.82 |
| nsclc_ffpe_Lung5_Rep2 | CosMx | lung | Bruker CosMx SMI | 106,660 | 960 | 96.53 | 85.44 |
| nsclc_ffpe_Lung5_Rep3 | CosMx | lung | Bruker CosMx SMI | 100,264 | 960 | 99.13 | 88.24 |
| nsclc_ffpe_Lung6 | CosMx | lung | Bruker CosMx SMI | 93,795 | 960 | 98.57 | 86.48 |
| nsclc_ffpe_Lung9_Rep1 | CosMx | lung | Bruker CosMx SMI | 91,972 | 960 | 94.70 | 87.16 |
| nsclc_ffpe_Lung9_Rep2 | CosMx | lung | Bruker CosMx SMI | 150,504 | 960 | 99.36 | 89.79 |
| lymph_node | CosMx | lymph_node | Bruker CosMx SMI | 1,852,946 | 6,174 | 20.29 | 92.55 |

**Supplementary Table 2**: **Composition of the held-out downstream benchmark datasets.** The "Cells $\leq$ 300 (%)" column reports the percentage of cells in which no more than 300 genes have non-zero expression. The "Sparsity (%)" column reports the percentage of zero cell–gene entries. Both statistics are computed from each post-QC, gene-vocabulary-filtered full-panel expression matrix.

| Dataset | Platform | Organ | Source | Cells | Panel | Cells $\leq$ 300 (%) | Sparsity (%) |
|---|---|---|---|---|---|---|---|
| MERFISH human brain | MERFISH | brain | Publication [37] | 218,099 | 300 | 100.00 | 80.62 |
| CosMx liver normal | CosMx | liver | Bruker CosMx SMI | 332,877 | 999 | 99.53 | 86.36 |
| CosMx liver cancer | CosMx | liver | Bruker CosMx SMI | 460,441 | 999 | 74.96 | 78.15 |
| Xenium lung fibrosis | Xenium | lung | Publication [38] | 1,630,319 | 342 | 100.00 | 89.43 |

**Supplementary Table 3**: **Estimated cumulative coverage of detected cell–gene entries during pretraining.** For each HumanST-46M slide, coverage denotes the entry-weighted probability that a detected cell–gene entry has been included in the model input at least once. The “1 exposure” column assumes one exposure per cell, whereas the remaining columns use the expected number of cell exposures accumulated by the indicated optimizer step under the pretraining sampler. Cells are weighted by their numbers of detected genes, and values are rounded to two decimal places.

| Slide | Panel | 1 exposure | 500 steps | 1,000 steps | 2,000 steps | 5,000 steps | 10,000 steps |
|---|---|---|---|---|---|---|---|
| HuBrain | 815 | 95.13% | 95.71% | 99.18% | 99.95% | 100.00% | 100.00% |
| HuBrain_1K_Alzheimer | 960 | 99.21% | 99.32% | 99.90% | 100.00% | 100.00% | 100.00% |
| HubcTMA_R1 | 815 | 99.83% | 99.86% | 99.99% | 100.00% | 100.00% | 100.00% |
| HubcTMA_R2 | 815 | 99.99% | 100.00% | 100.00% | 100.00% | 100.00% | 100.00% |
| HubcTMA_R3 | 815 | 100.00% | 100.00% | 100.00% | 100.00% | 100.00% | 100.00% |
| HubcTMA_R4 | 815 | 100.00% | 100.00% | 100.00% | 100.00% | 100.00% | 100.00% |
| HubcTMA_R5 | 815 | 100.00% | 100.00% | 100.00% | 100.00% | 100.00% | 100.00% |
| HumanBreastCancerPatient1 | 500 | 99.97% | 99.97% | 100.00% | 100.00% | 100.00% | 100.00% |
| HuColonCan | 815 | 99.98% | 99.98% | 100.00% | 100.00% | 100.00% | 100.00% |
| HumanColonCancerPatient1 | 500 | 100.00% | 100.00% | 100.00% | 100.00% | 100.00% | 100.00% |
| HumanColonCancerPatient2 | 500 | 100.00% | 100.00% | 100.00% | 100.00% | 100.00% | 100.00% |
| HumanLiverCancerPatient1 | 500 | 100.00% | 100.00% | 100.00% | 100.00% | 100.00% | 100.00% |
| HumanLiverCancerPatient2 | 500 | 100.00% | 100.00% | 100.00% | 100.00% | 100.00% | 100.00% |
| HumanLungCancerPatient1 | 500 | 100.00% | 100.00% | 100.00% | 100.00% | 100.00% | 100.00% |
| HumanLungCancerPatient2 | 500 | 100.00% | 100.00% | 100.00% | 100.00% | 100.00% | 100.00% |
| HumanOvarianCancerPatient1 | 500 | 100.00% | 100.00% | 100.00% | 100.00% | 100.00% | 100.00% |
| HumanOvarianCancerPatient2Slice1 | 500 | 100.00% | 100.00% | 100.00% | 100.00% | 100.00% | 100.00% |
| HumanOvarianCancerPatient2Slice2 | 500 | 100.00% | 100.00% | 100.00% | 100.00% | 100.00% | 100.00% |
| HumanOvarianCancerPatient2Slice3 | 500 | 100.00% | 100.00% | 100.00% | 100.00% | 100.00% | 100.00% |
| HumanProstateCancerPatient1 | 500 | 100.00% | 100.00% | 100.00% | 100.00% | 100.00% | 100.00% |
| HumanProstateCancerPatient2 | 500 | 100.00% | 100.00% | 100.00% | 100.00% | 100.00% | 100.00% |
| HumanMelanomaPatient1 | 500 | 100.00% | 100.00% | 100.00% | 100.00% | 100.00% | 100.00% |
| HumanMelanomaPatient2 | 500 | 100.00% | 100.00% | 100.00% | 100.00% | 100.00% | 100.00% |
| HumanUterineCancerPatient1 | 500 | 100.00% | 100.00% | 100.00% | 100.00% | 100.00% | 100.00% |
| HumanUterineCancerPatient2-RACostain | 500 | 100.00% | 100.00% | 100.00% | 100.00% | 100.00% | 100.00% |
| HumanUterineCancerPatient2-ROCostain | 500 | 100.00% | 100.00% | 100.00% | 100.00% | 100.00% | 100.00% |
| GSE284005_merfish_all::ms10r0 | 500 | 100.00% | 100.00% | 100.00% | 100.00% | 100.00% | 100.00% |
| GSE284005_merfish_all::ms11r1 | 500 | 100.00% | 100.00% | 100.00% | 100.00% | 100.00% | 100.00% |

*(continued on next page)*

(continued from previous page)

| Slide | Panel | 1 exposure | 500 steps | 1,000 steps | 2,000 steps | 5,000 steps | 10,000 steps |
|---|---|---|---|---|---|---|---|
| GSE284005_merfish_all::ms12r0 | 500 | 100.00% | 100.00% | 100.00% | 100.00% | 100.00% | 100.00% |
| GSE284005_merfish_all::ms12r1 | 500 | 100.00% | 100.00% | 100.00% | 100.00% | 100.00% | 100.00% |
| GSE284005_merfish_all::ms15r0 | 500 | 100.00% | 100.00% | 100.00% | 100.00% | 100.00% | 100.00% |
| GSE284005_merfish_all::ms15r1 | 500 | 100.00% | 100.00% | 100.00% | 100.00% | 100.00% | 100.00% |
| GSE284005_merfish_all::ms16r0 | 500 | 100.00% | 100.00% | 100.00% | 100.00% | 100.00% | 100.00% |
| GSE284005_merfish_all::ms16r1 | 500 | 100.00% | 100.00% | 100.00% | 100.00% | 100.00% | 100.00% |
| GSE284005_merfish_all::ms1r1 | 500 | 100.00% | 100.00% | 100.00% | 100.00% | 100.00% | 100.00% |
| GSE284005_merfish_all::ms1r2 | 500 | 100.00% | 100.00% | 100.00% | 100.00% | 100.00% | 100.00% |
| GSE284005_merfish_all::ms4r1 | 500 | 100.00% | 100.00% | 100.00% | 100.00% | 100.00% | 100.00% |
| GSE284005_merfish_all::ms4r2 | 500 | 100.00% | 100.00% | 100.00% | 100.00% | 100.00% | 100.00% |
| GSE284005_merfish_all::ms5r1 | 500 | 100.00% | 100.00% | 100.00% | 100.00% | 100.00% | 100.00% |
| GSE284005_merfish_all::ms6r0 | 500 | 100.00% | 100.00% | 100.00% | 100.00% | 100.00% | 100.00% |
| GSE284005_merfish_all::ms6r1 | 500 | 100.00% | 100.00% | 100.00% | 100.00% | 100.00% | 100.00% |
| GSE284005_merfish_all::ms7r1 | 500 | 100.00% | 100.00% | 100.00% | 100.00% | 100.00% | 100.00% |
| GSE284005_merfish_all::ms8r0 | 500 | 100.00% | 100.00% | 100.00% | 100.00% | 100.00% | 100.00% |
| gw15::UMB1117—F1a | 300 | 100.00% | 100.00% | 100.00% | 100.00% | 100.00% | 100.00% |
| gw15::UMB1117—F1b | 300 | 100.00% | 100.00% | 100.00% | 100.00% | 100.00% | 100.00% |
| gw15::UMB1117—F2a | 300 | 100.00% | 100.00% | 100.00% | 100.00% | 100.00% | 100.00% |
| gw15::UMB1117—F2b | 300 | 100.00% | 100.00% | 100.00% | 100.00% | 100.00% | 100.00% |
| gw15::UMB1117—O1 | 300 | 100.00% | 100.00% | 100.00% | 100.00% | 100.00% | 100.00% |
| gw15::UMB1117—P1 | 300 | 100.00% | 100.00% | 100.00% | 100.00% | 100.00% | 100.00% |
| gw15::UMB1117—T1 | 300 | 100.00% | 100.00% | 100.00% | 100.00% | 100.00% | 100.00% |
| gw15::UMB1367—O1 | 300 | 100.00% | 100.00% | 100.00% | 100.00% | 100.00% | 100.00% |
| gw15::UMB1367—P1 | 300 | 100.00% | 100.00% | 100.00% | 100.00% | 100.00% | 100.00% |
| gw18_umb1759 | 1,000 | 100.00% | 100.00% | 100.00% | 100.00% | 100.00% | 100.00% |
| gw20::FB080—F1 | 300 | 100.00% | 100.00% | 100.00% | 100.00% | 100.00% | 100.00% |
| gw20::FB080—F2a | 300 | 100.00% | 100.00% | 100.00% | 100.00% | 100.00% | 100.00% |
| gw20::FB080—F2b | 300 | 100.00% | 100.00% | 100.00% | 100.00% | 100.00% | 100.00% |
| gw20::FB080—O1a | 300 | 100.00% | 100.00% | 100.00% | 100.00% | 100.00% | 100.00% |
| gw20::FB080—O1b | 300 | 100.00% | 100.00% | 100.00% | 100.00% | 100.00% | 100.00% |
| gw20::FB080—O1c | 300 | 100.00% | 100.00% | 100.00% | 100.00% | 100.00% | 100.00% |
| gw20::FB080—O1d | 300 | 100.00% | 100.00% | 100.00% | 100.00% | 100.00% | 100.00% |
| gw20::FB080—P1a | 300 | 100.00% | 100.00% | 100.00% | 100.00% | 100.00% | 100.00% |
| gw20::FB080—P1b | 300 | 100.00% | 100.00% | 100.00% | 100.00% | 100.00% | 100.00% |

(continued on next page)

*(continued from previous page)*

| Slide | Panel | 1 exposure | 500 steps | 1,000 steps | 2,000 steps | 5,000 steps | 10,000 steps |
|---|---|---|---|---|---|---|---|
| gw20::FB080—P2 | 300 | 100.00% | 100.00% | 100.00% | 100.00% | 100.00% | 100.00% |
| gw20::FB080—T1 | 300 | 100.00% | 100.00% | 100.00% | 100.00% | 100.00% | 100.00% |
| gw20::FB121—F1 | 300 | 100.00% | 100.00% | 100.00% | 100.00% | 100.00% | 100.00% |
| gw20::FB121—F2 | 300 | 100.00% | 100.00% | 100.00% | 100.00% | 100.00% | 100.00% |
| gw20::FB121—O1 | 300 | 100.00% | 100.00% | 100.00% | 100.00% | 100.00% | 100.00% |
| gw20::FB121—P1 | 300 | 100.00% | 100.00% | 100.00% | 100.00% | 100.00% | 100.00% |
| gw20::FB121—P2 | 300 | 100.00% | 100.00% | 100.00% | 100.00% | 100.00% | 100.00% |
| gw20::FB121—T1 | 300 | 100.00% | 100.00% | 100.00% | 100.00% | 100.00% | 100.00% |
| gw20_umb1031 | 1,000 | 100.00% | 100.00% | 100.00% | 100.00% | 100.00% | 100.00% |
| gw21_umb1932 | 1,000 | 100.00% | 100.00% | 100.00% | 100.00% | 100.00% | 100.00% |
| gw22::FB123—F1 | 300 | 100.00% | 100.00% | 100.00% | 100.00% | 100.00% | 100.00% |
| gw22::FB123—F2 | 300 | 100.00% | 100.00% | 100.00% | 100.00% | 100.00% | 100.00% |
| gw22::FB123—F3 | 300 | 100.00% | 100.00% | 100.00% | 100.00% | 100.00% | 100.00% |
| gw22::FB123—O1 | 300 | 100.00% | 100.00% | 100.00% | 100.00% | 100.00% | 100.00% |
| gw22::FB123—O2 | 300 | 100.00% | 100.00% | 100.00% | 100.00% | 100.00% | 100.00% |
| gw22::FB123—P1 | 300 | 100.00% | 100.00% | 100.00% | 100.00% | 100.00% | 100.00% |
| adata_healthy_diseased_merfish::AM031 | 317 | 100.00% | 100.00% | 100.00% | 100.00% | 100.00% | 100.00% |
| adata_healthy_diseased_merfish::AM042 | 317 | 100.00% | 100.00% | 100.00% | 100.00% | 100.00% | 100.00% |
| adata_healthy_diseased_merfish::AM048 | 317 | 100.00% | 100.00% | 100.00% | 100.00% | 100.00% | 100.00% |
| adata_healthy_diseased_merfish::AM061 | 317 | 100.00% | 100.00% | 100.00% | 100.00% | 100.00% | 100.00% |
| adata_healthy_diseased_merfish::AM062 | 317 | 100.00% | 100.00% | 100.00% | 100.00% | 100.00% | 100.00% |
| adata_healthy_diseased_merfish::AM066 | 317 | 100.00% | 100.00% | 100.00% | 100.00% | 100.00% | 100.00% |
| adata_healthy_diseased_merfish::AM072 | 317 | 100.00% | 100.00% | 100.00% | 100.00% | 100.00% | 100.00% |
| Xenium_V1_FFPE_Human_Brain_Healthy_With_Addon | 319 | 100.00% | 100.00% | 100.00% | 100.00% | 100.00% | 100.00% |
| Xenium_V1_Human_Brain_GBM_FFPE | 480 | 100.00% | 100.00% | 100.00% | 100.00% | 100.00% | 100.00% |
| Xenium_Prime_Breast_Cancer_FFPE | 5,090 | 92.47% | 93.09% | 97.69% | 99.61% | 99.99% | 100.00% |
| Xenium_V1_FFPE_Human_Breast_IDC | 280 | 100.00% | 100.00% | 100.00% | 100.00% | 100.00% | 100.00% |
| Xenium_V1_FFPE_Human_Breast_IDC_Big_1 | 280 | 100.00% | 100.00% | 100.00% | 100.00% | 100.00% | 100.00% |
| Xenium_V1_FFPE_Human_Breast_IDC_Big_2 | 280 | 100.00% | 100.00% | 100.00% | 100.00% | 100.00% | 100.00% |
| Xenium_V1_FFPE_Human_Breast_IDC_With_Addon | 380 | 100.00% | 100.00% | 100.00% | 100.00% | 100.00% | 100.00% |
| Xenium_V1_FFPE_Human_Breast_ILC | 280 | 100.00% | 100.00% | 100.00% | 100.00% | 100.00% | 100.00% |
| Xenium_V1_Human_Colorectal_Cancer_Addon_FFPE | 480 | 100.00% | 100.00% | 100.00% | 100.00% | 100.00% | 100.00% |
| Xenium_V1_hColon_Cancer_Add_on_FFPE | 425 | 100.00% | 100.00% | 100.00% | 100.00% | 100.00% | 100.00% |
| Xenium_V1_hColon_Cancer_Base_FFPE | 325 | 100.00% | 100.00% | 100.00% | 100.00% | 100.00% | 100.00% |

*(continued on next page)*

*(continued from previous page)*

| Slide | Panel | 1 exposure | 500 steps | 1,000 steps | 2,000 steps | 5,000 steps | 10,000 steps |
|---|---|---|---|---|---|---|---|
| Xenium_V1_hColon_Non_diseased_Add_on_FFPE | 425 | 100.00% | 100.00% | 100.00% | 100.00% | 100.00% | 100.00% |
| Xenium_V1_hColon_Non_diseased_Base_FFPE | 325 | 100.00% | 100.00% | 100.00% | 100.00% | 100.00% | 100.00% |
| Xenium_V1_Human_Kidney_FFPE_Protein_updated | 405 | 100.00% | 100.00% | 100.00% | 100.00% | 100.00% | 100.00% |
| Xenium_V1_hKidney_cancer_section | 377 | 100.00% | 100.00% | 100.00% | 100.00% | 100.00% | 100.00% |
| Xenium_V1_hKidney_nondiseased_section | 377 | 100.00% | 100.00% | 100.00% | 100.00% | 100.00% | 100.00% |
| Xenium_V1_hLiver_cancer_section_FFPE | 474 | 100.00% | 100.00% | 100.00% | 100.00% | 100.00% | 100.00% |
| Xenium_V1_hLiver_nondiseased_section_FFPE | 377 | 100.00% | 100.00% | 100.00% | 100.00% | 100.00% | 100.00% |
| Xenium_Preview_Human_Lung_Cancer_With_Add_on_2_FFPE | 392 | 100.00% | 100.00% | 100.00% | 100.00% | 100.00% | 100.00% |
| Xenium_Preview_Human_Non_diseased_Lung_With_Add_on_FFPE | 392 | 100.00% | 100.00% | 100.00% | 100.00% | 100.00% | 100.00% |
| Xenium_V1_Human_Lung_Cancer_Addon_FFPE | 480 | 100.00% | 100.00% | 100.00% | 100.00% | 100.00% | 100.00% |
| Xenium_V1_hLung_cancer_section | 377 | 100.00% | 100.00% | 100.00% | 100.00% | 100.00% | 100.00% |
| Xenium_Prime_Human_Lymph_Node_Reactive_FFPE | 4,624 | 92.63% | 93.33% | 98.04% | 99.70% | 99.99% | 100.00% |
| Xenium_Prime_Human_Ovary_Cancer_FF | 5,001 | 36.19% | 38.07% | 58.42% | 79.75% | 96.86% | 99.78% |
| Xenium_Prime_Human_Ovary_FF | 5,001 | 34.37% | 36.18% | 56.12% | 77.73% | 96.22% | 99.71% |
| Xenium_Prime_Ovarian_Cancer_FFPE_XRrun | 5,090 | 82.78% | 84.03% | 93.83% | 98.69% | 99.96% | 100.00% |
| Xenium_V1_Human_Ovary_Cancer_FF | 477 | 100.00% | 100.00% | 100.00% | 100.00% | 100.00% | 100.00% |
| frontal_cortex | 6,077 | 46.60% | 48.32% | 65.53% | 81.78% | 95.68% | 99.36% |
| colon_wtx | 18,935 | 26.07% | 27.49% | 43.55% | 63.11% | 86.51% | 96.29% |
| nsclc_ffpe_Lung12 | 960 | 97.57% | 97.83% | 99.47% | 99.95% | 100.00% | 100.00% |
| nsclc_ffpe_Lung13 | 960 | 98.93% | 99.07% | 99.84% | 99.99% | 100.00% | 100.00% |
| nsclc_ffpe_Lung5_Rep1 | 960 | 99.60% | 99.66% | 99.95% | 100.00% | 100.00% | 100.00% |
| nsclc_ffpe_Lung5_Rep2 | 960 | 99.02% | 99.15% | 99.86% | 99.99% | 100.00% | 100.00% |
| nsclc_ffpe_Lung5_Rep3 | 960 | 99.74% | 99.78% | 99.97% | 100.00% | 100.00% | 100.00% |
| nsclc_ffpe_Lung6 | 960 | 99.63% | 99.68% | 99.95% | 100.00% | 100.00% | 100.00% |
| nsclc_ffpe_Lung9_Rep1 | 960 | 98.08% | 98.33% | 99.70% | 99.98% | 100.00% | 100.00% |
| nsclc_ffpe_Lung9_Rep2 | 960 | 99.80% | 99.83% | 99.98% | 100.00% | 100.00% | 100.00% |
| lymph_node | 6,174 | 61.96% | 64.09% | 82.84% | 94.99% | 99.70% | 99.99% |

**Supplementary Table 4: Pretraining configuration of NexuST.**

| Category | Hyperparameter | Value |
|---|---|---|
| Model | Gene vocabulary size | 19,228 |
| | Number of organs | 11 |
| | Maximum gene tokens per cell | $M = 300$ |
| | Embedding dimension | $D = 512$ |
| | Hierarchical encoder blocks | 8 |
| | Transformer layers | 16 |
| | Attention heads | 8 |
| | Dense MoE experts | 8 |
| | Dropout | 0.1 |
| Input | Cells per patch | $N = 512$ |
| | Gene value mask ratio | 20% |
| | Maximum expression value | 512 |
| Optimization | Optimizer | AdamW |
| | Learning rate | $3 \times 10^{-4}$ |
| | Weight decay | 0.01 |
| | Adam betas | $(0.9, 0.999)$ |
| | Training steps | 10,000 |
| | Warm-up steps | 1,000 |
| | Scheduler | Cosine decay |
| | Minimum learning rate | $10^{-6}$ |
| Training | Gradient clipping | 1.0 |
| | Per-GPU batch size | 6 |
| | Global batch size | 192 |
| | Precision | bfloat16 mixed precision |
| | Parallelism strategy | FSDP full sharding |
| | Activation checkpointing | Enabled |
| Hardware | GPUs | 32 NVIDIA A100 64GB GPUs |
| | Nodes | 8 nodes, 4 GPUs per node |
| | Pretraining time | Approximately 40 hours |

**Supplementary Table 5**: **Dataset-specific radii used for niche composition prediction.** For each dataset, the four radii were selected to define neighbourhoods containing approximately 10, 20, 50, and 100 neighbouring cells, respectively.

| Dataset | ∼10 neighbours | ∼20 neighbours | ∼50 neighbours | ∼100 neighbours |
|---|---|---|---|---|
| MERFISH human brain | 0.29 | 0.42 | 0.68 | 0.99 |
| CosMx liver normal | 0.25 | 0.35 | 0.56 | 0.81 |
| CosMx liver cancer | 0.21 | 0.30 | 0.49 | 0.71 |
| Xenium lung fibrosis | 0.12 | 0.17 | 0.28 | 0.40 |

**Supplementary Table 6**: **Per-dataset radii for neighbourhood density prediction.** For each dataset, neighbourhood density was evaluated at three spatial radii (in normalised coordinate units), selected via `--radius_idx` $\in \{0, 1, 2\}$. Each radius defines a separate prediction task that is trained and reported independently.

| Dataset | Radius 0 | Radius 1 | Radius 2 |
|---|---|---|---|
| MERFISH human brain | 0.25 | 0.50 | 1.00 |
| CosMx liver normal | 0.25 | 0.50 | 0.75 |
| CosMx liver cancer | 0.25 | 0.50 | 0.75 |
| Xenium lung fibrosis | 0.15 | 0.25 | 0.40 |

**Supplementary Table 7**: **HVG300 coverage across model vocabularies and shared genes used for held-out gene recovery.** For each downstream dataset, the four model columns report the number of genes retained after intersecting the shared HVG300 candidate panel with each model's pretrained vocabulary. The Shared column denotes the intersection retained by all four foundation models. The Held-out column gives the number of genes randomly selected from this shared set as prediction targets for held-out gene recovery; the remaining shared genes form the observed input panel.

| Dataset | NexuST | scGPT-spatial | Nicheformer | CellPLM | Shared | Held-out |
|---|---|---|---|---|---|---|
| MERFISH human brain | 300 | 295 | 269 | 266 | 266 | 53 |
| CosMx liver normal | 300 | 299 | 299 | 287 | 286 | 57 |
| CosMx liver cancer | 300 | 299 | 298 | 287 | 286 | 57 |
| Xenium lung fibrosis | 300 | 300 | 299 | 298 | 297 | 59 |

**Supplementary Table 8**: **Linear-probing configurations.**

| Task | Loss | Learning rate | Batch size | Max epochs | Patience |
|---|---|---|---|---|---|
| Cell annotation | CE | $1 \times 10^{-1}$ | 64 | 50 | 5 |
| Region prediction | CE | $1 \times 10^{-2}$ | 64 | 50 | 5 |
| Held-out gene recovery | MSE | $1 \times 10^{-3}$ | 64 | 50 | 5 |
| Niche composition prediction | MSE | $1 \times 10^{-3}$ | 64 | 50 | 5 |
| Density prediction | MSE | $1 \times 10^{-3}$ | 64 | 50 | 5 |

**Supplementary Table 9**: **Fine-tuning learning rates across foundation models.**

| Method | Cell-type annotation | Region prediction | Held-out gene recovery | Niche composition prediction |
|---|---|---|---|---|
| NexuST | $1 \times 10^{-4}$ | $1 \times 10^{-4}$ | $1 \times 10^{-4}$ | $3 \times 10^{-5}$ |
| Nicheformer | $1 \times 10^{-4}$ | $1 \times 10^{-4}$ | $1 \times 10^{-4}$ | $1 \times 10^{-4}$ |
| scGPT-spatial | $1 \times 10^{-4}$ | $1 \times 10^{-4}$ | $1 \times 10^{-4}$ | $1 \times 10^{-4}$ |
| CellPLM | $5 \times 10^{-3}$ | $5 \times 10^{-3}$ | $5 \times 10^{-4}$ | $1 \times 10^{-3}$ |

**Supplementary Table 10**: **Fixed validation FOVs used for attention analysis.** All cells from the five listed FOVs in each dataset were analysed using the pretrained encoder. MERFISH identifiers denote pseudo-FOVs from the 10×10 spatial grid.

| **Dataset** | **FOV name** |
|---|---|
| MERFISH human brain | 10x10_37 |
| | 10x10_25 |
| | 10x10_58 |
| | 10x10_22 |
| | 10x10_51 |
| CosMx liver normal | 174 |
| | 78 |
| | 15 |
| | 69 |
| | 163 |
| CosMx liver cancer | 92 |
| | 137 |
| | 377 |
| | 113 |
| | 216 |
| Xenium lung fibrosis | VUILD105MA1 |
| | VUILD115MA |
| | VUILD96MA |
| | TILD175MA |
| | VUILD96LA |

**Supplementary Table 11**: **Region-label mapping for the Xenium lung fibrosis dataset.** The original C1–C12 identifiers denote the 12 cell-based niches reported by Vannan et al. We assigned descriptive labels reflecting the dominant cell composition and tissue context of each niche and used these labels throughout the region-prediction analyses and figures.

| **Original label** | **Descriptive label used in this work** |
|---|---|
| C1 | Airway |
| C2 | Transitional epithelial |
| C3 | Epithelial detachment |
| C4 | Interstitial fibrotic |
| C5 | AT2-enriched alveolar |
| C6 | T/plasma immune |
| C7 | Mixed immune |
| C8 | Healthy alveolar |
| C9 | Lymphoid/TLS |
| C10 | Adventitial/vascular |
| C11 | Perivascular |
| C12 | Macrophage accumulation |

**Supplementary Table 12: Cell type labels and six analysis lineages in the Xenium lung fibrosis dataset.** Cell counts refer to the held-out validation split used for downstream evaluation. Cell types represented by fewer than 100 validation cells were excluded from the per-cell-type performance heatmap but retained in the complete label set.

| Cell type | Analysis lineage | Validation cells |
|---|---|---|
| Arteriole | Endothelial | 1,847 |
| Capillary | Endothelial | 25,374 |
| Lymphatic | Endothelial | 2,540 |
| Venous | Endothelial | 17,340 |
| AT1 | Epithelial | 5,887 |
| AT2 | Epithelial | 25,162 |
| Basal | Epithelial | 6,173 |
| Goblet | Epithelial | 4,776 |
| KRT5-/KRT17+ | Epithelial | 604 |
| Multiciliated | Epithelial | 18,820 |
| PNEC | Epithelial | 115 |
| Proliferating AT2 | Epithelial | 140 |
| Proliferating Airway | Epithelial | 67 |
| RASC | Epithelial | 1,374 |
| Secretory | Epithelial | 1,960 |
| Transitional AT2 | Epithelial | 2,109 |
| Alveolar Macrophages | Myeloid | 6,054 |
| B cells | Lymphoid | 6,978 |
| Basophils | Myeloid | 132 |
| CD4+ T-cells | Lymphoid | 24,884 |
| CD8+ T-cells | Lymphoid | 17,172 |
| Interstitial Macrophages | Myeloid | 30,951 |
| Langerhans cells | Myeloid | 25 |
| Macrophages - IFN-activated | Myeloid | 271 |
| Mast | Myeloid | 8,238 |
| Migratory DCs | Myeloid | 2,258 |
| Monocytes/MDMs | Myeloid | 8,461 |
| Neutrophils | Myeloid | 9,754 |
| NK/NKT | Lymphoid | 4,708 |
| Plasma | Lymphoid | 8,181 |
| Proliferating B cells | Lymphoid | 2 |
| Proliferating Myeloid | Myeloid | 1,751 |
| Proliferating NK/NKT | Lymphoid | 5 |
| Proliferating T-cells | Lymphoid | 345 |
| SPP1+ Macrophages | Myeloid | 5,671 |
| Tregs | Lymphoid | 2,885 |
| cDCs | Myeloid | 1,747 |
| pDCs | Myeloid | 420 |
| Activated Fibrotic FBs | Fibroblast/stromal | 10,090 |
| Adventitial FBs | Fibroblast/stromal | 3,368 |
| Alveolar FBs | Fibroblast/stromal | 48,637 |
| Inflammatory FBs | Fibroblast/stromal | 4,556 |
| Mesothelial | Fibroblast/stromal | 0 |
| Myofibroblasts | Fibroblast/stromal | 2,696 |
| Proliferating FBs | Fibroblast/stromal | 1,974 |
| SMCs/Pericytes | Mural | 20,757 |
| Subpleural FBs | Fibroblast/stromal | 1,286 |

**Supplementary Table 13**: **Ablation results across downstream datasets.** Cell-type annotation and region prediction use macro-F1 (higher is better), whereas niche-composition prediction uses mean absolute error (MAE; lower is better) at $r_3$. Values are means over three seeds. Ranks were computed from the unrounded seed-averaged metric within each task–dataset combination, and the mean rank in each panel averages across the applicable datasets; lower mean rank is better. All models used the same 25% complete-subslide subset; single-stage variants were pretrained for 2,500 steps and the two-stage model for 2,500 steps per stage. "w/o" denotes "without". The w/o KNN-based patching control randomises patch membership while preserving cells, patch number and size, and per-cell exposure. Bold indicates the best dataset mean and the lowest mean rank.

**a, Cell-type annotation (macro-F1 ↑)**

| Variant | MERFISH human brain | CosMx liver normal | CosMx liver cancer | Xenium lung fibrosis | Mean rank ↓ |
|---|---|---|---|---|---|
| Baseline | 0.8833 | 0.6224 | 0.5141 | 0.5900 | 2.50 |
| w/o cell-level attention | **0.8962** | **0.6295** | 0.5061 | 0.5779 | 2.25 |
| w/o interleaving (stacked) | 0.8914 | 0.6122 | 0.5153 | **0.5928** | **2.00** |
| Two-stage training | 0.8513 | 0.6078 | 0.4950 | 0.5361 | 5.25 |
| w/o MoE | 0.8615 | 0.5731 | **0.5249** | 0.5625 | 4.25 |
| w/o library-size head | 0.8300 | 0.4947 | 0.4355 | 0.4405 | 7.00 |
| w/o KNN-based patching | 0.8677 | 0.5936 | 0.4925 | 0.5646 | 4.75 |

**b, Region prediction (macro-F1 ↑)**

| Variant | CosMx liver normal | CosMx liver cancer | Xenium lung fibrosis | Mean rank ↓ |
|---|---|---|---|---|
| Baseline | 0.5081 | **0.6358** | **0.4252** | **1.33** |
| w/o cell-level attention | 0.4416 | 0.5837 | 0.3794 | 7.00 |
| w/o interleaving (stacked) | 0.5018 | 0.6269 | 0.4238 | 2.33 |
| Two-stage training | **0.5167** | 0.5987 | 0.4174 | 3.33 |
| w/o MoE | 0.4940 | 0.6218 | 0.4135 | 3.67 |
| w/o library-size head | 0.4495 | 0.6108 | 0.3917 | 5.00 |
| w/o KNN-based patching | 0.4872 | 0.6046 | 0.3915 | 5.33 |

**c, Niche-composition prediction at $r_3$ (MAE ↓)**

| Variant | MERFISH human brain | CosMx liver normal | CosMx liver cancer | Xenium lung fibrosis | Mean rank ↓ |
|---|---|---|---|---|---|
| Baseline | **0.04853** | 0.01930 | **0.01454** | 0.01450 | **1.75** |
| w/o cell-level attention | 0.06565 | 0.02065 | 0.01680 | 0.01741 | 7.00 |
| w/o interleaving (stacked) | 0.04891 | 0.01950 | 0.01498 | 0.01424 | 2.75 |
| Two-stage training | 0.05094 | **0.01923** | 0.01460 | **0.01416** | **1.75** |
| w/o MoE | 0.05184 | 0.01960 | 0.01485 | 0.01478 | 3.75 |
| w/o library-size head | 0.05435 | 0.02020 | 0.01511 | 0.01525 | 5.25 |
| w/o KNN-based patching | 0.05791 | 0.01990 | 0.01579 | 0.01555 | 5.75 |

**Supplementary Table 14**: **Rank summary of NexuST ablation variants.** Task-specific mean ranks average the within-dataset ranks reported in Supplementary Table 13; the overall mean rank is the unweighted average across all 11 task–dataset combinations. Ranks were computed from the unrounded seed-averaged metric using the task-appropriate direction. Lower values indicate better and more consistent performance.

| Variant | Annotation | Region | Niche | Overall |
|---|---|---|---|---|
| Baseline | 2.50 | **1.33** | **1.75** | **1.91** |
| w/o interleaving (stacked) | **2.00** | 2.33 | 2.75 | 2.36 |
| Two-stage training | 5.25 | 3.33 | **1.75** | 3.45 |
| w/o MoE | 4.25 | 3.67 | 3.75 | 3.91 |
| w/o cell-level attention | 2.25 | 7.00 | 7.00 | 5.27 |
| w/o KNN-based patching | 4.75 | 5.33 | 5.75 | 5.27 |
| w/o library-size head | 7.00 | 5.00 | 5.25 | 5.82 |

**Supplementary Table 15**: **Full-panel linear-probing results across downstream datasets.** Cell-type annotation and region prediction use macro-F1 (higher is better), whereas niche-composition prediction uses mean absolute error (MAE; lower is better) at $r_3$. Values are means over three seeds. Each foundation model received all measured genes compatible with its pretrained vocabulary using its native input-encoding procedure; the expression-only PCA baseline was fitted using 50 principal components (PCA-50) from the full measured panel. The measured panel sizes were 300 genes for MERFISH human brain, 999 for CosMx liver normal and liver cancer, and 342 for Xenium lung fibrosis. Ranks were computed from the unrounded seed-averaged metric within each task–dataset combination, and the mean rank in each panel averages across the applicable datasets; lower mean rank is better. All methods used the same dataset splits, linear-probing protocol and validation-based early stopping. Bold indicates the best dataset mean and the lowest mean rank.

**a, Cell-type annotation (macro-F1 ↑)**

| Model | MERFISH human brain (300 genes) | CosMx liver normal (999 genes) | CosMx liver cancer (999 genes) | Xenium lung fibrosis (342 genes) | Mean rank ↓ |
|---|---|---|---|---|---|
| NexuST | **0.9176** | **0.6381** | **0.5569** | 0.6480 | **1.25** |
| Nicheformer | 0.9024 | 0.5956 | 0.5143 | **0.6733** | 2.00 |
| scGPT-spatial | 0.1363 | 0.0460 | 0.0616 | 0.0088 | 5.00 |
| CellPLM | 0.6005 | 0.3033 | 0.2443 | 0.3648 | 4.00 |
| PCA-50 | 0.9144 | 0.5333 | 0.3540 | 0.3961 | 2.75 |

**b, Region prediction (macro-F1 ↑)**

| Model | CosMx liver normal (999 genes) | CosMx liver cancer (999 genes) | Xenium lung fibrosis (342 genes) | Mean rank ↓ |
|---|---|---|---|---|
| NexuST | **0.5586** | **0.6766** | **0.4601** | **1.00** |
| Nicheformer | 0.4419 | 0.6019 | 0.4045 | 2.33 |
| scGPT-spatial | 0.1535 | 0.2505 | 0.0397 | 5.00 |
| CellPLM | 0.2249 | 0.4959 | 0.3136 | 4.00 |
| PCA-50 | 0.4653 | 0.6004 | 0.3804 | 2.67 |

**c, Niche-composition prediction at $r_3$ (MAE ↓)**

| Model | MERFISH human brain (300 genes) | CosMx liver normal (999 genes) | CosMx liver cancer (999 genes) | Xenium lung fibrosis (342 genes) | Mean rank ↓ |
|---|---|---|---|---|---|
| NexuST | **0.04143** | **0.01820** | **0.01313** | **0.01316** | **1.00** |
| Nicheformer | 0.06510 | 0.02090 | 0.01607 | 0.01661 | 2.25 |
| scGPT-spatial | 0.10609 | 0.02805 | 0.03227 | 0.02337 | 5.00 |
| CellPLM | 0.08689 | 0.02417 | 0.01990 | 0.01869 | 4.00 |
| PCA-50 | 0.06653 | 0.02084 | 0.01633 | 0.01717 | 2.75 |

### Supplementary Figures

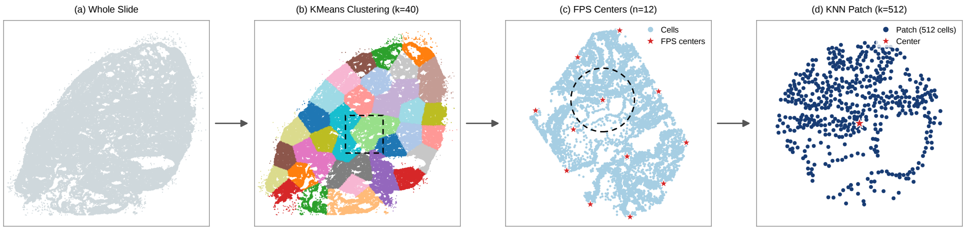


**Fig. 1**: **Spatially coherent patch-sampling pipeline for pretraining.** Schematic of the three-stage procedure used to convert a whole tissue slide into fixed-size cell patches for self-supervised pretraining, shown as a left-to-right magnification cascade on a representative training slide (Xenium Prime Human Ovary Cancer FF; 200,900 cells). Spatial coordinates are first translated and rescaled to a $[0, 100]$ box. **a,** Whole-slide view with all cells shown in grey. **b,** MiniBatch $k$-means clustering on cell coordinates partitions the slide into spatially coherent sub-slides, with the number of clusters chosen adaptively to obtain approximately 5,000 cells per sub-slide (here, $K = 40$; clusters are shown in distinct colours). The dashed box marks one selected sub-slide, magnified in **c**. **c,** Within the selected sub-slide, farthest-point sampling (FPS) selects $n = 12$ well-separated patch centres (red stars). The dashed circle marks one selected centre, magnified in **d**. **d,** For each FPS centre, its $k = 512$ nearest neighbours in spatial coordinates are selected to form one cell patch. This 512-cell patch is the unit fed to NexuST during pretraining. Each sub-slide therefore yields 12 overlapping patches, and sampling is repeated across epochs with independent FPS runs to increase patch diversity.

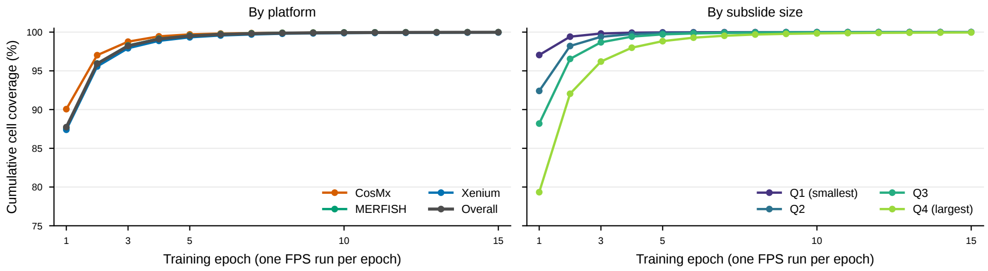


**Fig. 2**: **Cumulative cell coverage under repeated FPS-based patch sampling.** At each training epoch, 12 farthest-point sampling (FPS) centres were independently selected within every $k$-means subslide, with each centre defining an overlapping 512-cell kNN patch. Cumulative coverage denotes the fraction of unique cells included in at least one sampled patch up to and including the indicated epoch. **Left,** cumulative coverage across training epochs, reported overall and separately for each spatial transcriptomics platform. **Right,** cumulative coverage stratified by quartiles of subslide cell count: Q1, the smallest 25% of subslides ($\leq$ 4,279 cells); Q2, 4,280–5,054 cells; Q3, 5,055–5,816 cells; and Q4, the largest 25% of subslides ($>$ 5,816 cells). Repeated FPS resampling rapidly increased coverage across all platforms and subslide-size groups, with even the largest subslides approaching complete cumulative coverage after a small number of epochs despite overlap between patches.

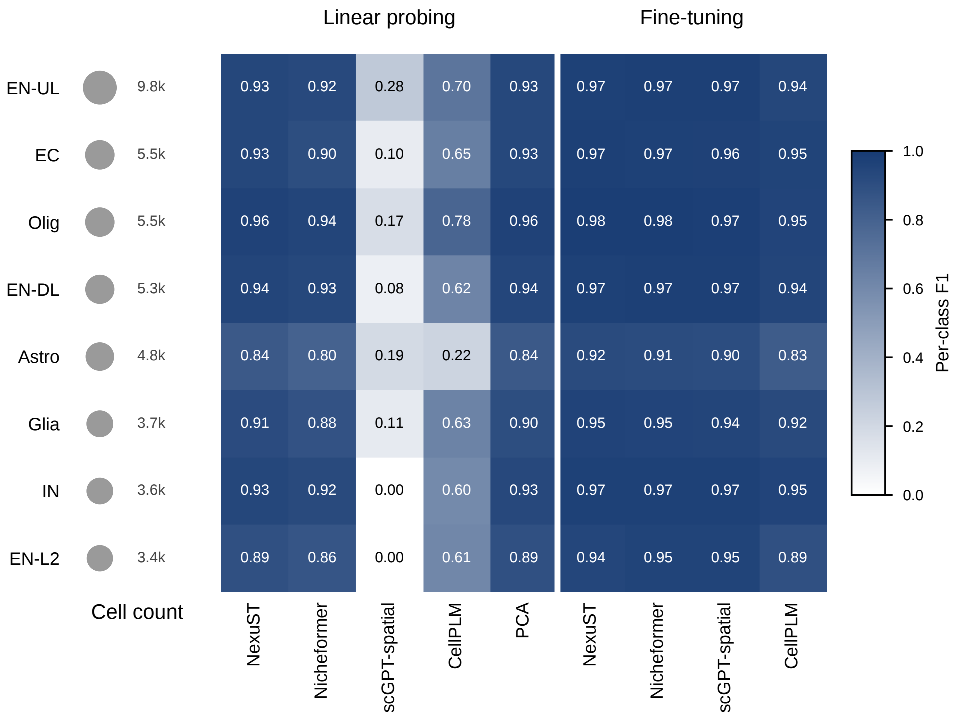


**Fig. 3**: **Per-cell-type cell annotation performance in the MERFISH human brain dataset.** Per-cell-type F1 scores are shown for NexuST, Nicheformer, scGPT-spatial, CellPLM and PCA under linear probing, and for NexuST, Nicheformer, scGPT-spatial and CellPLM under fine-tuning. Rows are cell types ordered by abundance, with the dot size and adjacent label indicating per-type cell count.

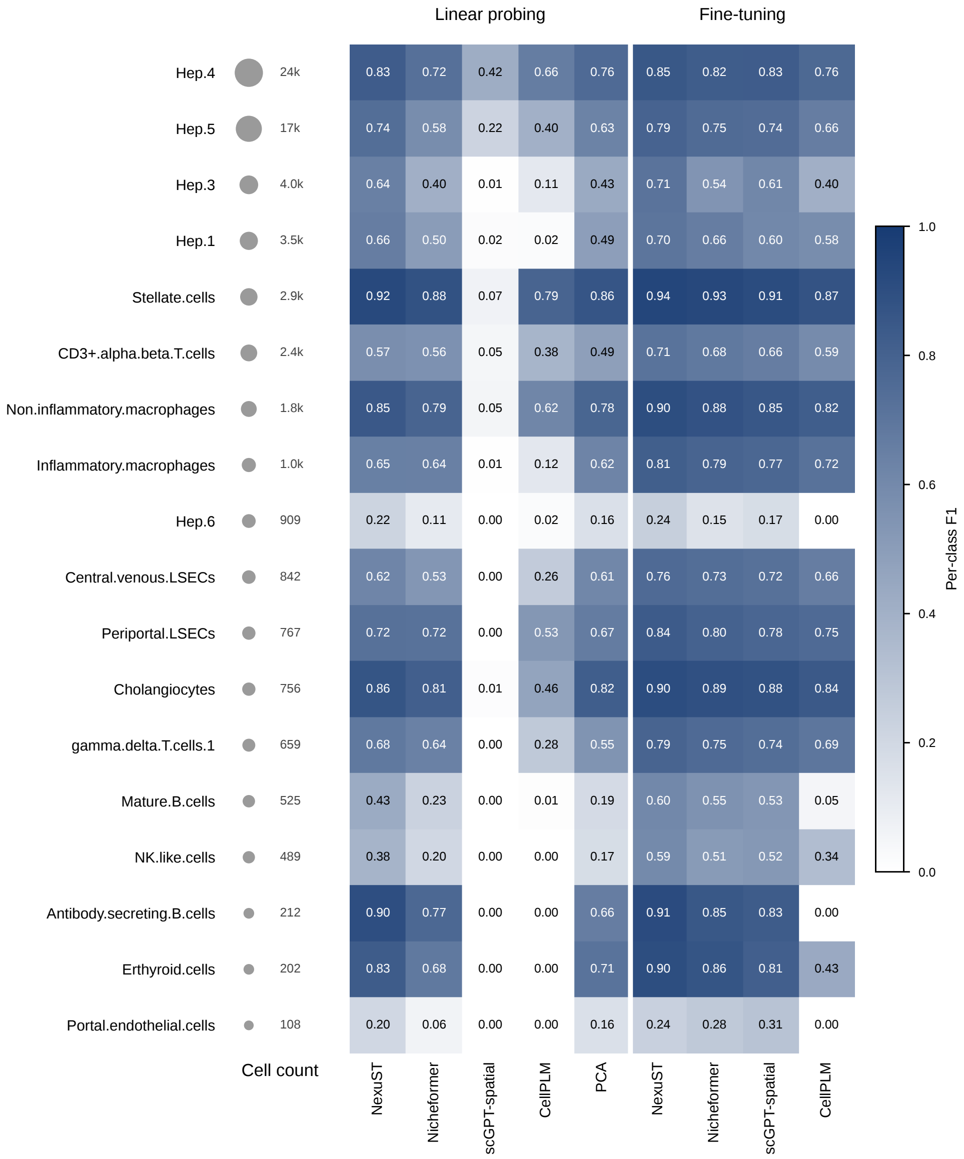


**Fig. 4**: **Per-cell-type cell annotation performance in the CosMx liver normal dataset.** Per-cell-type F1 scores are shown for NexuST, Nicheformer, scGPT-spatial, CellPLM and PCA under linear probing, and for NexuST, Nicheformer, scGPT-spatial and CellPLM under fine-tuning. Rows are cell types ordered by abundance, with the dot size and adjacent label indicating per-type cell count.

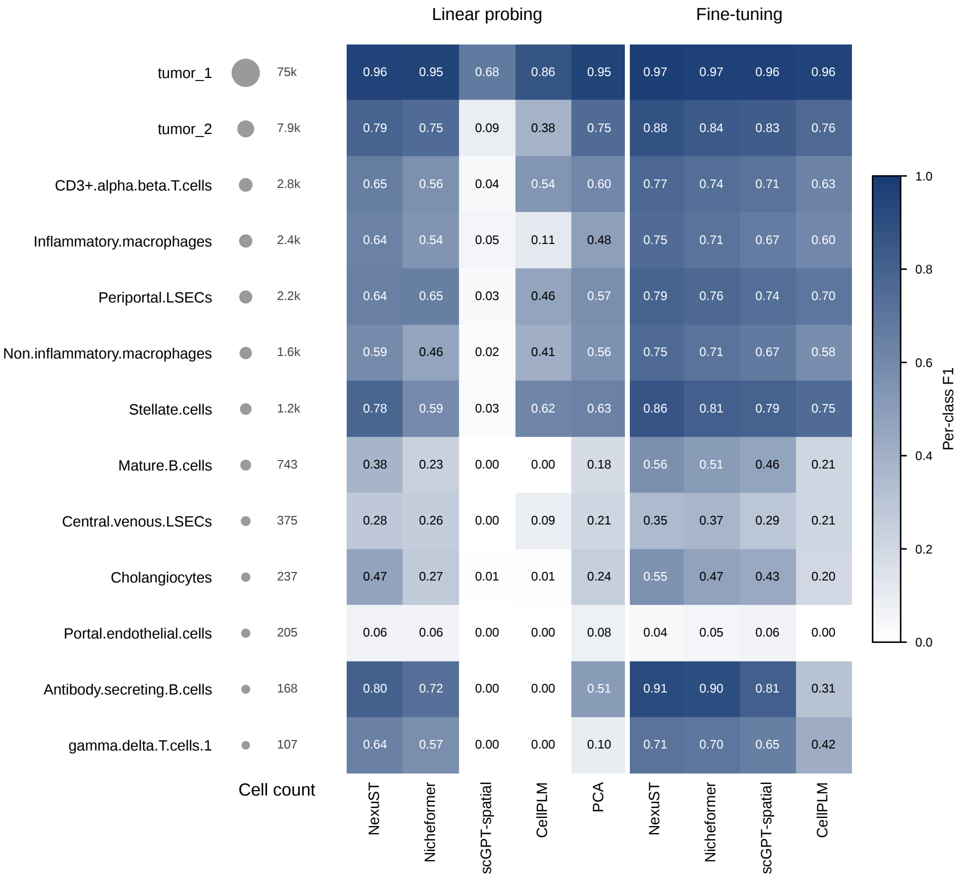


**Fig. 5**: **Per-cell-type cell annotation performance in the CosMx liver cancer dataset.** Per-cell-type F1 scores are shown for NexuST, Nicheformer, scGPT-spatial, CellPLM and PCA under linear probing, and for NexuST, Nicheformer, scGPT-spatial and CellPLM under fine-tuning. Rows are cell types ordered by abundance, with the dot size and adjacent label indicating per-type cell count.

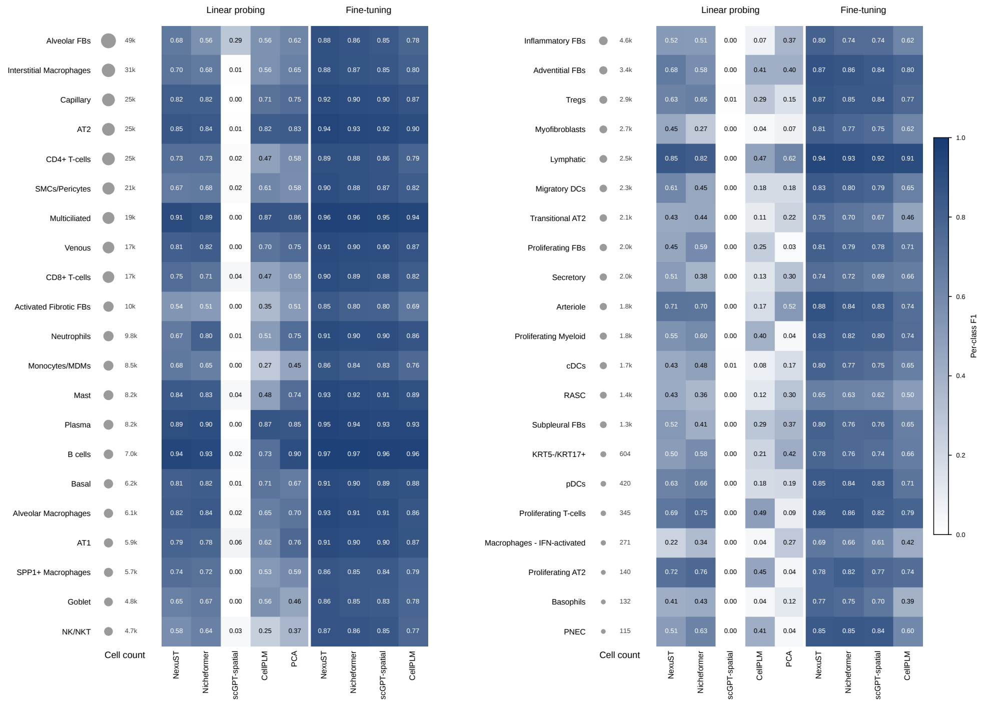


**Fig. 6**: **Per-cell-type cell annotation performance in the Xenium lung fibrosis dataset.** Per-cell-type F1 scores are shown for NexuST, Nicheformer, scGPT-spatial, CellPLM and PCA under linear probing, and for NexuST, Nicheformer, scGPT-spatial and CellPLM under fine-tuning. Rows are cell types ordered by abundance, with the dot size and adjacent label indicating per-type cell count.

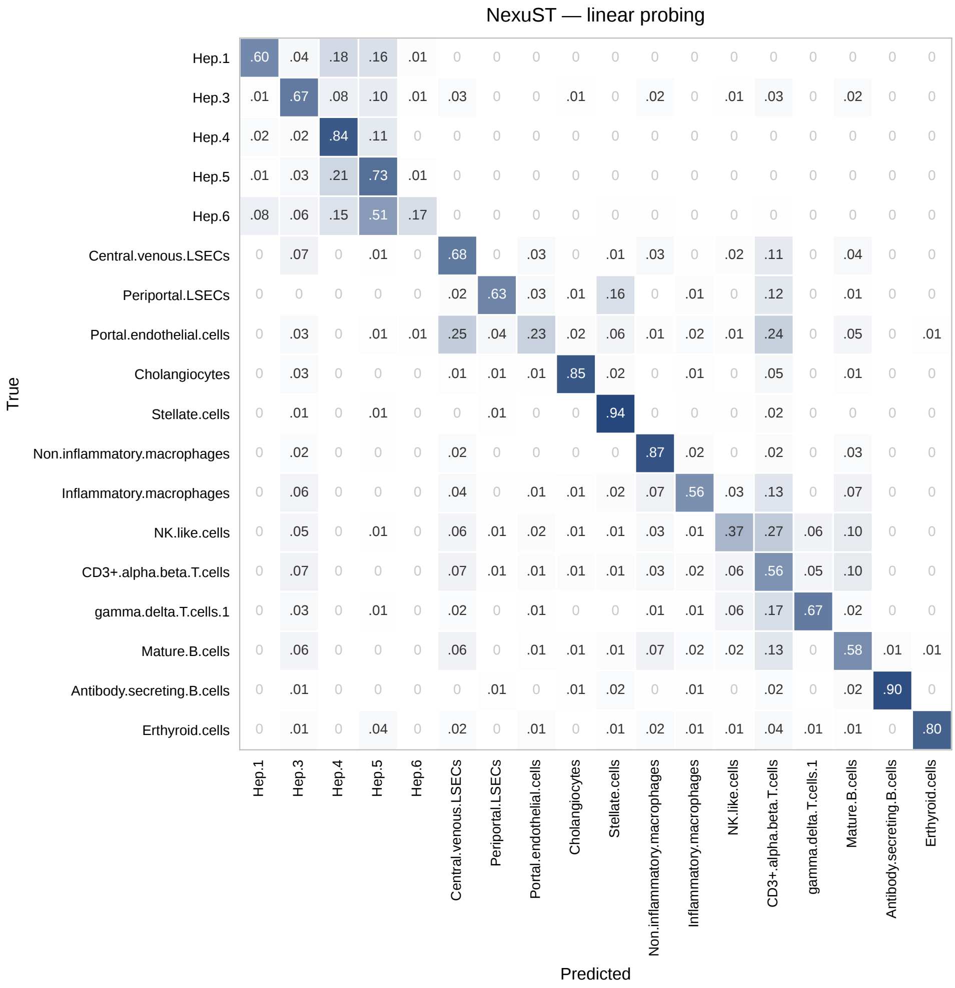


**Fig. 7**: **Cell annotation confusion matrix for NexuST under linear probing in the CosMx liver normal dataset.** Row-normalized confusion matrix over the annotated cell types (each row sums to one); diagonal entries give the per-type recall.

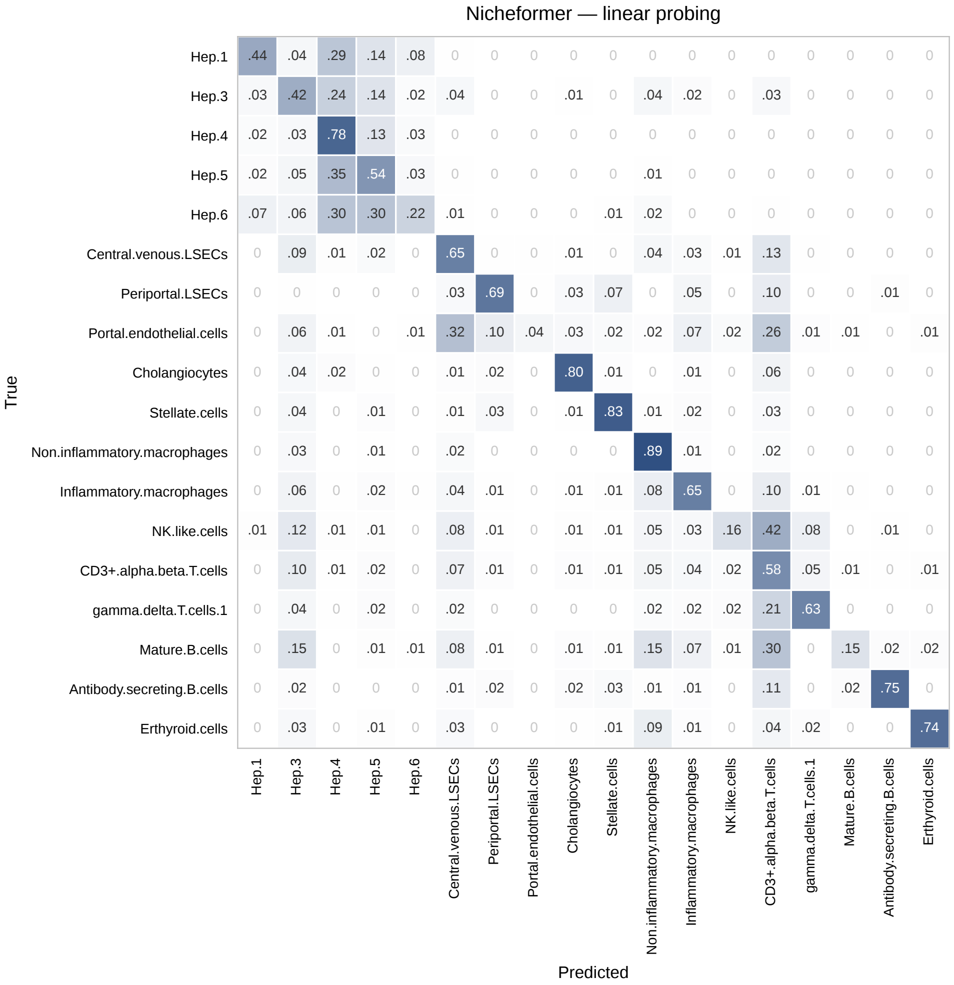


**Fig. 8**: **Cell annotation confusion matrix for Nicheformer under linear probing in the CosMx liver normal dataset.** Row-normalized confusion matrix over the annotated cell types (each row sums to one); diagonal entries give the per-type recall.

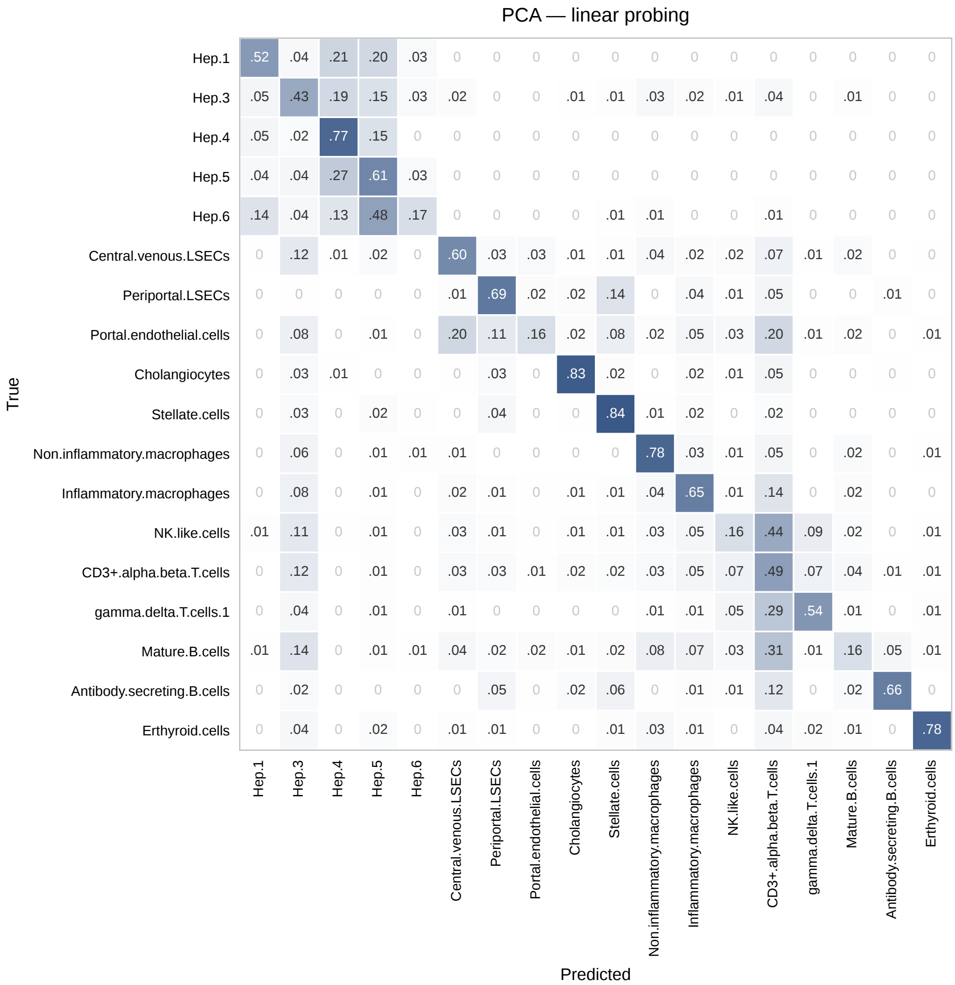


**Fig. 9**: **Cell annotation confusion matrix for PCA under linear probing in the CosMx liver normal dataset.** Row-normalized confusion matrix over the annotated cell types (each row sums to one); diagonal entries give the per-type recall.

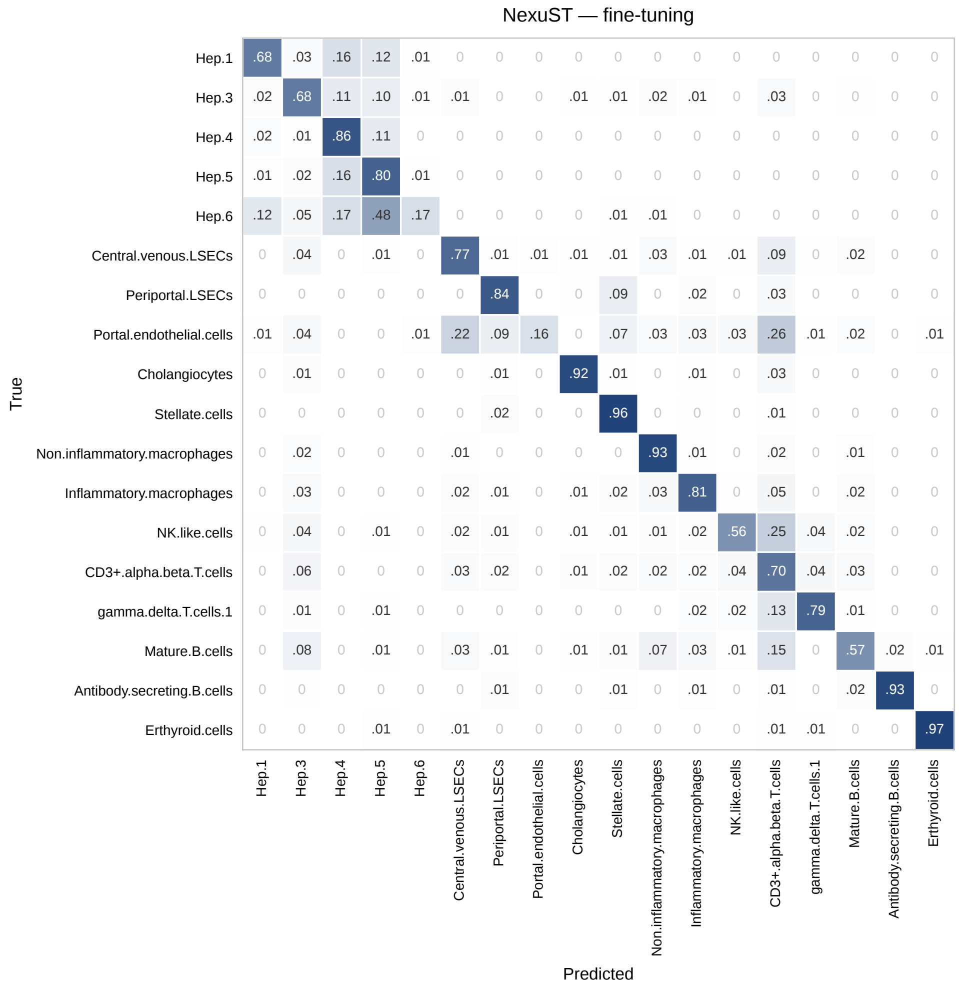


**Fig. 10**: **Cell annotation confusion matrix for NexuST under fine-tuning in the CosMx liver normal dataset.** Row-normalized confusion matrix over the annotated cell types (each row sums to one); diagonal entries give the per-type recall.

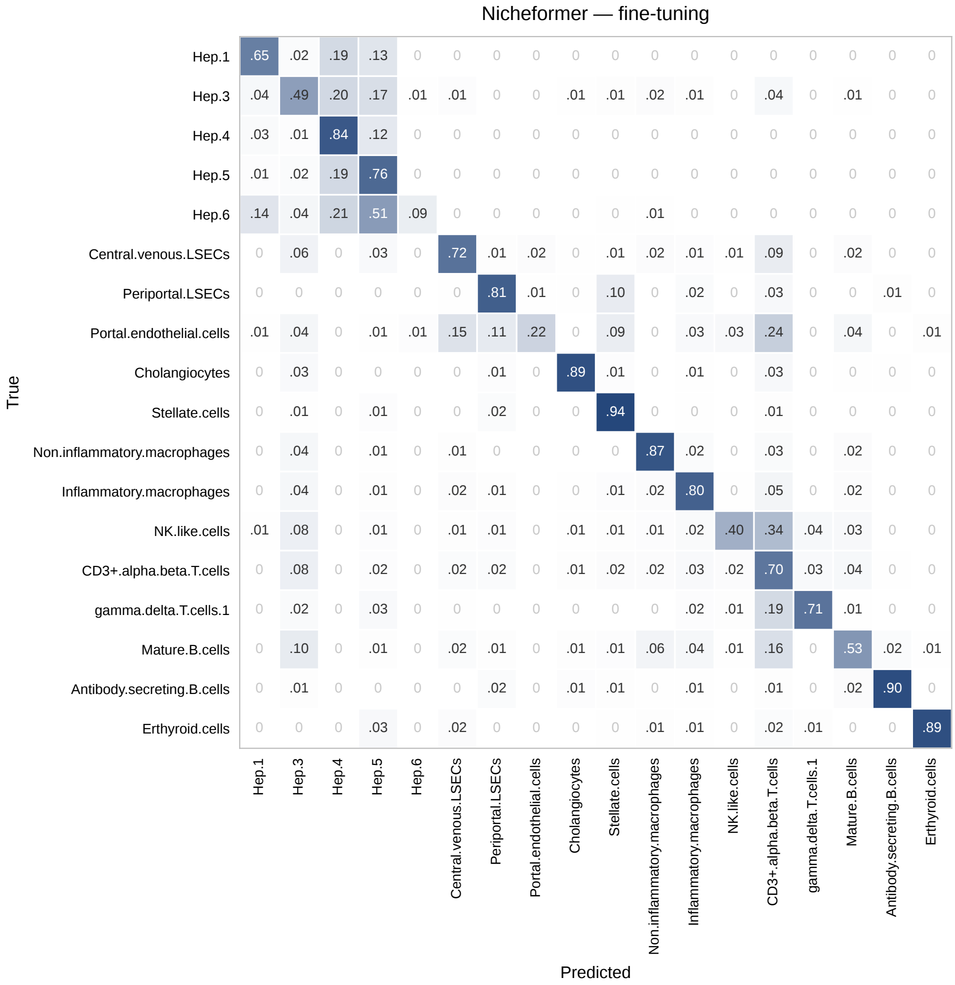


**Fig. 11**: **Cell annotation confusion matrix for Nicheformer under fine-tuning in the CosMx liver normal dataset.** Row-normalized confusion matrix over the annotated cell types (each row sums to one); diagonal entries give the per-type recall.

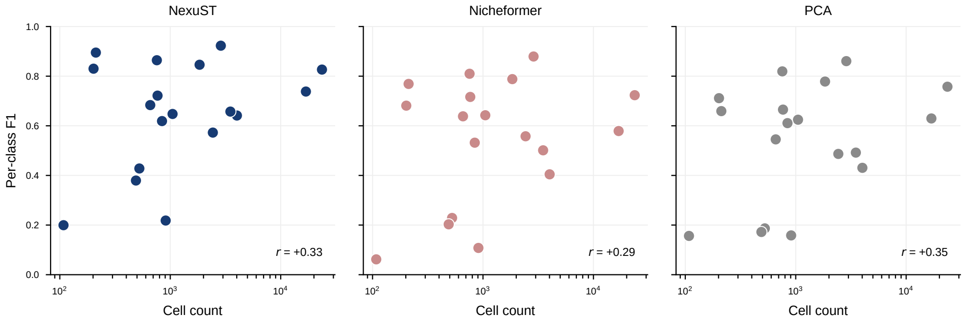


**Fig. 12**: **Per-class F1 versus cell-type abundance in the CosMx liver normal dataset.** Per-class F1 under linear probing against cell count (log scale) for NexuST, Nicheformer, and PCA; the Pearson correlation $r$ is annotated in each panel.

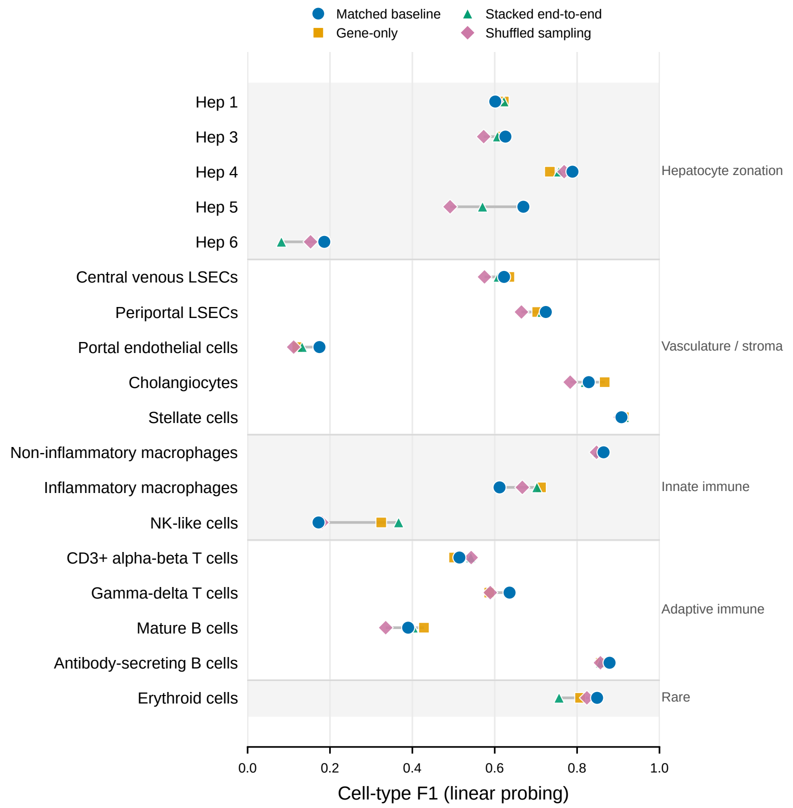


**Fig. 13**: **Cell-type-resolved annotation performance across spatially relevant NexuST ablations in CosMx liver normal.** Per-cell-type F1 under frozen-encoder linear probing is shown for the matched reduced-size ablation baseline and three variants that directly alter spatial or contextual modelling. The matched baseline retains alternating gene- and cell-level attention. The gene-only encoder replaces cell-level attention with additional gene-level layers while preserving total network depth; the stacked end-to-end encoder applies all gene-level layers before all cell-level layers, removing repeated gene–cell information exchange; and shuffled sampling disrupts spatially coherent patch membership while preserving the selected cells, patch number and size, and per-cell exposure. Points denote means over three random seeds, and horizontal segments span the minimum to maximum seed-averaged F1 across the four configurations within each cell type; error bars are omitted for visual clarity. Cell types are arranged by biological group. The largest reductions relative to the matched baseline occur in several zonated hepatocyte subtypes, particularly Hep 5, whereas immune populations show mixed responses, indicating that the benefit of spatially coherent, repeated gene–cell interaction is cell-state dependent rather than uniform across cell types.

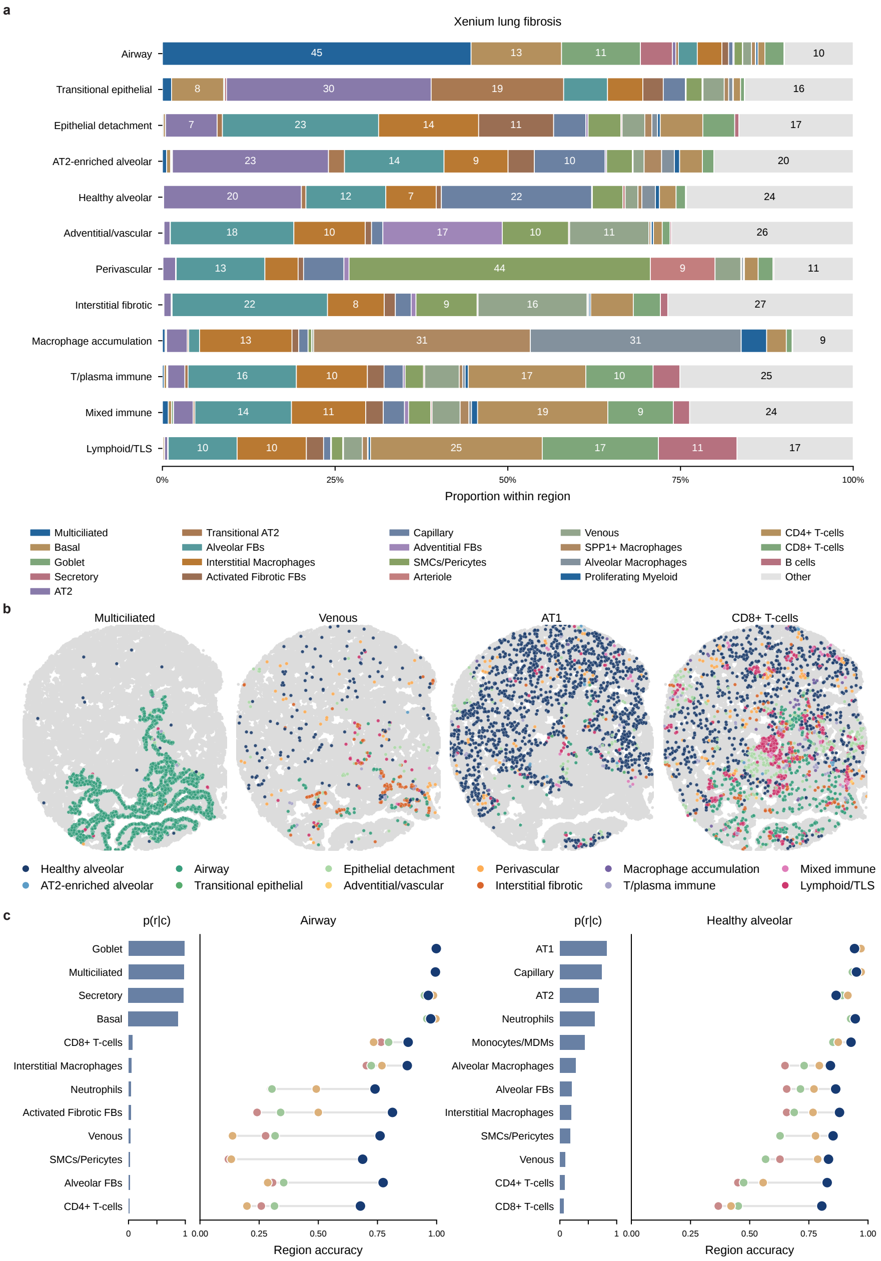


**Fig. 14**: **Cell-type composition and region specificity in Xenium lung fibrosis.**

**Fig. 14**: **a** Cell-type composition of each region, shown as the proportion of each cell type within a region. Stromal and myeloid lineages recur across the vascular and fibrotic regions (Adventitial/vascular, Perivascular, Interstitial fibrotic), and lymphoid and myeloid cells recur across the immune regions, so most regions are not defined by a unique resident cell type. This is the full version of the lineage-level summary in Fig. 3d. **b** Spatial distribution of four representative cell types, with each cell coloured by its region. Multiciliated cells trace airway structures and are almost exclusively assigned to the Airway region; AT1 cells, though spread throughout the parenchyma, are predominantly assigned to Healthy alveolar tissue (dark blue). In contrast, venous endothelial and $CD8^{+}$ T cells are scattered across many regions, with the latter further enriched in lymphoid/TLS foci, so their region cannot be read from identity alone. **c** For Airway and Healthy alveolar, region specificity for each cell type is measured as $p(r \mid c)$, the fraction of that cell type assigned to the region (left bars), alongside its per-method region-prediction accuracy (right bars; NexuST in dark navy). Region-specific cell types (high $p(r \mid c)$) are predicted accurately by all methods, whereas for cell types dispersed across regions (low $p(r \mid c)$), NexuST generally achieves the highest accuracy, often by a substantial margin. This is the full version of Fig. 3h.

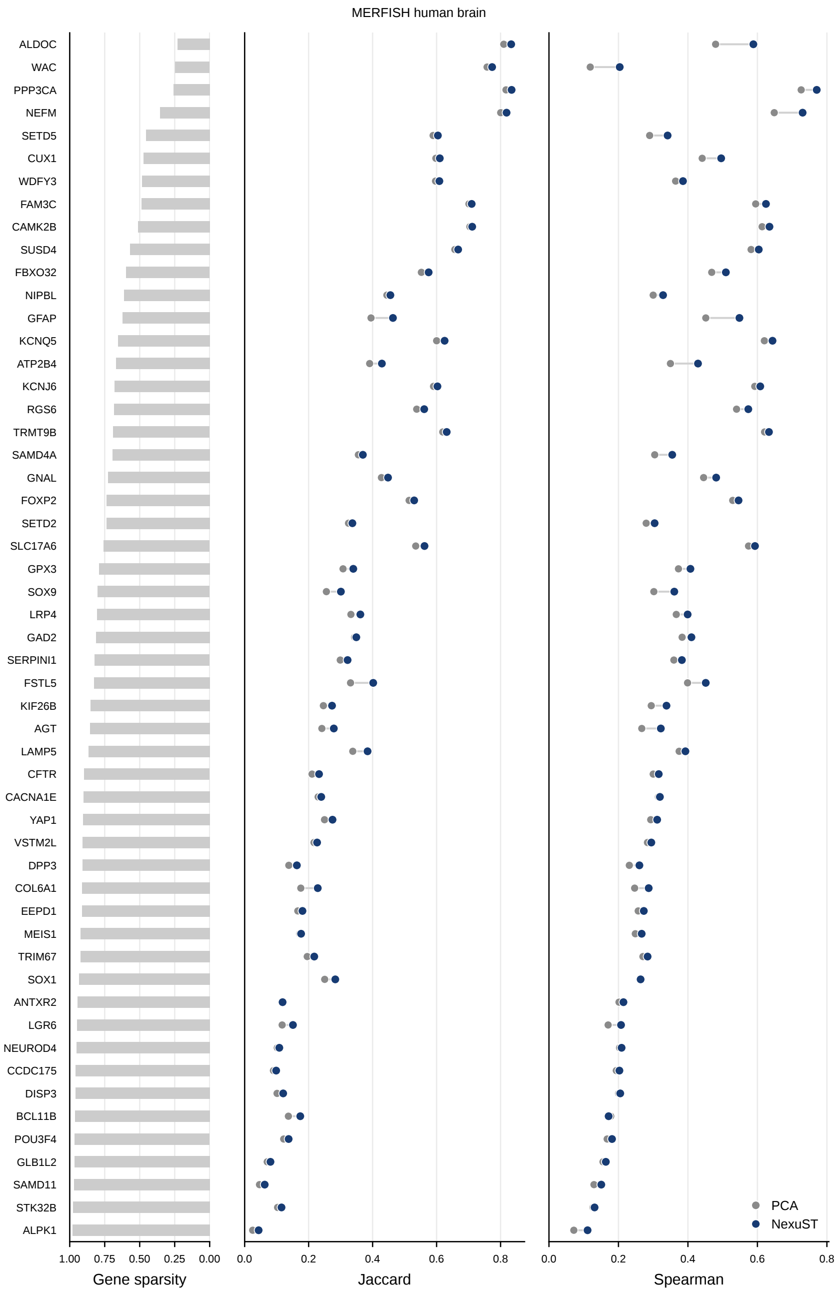


**Fig. 15**: **Gene-level held-out recovery performance in the MERFISH human brain dataset.** Gene sparsity is shown alongside Jaccard similarity and Spearman correlation for NexuST and PCA predictions across selected held-out genes.

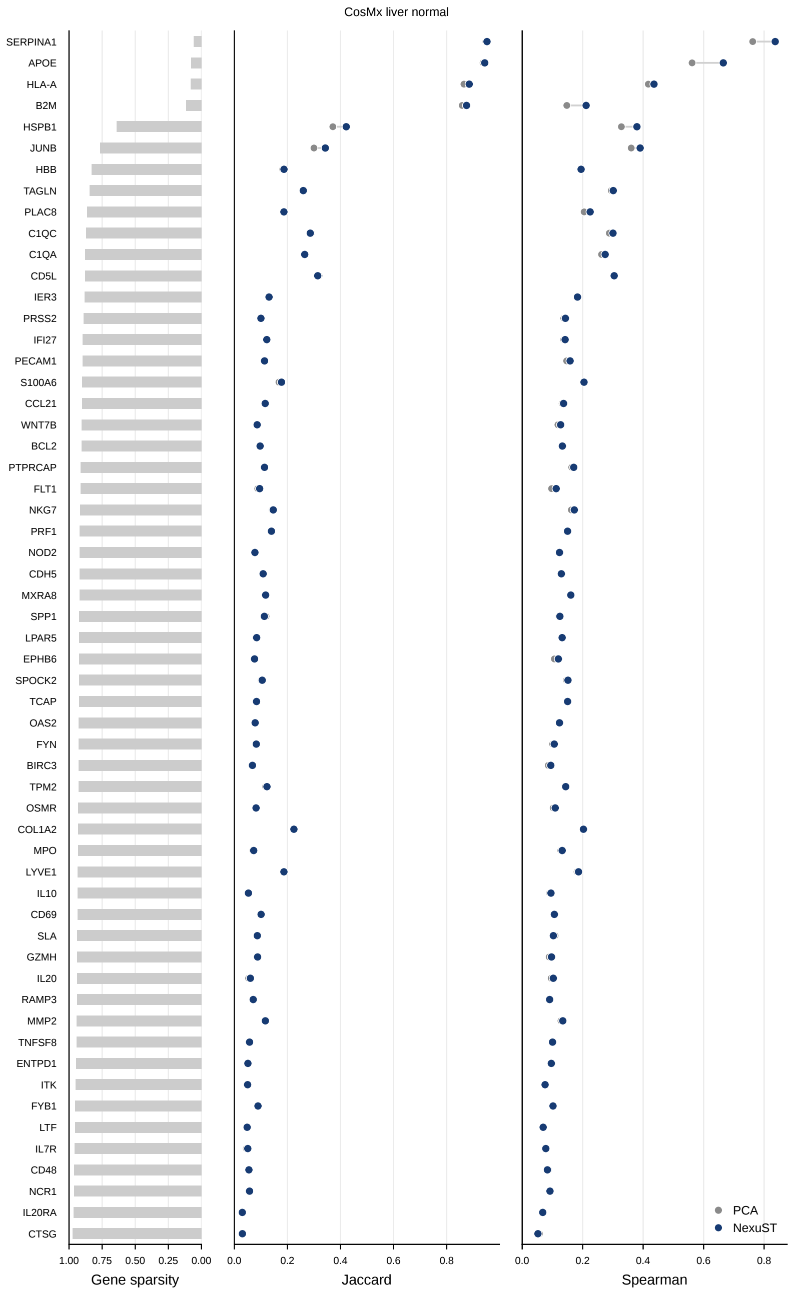


**Fig. 16**: **Gene-level held-out recovery performance in the CosMx liver normal dataset.** Gene sparsity is shown alongside Jaccard similarity and Spearman correlation for NexuST and PCA predictions across selected held-out genes.

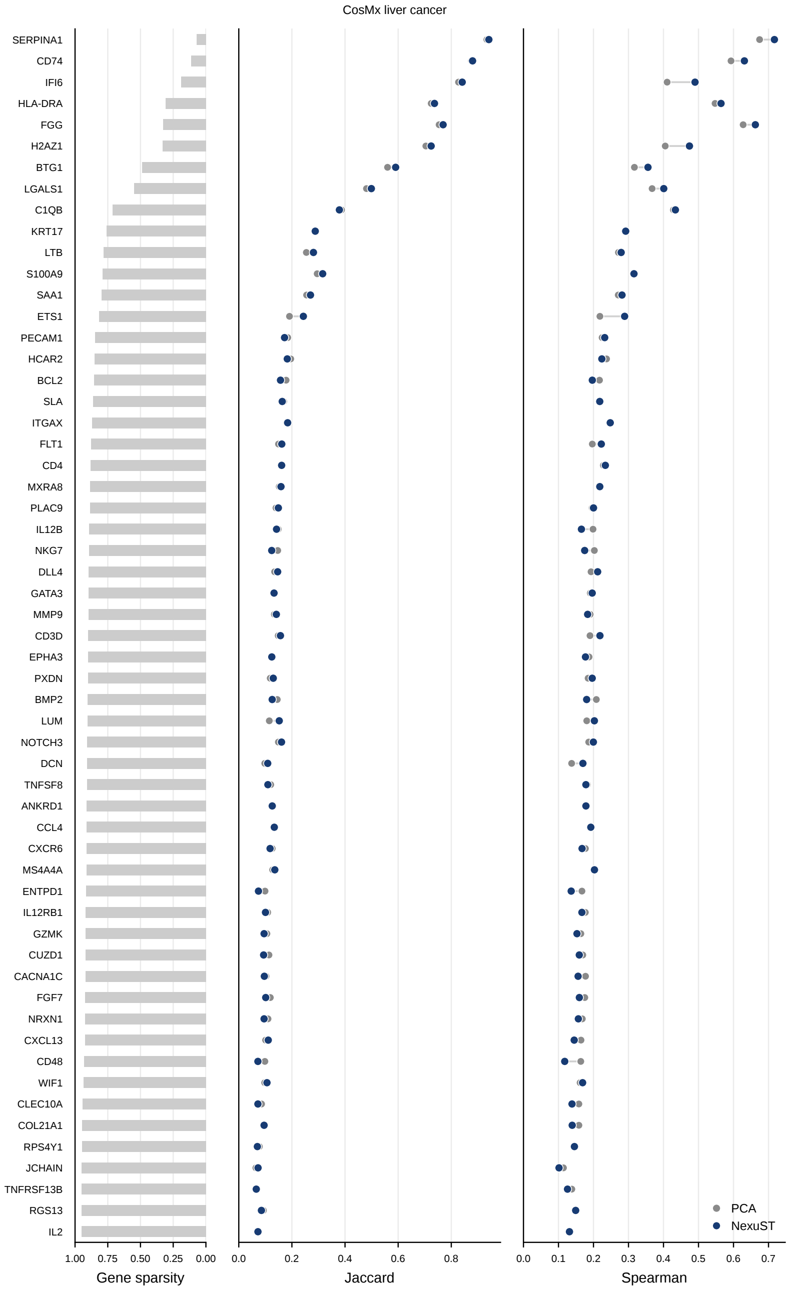


**Fig. 17**: **Gene-level held-out recovery performance in the CosMx liver cancer dataset.** Gene sparsity is shown alongside Jaccard similarity and Spearman correlation for NexuST and PCA predictions across selected held-out genes.

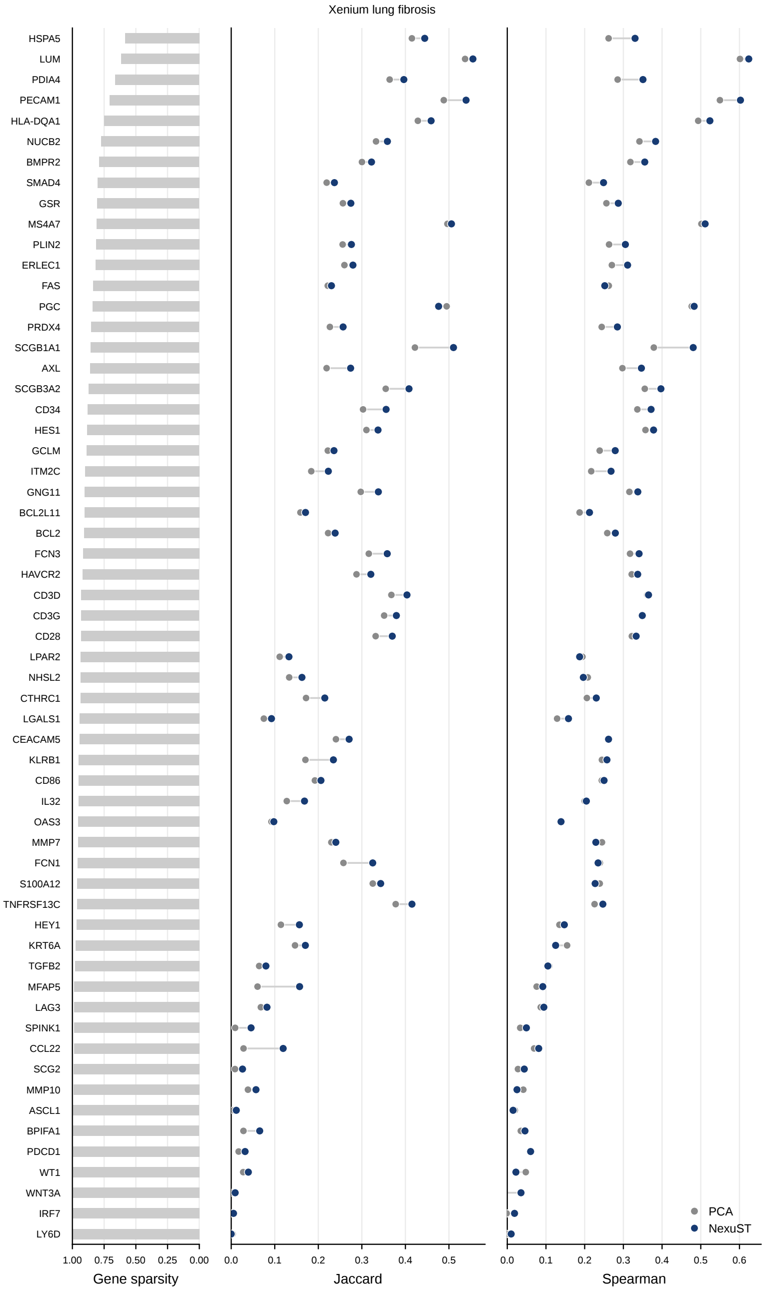


**Fig. 18**: **Gene-level held-out recovery performance in the Xenium lung fibrosis dataset.** Gene sparsity is shown alongside Jaccard similarity and Spearman correlation for NexuST and PCA predictions across selected held-out genes.

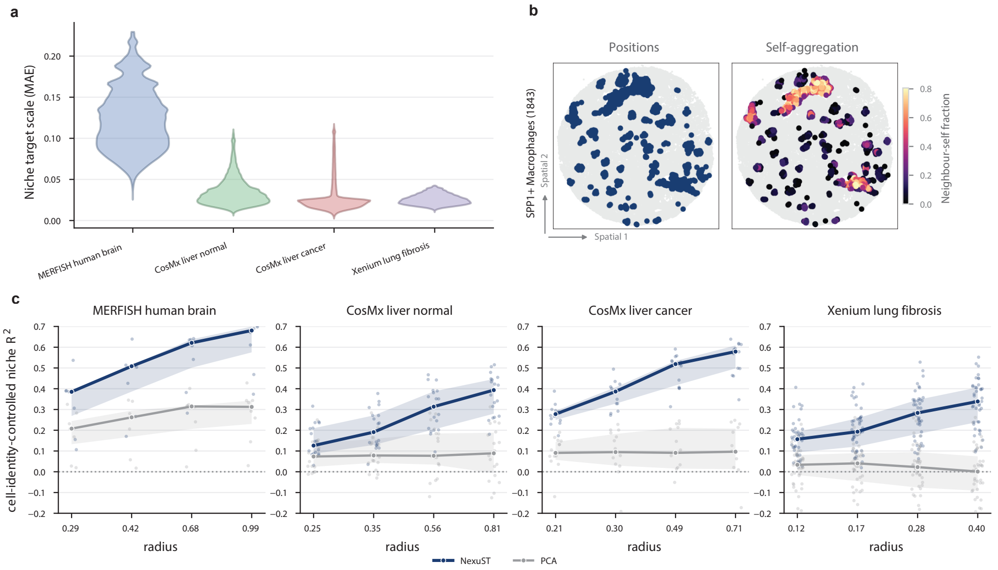


**Fig. 19**: **a. Dataset-specific niche target scale.** Distribution of the per-cell predict-the-mean baseline MAE for each dataset; the intrinsic target scale of niche composition differs several-fold across datasets, so raw MAE should be compared within, not across, datasets. **b. SPP1$^+$ macrophage self-aggregation in Xenium lung fibrosis.** Left: spatial positions of SPP1$^+$ macrophages ($n = 1843$). Right: the same cells coloured by their neighbour-self fraction, the fraction of their neighbours that are also SPP1$^+$ macrophages, showing local self-aggregation. **c. Cell-identity-controlled niche $R^2$ versus radius.** For each dataset, the cell-identity-controlled niche $R^2$ is shown against neighbourhood radius for NexuST (navy) and PCA (grey). Pale points show individual cell-type values, lines show medians across cell types and shaded bands show interquartile ranges. The median for NexuST rises monotonically with radius and remains consistently above PCA, with the gap widening at larger scales.

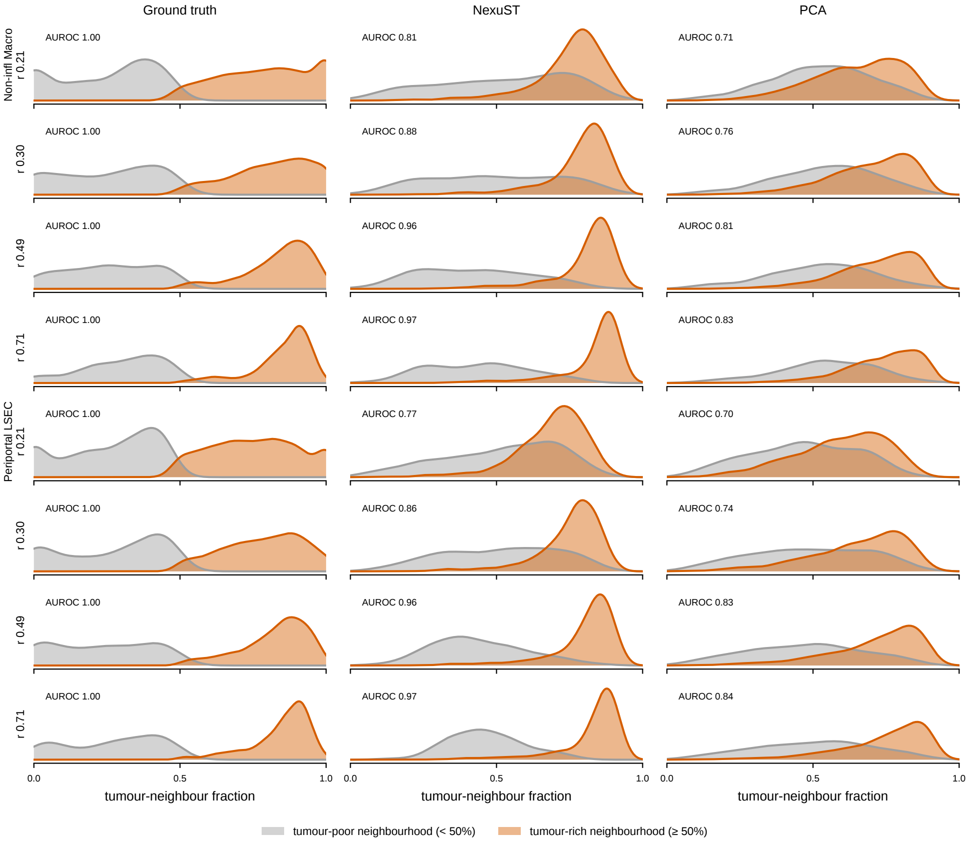


**Fig. 20**: **Tumour-neighbour separability across radii in CosMx liver cancer.** For non-inflammatory macrophages and periportal LSECs, cells are split by their true tumour-neighbour fraction into tumour-poor ($< 50\%$, grey) and tumour-rich ($\geq 50\%$, orange) groups; each panel shows the distribution of the tumour-neighbour fraction for the ground truth and for NexuST and PCA predictions at four radii ($r = 0.21$–$0.71$), with AUROC quantifying how well the predicted score ranks tumour-rich above tumour-poor cells. NexuST separates the two groups far better than PCA at every radius and sharpens with scale (AUROC up to 0.97), whereas PCA remains lower across all radii (AUROC up to 0.84 after rounding).

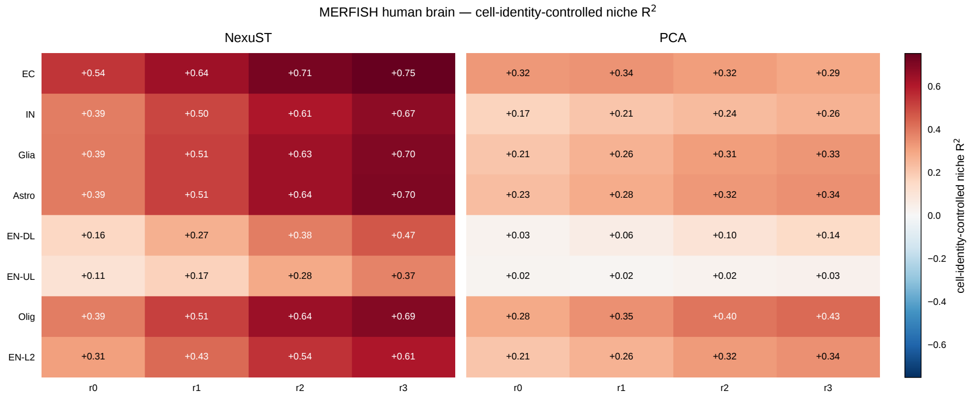


**Fig. 21**: **Cell-identity-controlled niche $R^2$ per cell type and radius in MERFISH human brain.** For each cell type (rows) and neighbourhood radius ($r_0$–$r_3$, columns), the cell-identity-controlled niche $R^2$ is shown for NexuST (left block) and PCA (right block). Both methods remain positive across all cell types and radii, but NexuST exceeds PCA in every case, and the mean gap across cell types widens with increasing radius.

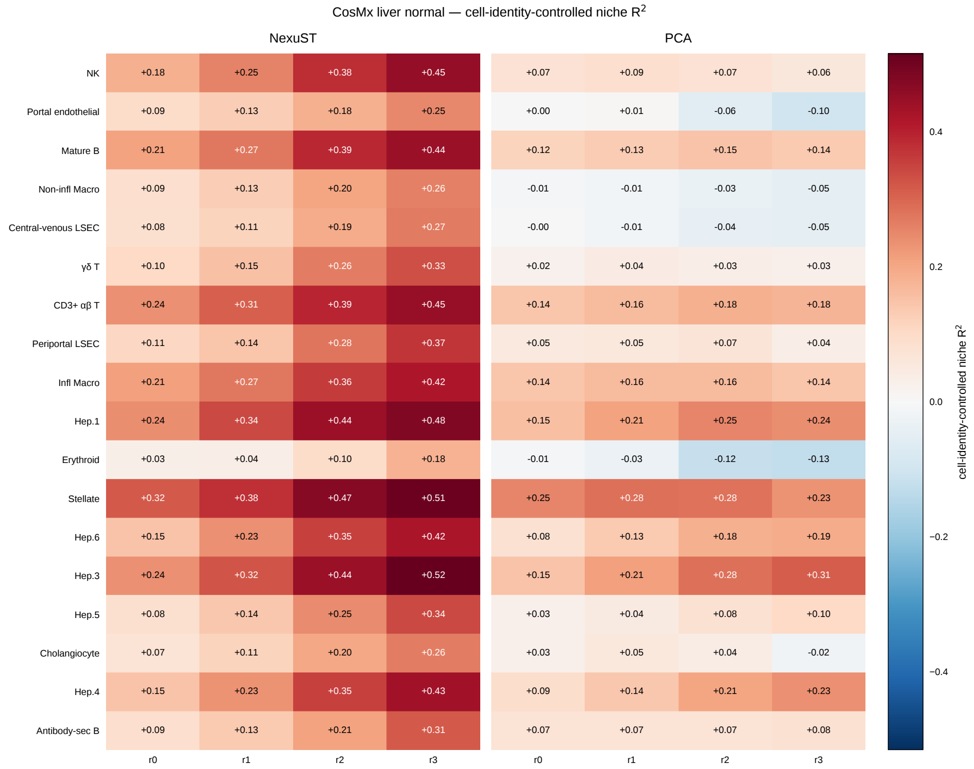


**Fig. 22**: **Cell-identity-controlled niche $R^2$ per cell type and radius in CosMx liver normal.** For each cell type (rows) and neighbourhood radius ($r_0$–$r_3$, columns), the cell-identity-controlled niche $R^2$ is shown for NexuST (left block) and PCA (right block). NexuST remains positive and exceeds PCA for every cell type and radius; PCA is consistently lower and becomes negative for a subset of cell types at one or more radii.

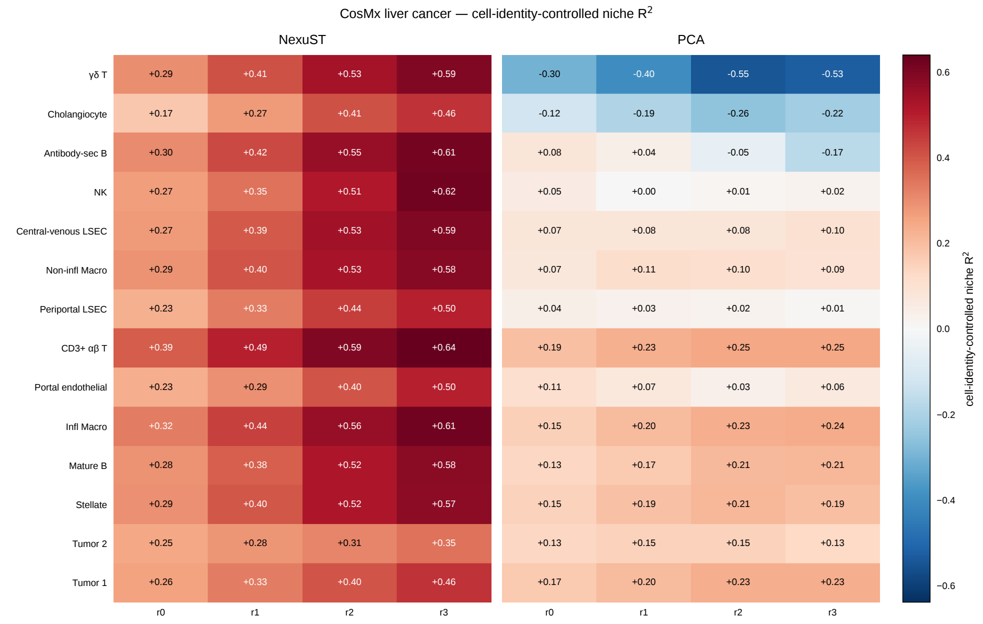


**Fig. 23**: **Cell-identity-controlled niche $R^2$ per cell type and radius in CosMx liver cancer.** For each cell type (rows) and neighbourhood radius ($r_0$–$r_3$, columns), the cell-identity-controlled niche $R^2$ is shown for NexuST (left block) and PCA (right block). NexuST remains positive and exceeds PCA for every cell type and radius; PCA becomes negative for a subset of cell types at one or more radii.

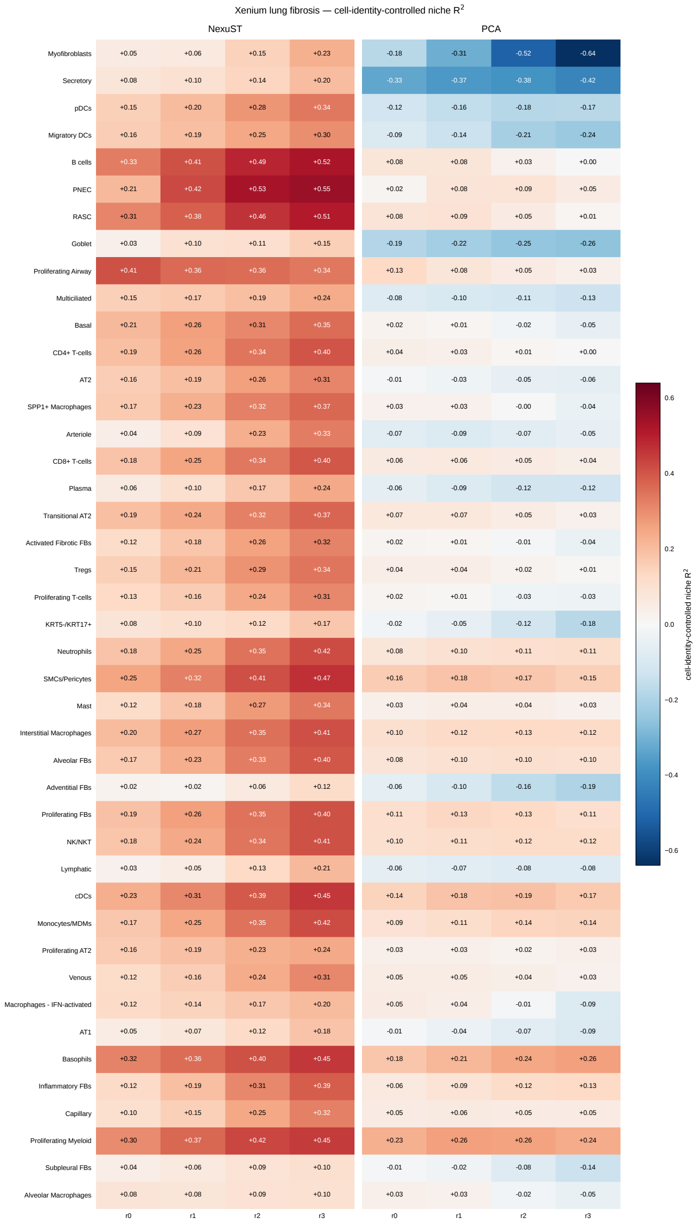


**Fig. 24**: **Cell-identity-controlled niche $R^2$ per cell type and radius in Xenium lung fibrosis.** For each cell type (rows) and neighbourhood radius ($r_0$–$r_3$, columns), the cell-identity-controlled niche $R^2$ is shown for NexuST (left block) and PCA (right block). NexuST remains positive and exceeds PCA for every cell type and radius, whereas PCA remains close to zero overall and becomes negative for many cell types at one or more radii.

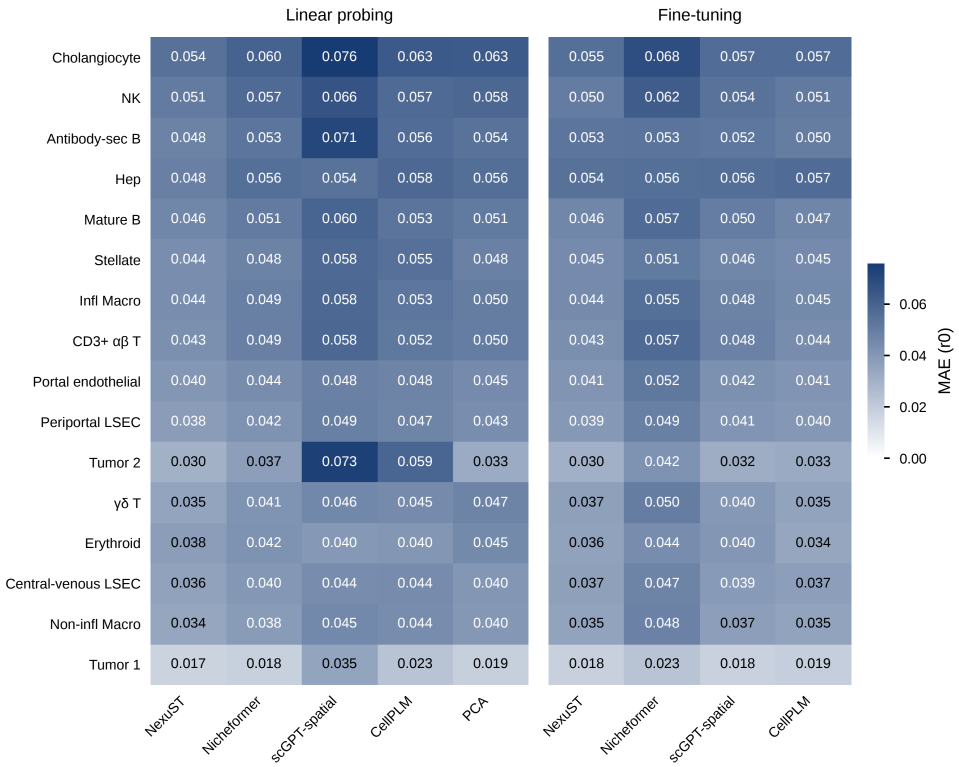


**Fig. 25**: **Raw per-cell-type niche-composition MAE in CosMx liver cancer at radius $r_0$.** Mean absolute error per cell type (rows) for each method (columns) under linear probing (left block) and fine-tuning (right block). Even on the raw MAE scale, NexuST attains the lowest error for every cell type under linear probing.

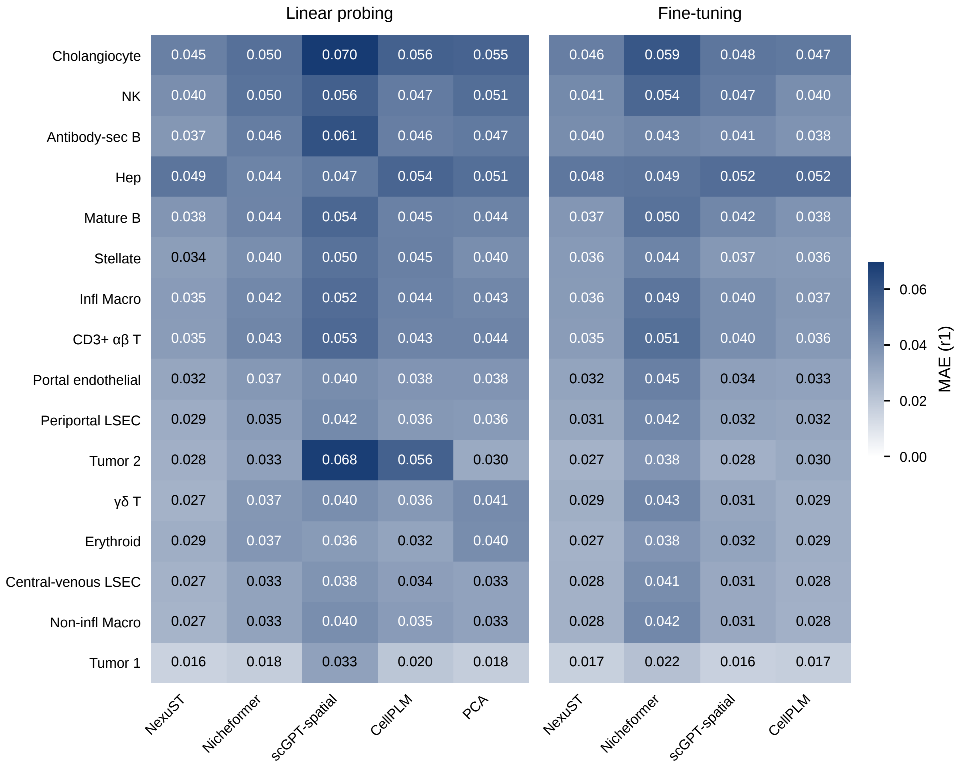


**Fig. 26**: **Raw per-cell-type niche-composition MAE in CosMx liver cancer at radius $r_1$.** Mean absolute error per cell type (rows) for each method (columns) under linear probing (left block) and fine-tuning (right block). Even on the raw MAE scale, NexuST attains the lowest error for every cell type except hepatocytes under linear probing.

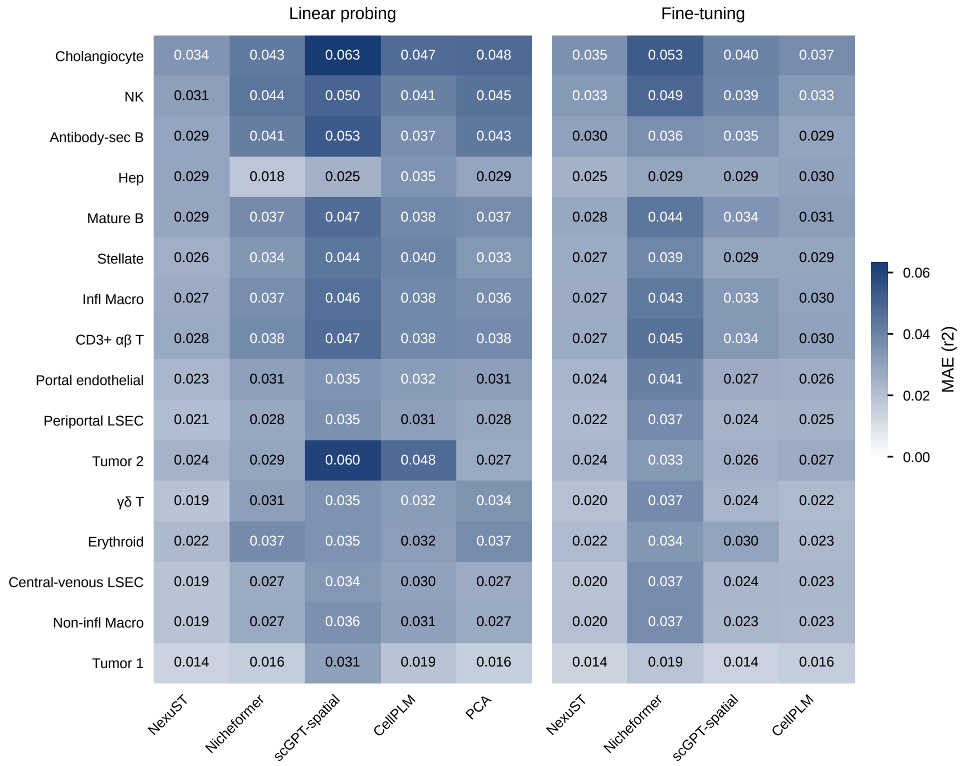


**Fig. 27**: **Raw per-cell-type niche-composition MAE in CosMx liver cancer at radius $r_2$.** Mean absolute error per cell type (rows) for each method (columns) under linear probing (left block) and fine-tuning (right block). Even on the raw MAE scale, NexuST attains the lowest error for every cell type except hepatocytes under linear probing.

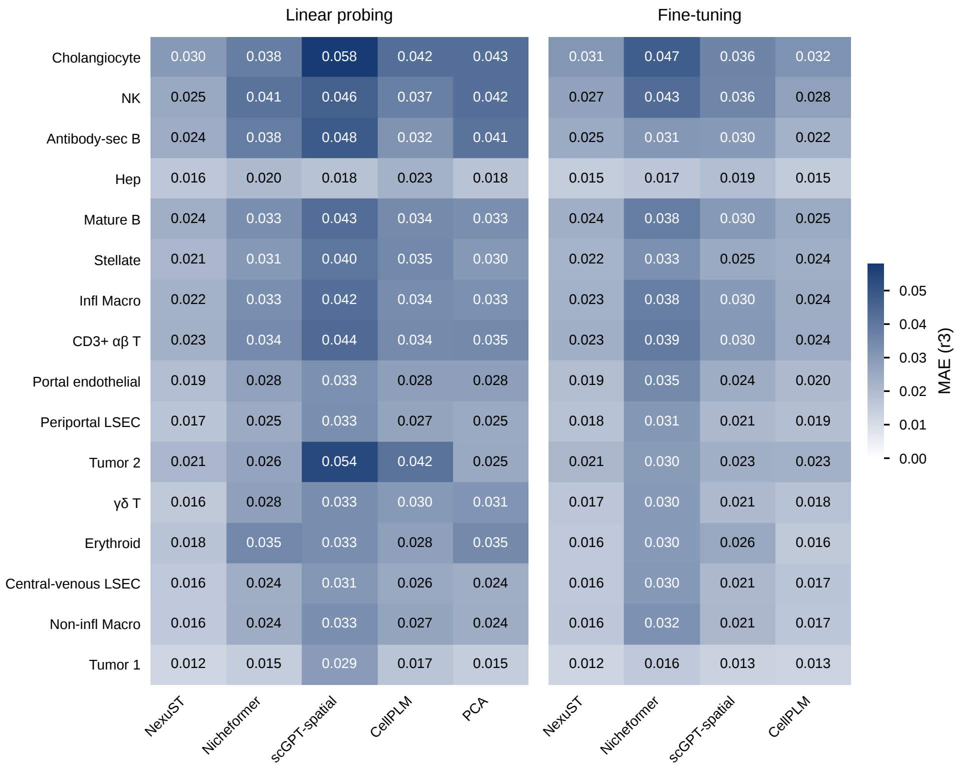


**Fig. 28**: **Raw per-cell-type niche-composition MAE in CosMx liver cancer at radius $r_3$.** Mean absolute error per cell type (rows) for each method (columns) under linear probing (left block) and fine-tuning (right block). Even on the raw MAE scale, NexuST attains the lowest error for every cell type under linear probing.

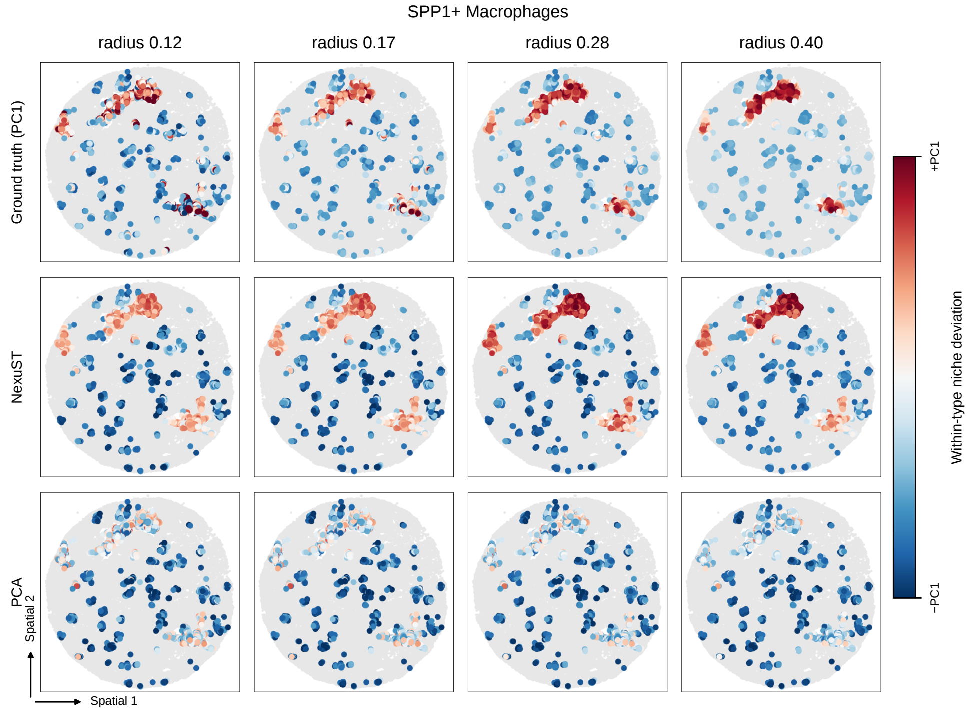


**Fig. 29**: **Within-type niche deviation of SPP1$^{+}$ macrophages across radii in Xenium lung fibrosis.** Spatial maps of within-type niche deviation (first principal component, ±PC1) at four neighbourhood radii (0.12–0.40), comparing the ground truth with NexuST and PCA predictions (rows). NexuST tracks the growing self-aggregates across scales more closely than PCA.

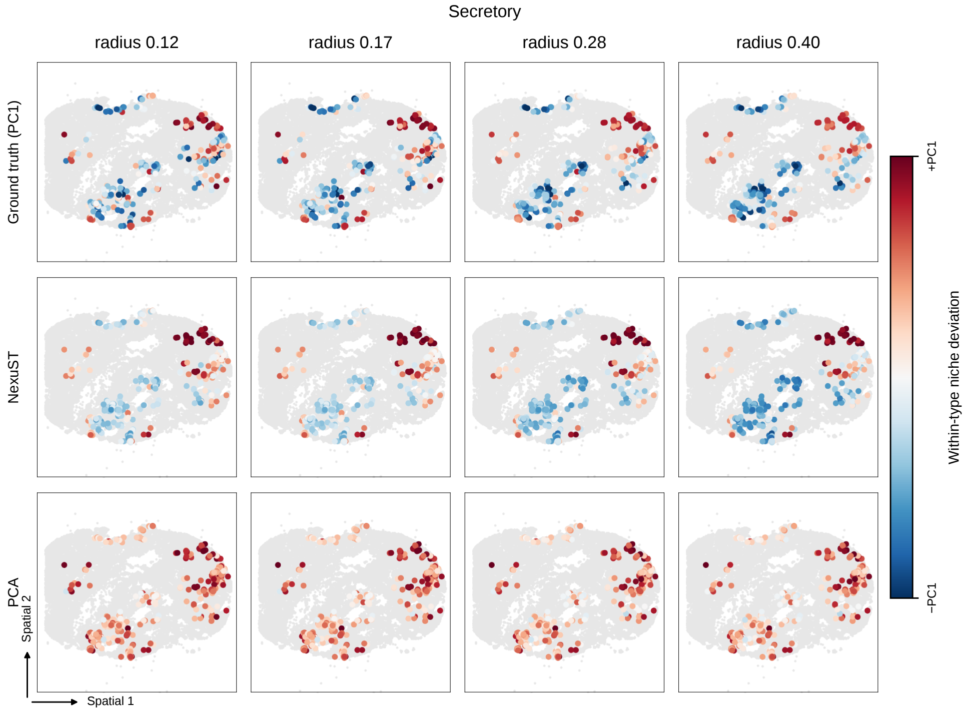


**Fig. 30**: **Within-type niche deviation of Secretory cells across radii in Xenium lung fibrosis.** Spatial maps of within-type niche deviation (first principal component, ±PC1) at four neighbourhood radii (0.12–0.40), comparing the ground truth with NexuST and PCA predictions (rows). NexuST more closely reproduces the ground-truth spatial pattern across scales than PCA.

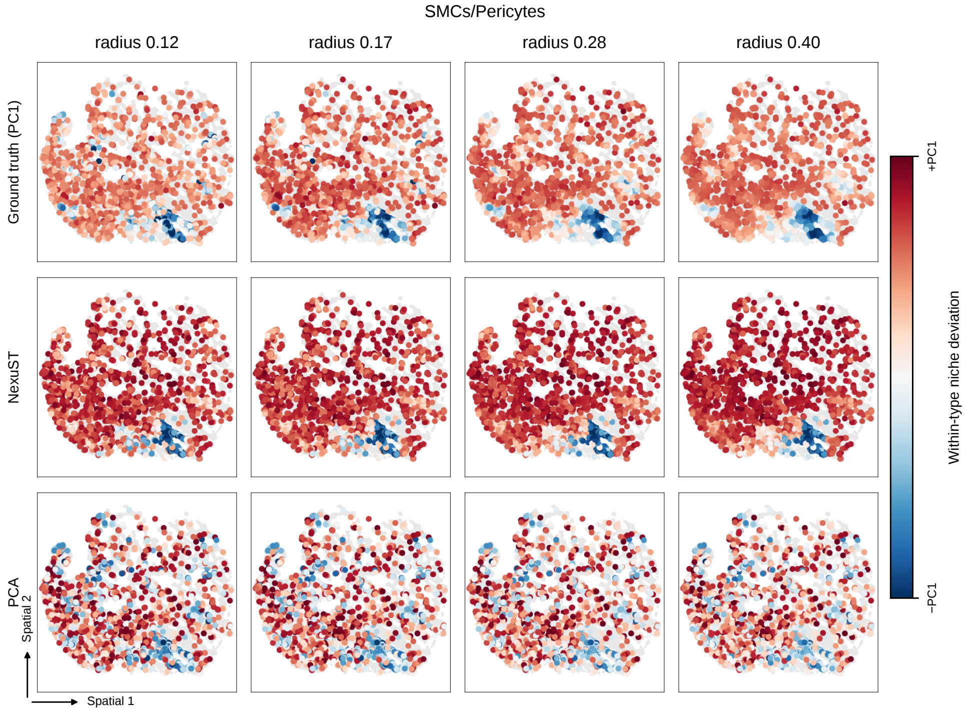


**Fig. 31**: **Within-type niche deviation of smooth muscle cells/pericytes across radii in Xenium lung fibrosis.** Spatial maps of within-type niche deviation (first principal component, ±PC1) at four neighbourhood radii (0.12–0.40), comparing the ground truth with NexuST and PCA predictions (rows). NexuST more closely reproduces the ground-truth spatial pattern across scales than PCA.

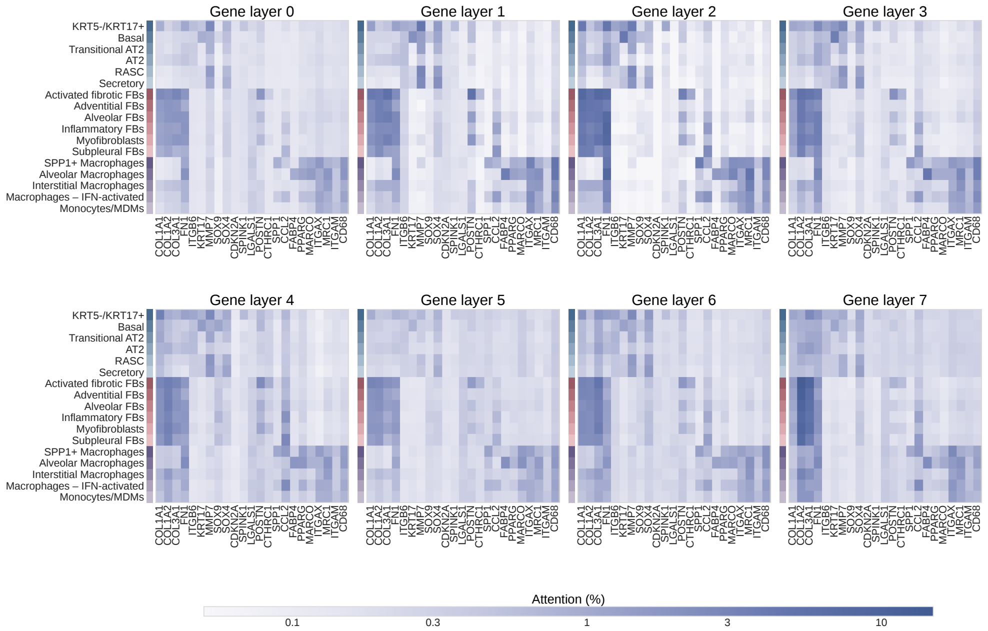


**Fig. 32**: **Pretrained cell-to-gene attention across all gene layers in Xenium lung fibrosis.** Mean attention from the cell token to selected gene tokens is shown for 17 representative cell types (rows) and 23 selected genes (columns) across gene layers L0–L7. Attention was pooled across the five fixed validation FOVs and is reported as a percentage using a common logarithmic colour scale across all layers.

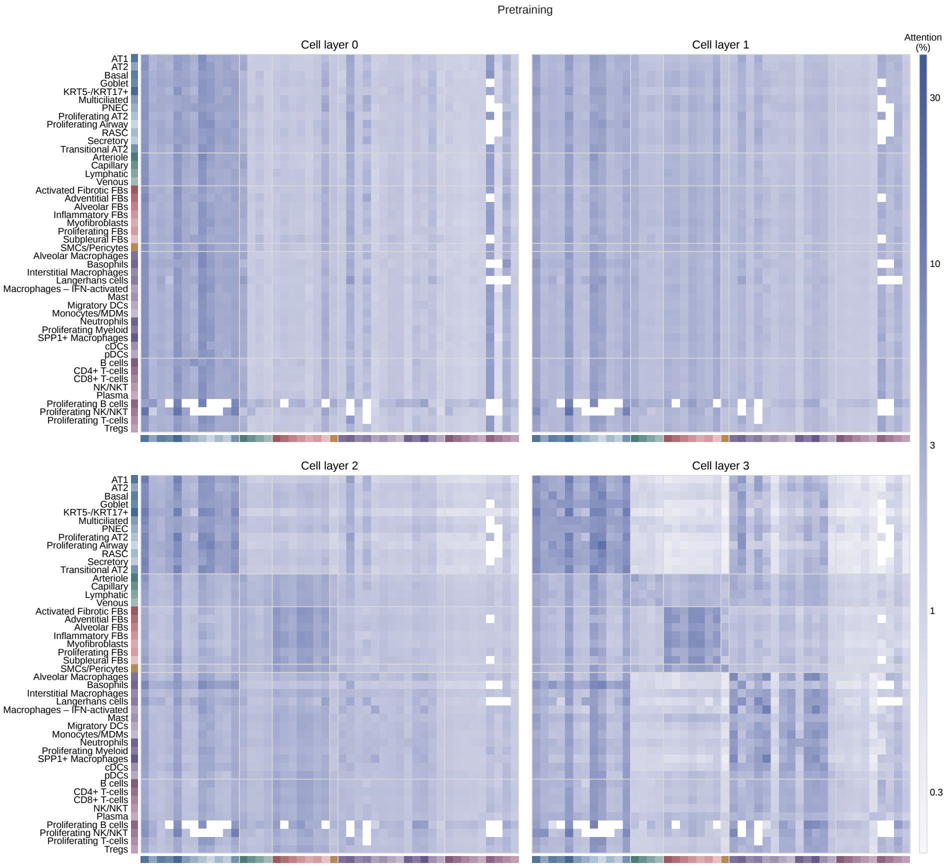


**Fig. 33**: **Pretrained cell-to-cell attention across all cell layers in Xenium lung fibrosis.** Abundance-normalised cell-to-cell attention among all 46 cell types observed in the five fixed validation FOVs is shown for cell layers L0–L3 on this page and L4–L7 on the continued page. Rows denote source cell types and columns denote target cell types, with cell types ordered into six lineages indicated by the colour strips and block boundaries. Exact self-cell pairs were excluded before abundance normalisation, so diagonal entries represent attention to other cells of the same annotated type. Colour indicates the percentage of each source cell type's normalised attention assigned to each target cell type on a common logarithmic scale. White matrix entries indicate source–target cell-type pairs for which no eligible within-chunk cell pairs were available across the selected FOVs.

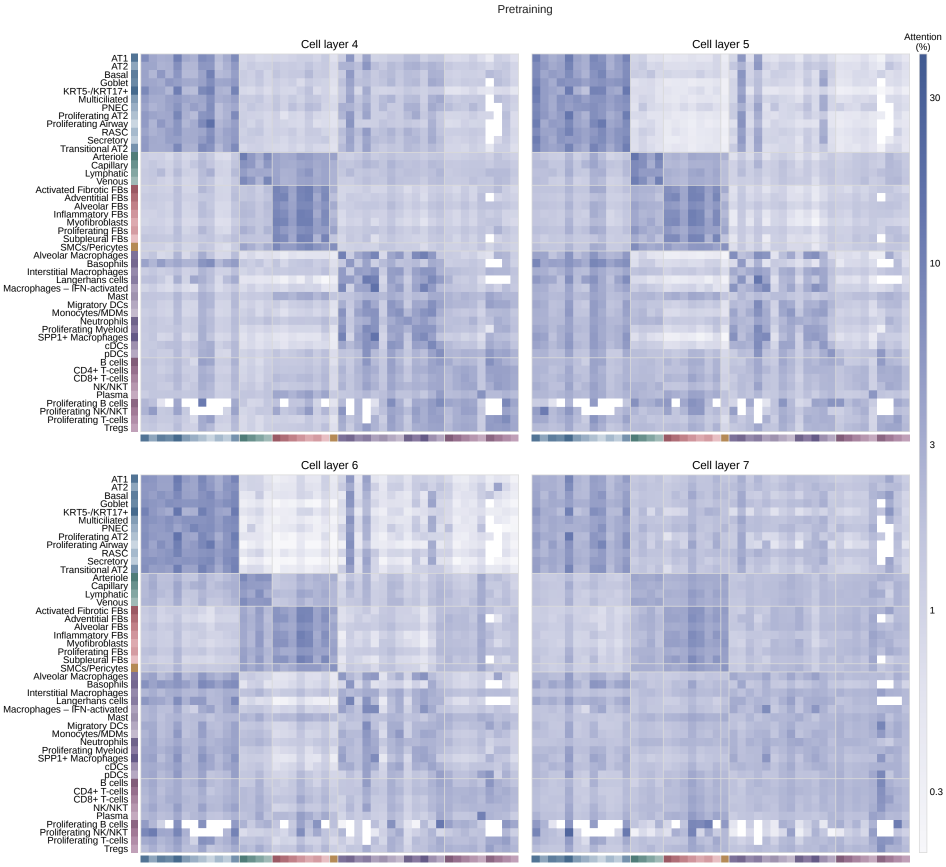


**Fig. 33**: (continued)
